# Calculating metalation in cells reveals CobW acquires Co^II^ for vitamin B_12_ biosynthesis upon binding nucleotide

**DOI:** 10.1101/2020.06.26.173062

**Authors:** Tessa R. Young, Maria Alessandra Martini, Deenah Osman, Richard J. Morton, Evelyne Deery, Martin J. Warren, Nigel J. Robinson

**Affiliations:** Department of Biosciences, Durham University, Durham, UK; Department of Chemistry, Durham University, Durham, UK; Max Planck Institute for Chemical Energy Conversion, Mülheim an der Ruhr, Germany; Department of Mathematics, Physics, and Electrical Engineering, Northumbria University, Newcastle-upon-Tyne, UK; School of Biosciences, University of Kent, Canterbury, Kent, UK; Quadram Institute Bioscience, Norwich Research Park, Norfolk, UK

## Abstract

Protein metal-occupancy (metalation) *in vivo* has been elusive. Here we develop a metalation-calculator which accounts for inter-metal competition and changing metal-availabilities inside cells. The calculations are based on available free-energies of metals determined from the responses of metal sensors. We use the calculator to understand the function and mechanism of CobW, a predicted Co^II^-chaperone for vitamin B_12_. CobW is calculated to acquire negligible metal alone: But, upon binding nucleotide (GTP) and Mg^II^, CobW assembles a high-affinity site that can obtain Co^II^ or Zn^II^ from the intracellular milieu. In idealised cells with sensors at the mid-points of their responses, competition within the cytosol enables Co^II^ to outcompete Zn^II^ for binding CobW. Thus, Co^II^ is the cognate metal. However, after growth in different [Co^II^], Co^II^-occupancy ranges from 10 to 97% which matches CobW-dependent B_12_ synthesis. The calculator reveals how CobW acquires its metal and is made available for use with other proteins.

---

Paradoxically, *in vitro*, most metalloproteins prefer to bind incorrect metals^1,2^. A non-cognate metal may bind more tightly to the native site or bind by using a subset of the native ligands, by recruiting additional ligand(s) and/or by distorting the geometry of a binding site. Some enzymes are cambialistic and can function with alternative metals^3^, but more commonly a non-cognate metal will inactivate an enzyme^4,5^. Correct metalation occurs *in vivo* because cells carefully control the availability of metals to nascent proteins^1,6–8^. For example, specialised delivery proteins support metal acquisition by about a third of metalloproteins, (which in turn represent about a third of all proteins and about a half of all enzymes)^1,8^. A substantial fraction (>80%) of these delivery systems initially supply metal to cofactors such as heme, chlorophyll, iron sulphur clusters and vitamin B_12_^1^. Subsequent acquisition of a preassembled cofactor is then less of a challenge since a binding pocket is more readily selective for a complex molecule as opposed to a single metal atom. However, metal delivery proteins do not ultimately solve the challenge of metalation because now the correct metal must somehow partition onto the delivery protein. Here we discover how the correct metal is acquired by a metal delivery protein.

The G3E GTPase superfamily contains three branches of delivery proteins involved in the assembly of metal centres, two for Ni^II^ (HypB, UreG), one for handling the cobalamin cofactor (MeaB), plus a fourth family, COG0523^9,10^. Though ubiquitous, from bacteria to plants and humans, members of COG0523 have been persistently enigmatic^10^. Gene context and informatics have linked subgroups of this family to at least three different metals: These include Nha3 associated with Fe^III^-requiring nitrile hydratases^11–13^, various subgroups (including YeiR, YjiA, ZigA and ZagA) implicated in Zn^II^ metallostasis^10,14–18^, and CobW associated with the aerobic biosynthesis of cobalamin (vitamin B_12_) and hence Co^II^ (ref.^19^). Metal insertion into the preformed corrin ring in the aerobic pathway for vitamin B_12_biosynthesis appears to be irreversible^20,21^, highlighting the importance of Co^II^ specificity at this step. Disruption of *cobW* impairs B_12_ biosynthesis^19^, and a role in Co^II^ delivery has been suggested^22^, but not established.

Nucleotide hydrolysis is critical for the metallochaperone activities of HypB^23^, UreG^24^ and MeaB^25^, and evidence is emerging that this is also the case for other COG0523 proteins. Recently, the putative Zn^II^ chaperone ZagA was shown to interact with a Zn^II^-requiring enzyme of folate biosynthesis (FolE)^18^. Rather than the anticipated GTP, this interaction is stimulated by the purine intermediate ZTP (5-amino 4-imidazole carboxamide riboside 5’-triphosphate), an alarmone that accumulates during low Zn^II^ (ref.^18,26^). A Zn^II^-requiring histidine lyase (HutH) together with ZigA enables depletion of histidine in cells cultured in low Zn^II^ and this may serve to liberate histidine-bound Zn^II^ (ref.^16^). Zn^II^ binding to ZigA enhances GTP hydrolysis and weakens GDP binding^17^. The impact of triphosphonucleotide binding on metal binding by COG0523 proteins remains to be tested.

For metalloproteins generally, there is a need to relate metal binding to the intracellular availability of metals. Our recent work provides the basis for such contextualisation^27^. Cells are thought to assist protein metalation by maintaining availabilities to the opposite of the Irving-Williams series with weaker binding metals such as Mg^II^, Mn^II^ and Fe^II^ highly available and tighter binding metals such as Ni^II^, Zn^II^ and Cu^I^ at low availabilities^28–30^. We have demonstrated this to be correct by determining the sensitivities of the DNA-binding metal-sensing transcriptional regulators of *Salmonella enterica* serovar Typhimurium (hereafter *Salmonella*)^27^. The sensors trigger expression of genes whose products, for example, import metals that are deficient or export those in excess^6,31^. A collection of thermodynamic parameters were measured for each sensor and used to calculate the (dynamic range of) buffered intracellular metal concentrations to which each sensor is finely tuned to switch gene expression^27,32^. For the more competitive metals, detection is so sensitive as to suggest that there is no hydrated metal in the cell^27,28^. Instead, rapid associative metal-exchange can occur between labile ligands in the crowded cytosol and the binding sites of metalloproteins, making it unhelpful to express metal availabilities as concentrations of the (largely irrelevant and negligible) hydrated species: Thus, the chemical potentials of the bound available metals were expressed as free energies Δ*G*^27^. It is hypothesised that metal-delivery proteins acquire their metals from these exchangeable, buffered pools. By reference to available Δ*G* values and by assuming an idealised cell in which the sensors are at the mid-points of their dynamic ranges, the correct metal (Co^II^) was previously predicted to partition to the known chelatase of the anaerobic cobalamin biosynthetic pathway, CbiK^27^. Here, we build upon this approach to account for (1) multiple competing metals and (2) non-idealised (conditional) cell cultures, in order to understand the actions of the putative metal delivery protein CobW. With so many enzymes requiring metals, an ability to calculate and optimise *in vivo* metalation has far-reaching applications, for example in industrial biotechnology.

Vitamin B_12_ is an essential nutrient that is neither made nor required by plants^33^. Prokaryotes present in the ruminant microbiome produce B_12_ and hence dairy products provide a dietary source^34^. Vitamin B_12_ supplements are recommended for those on a vegan diet and its biomanufacture (the only feasible production method for such a complex molecule) is increasingly in demand^35^. *E. coli* has significant advantages (namely, it is fast-growing and genetically tractable) over currently employed production strains^36^. Native *E. coli* does not make vitamin B_12_ but strains containing functional B_12_ pathways have been created, initially utilising genes of the anaerobic pathway from *Salmonella^37^* and more recently using those of the aerobic pathway primarily from *Rhodobacter capsulatus*^38–40^. The latter has enabled the production of previously difficult to isolate intermediates, including the metal-free corrinoids hydrogenobyrinic acid and its diamide^38–40^. In *R. capsulatus* Co^II^ is inserted into the corrin ring of hydrogenobyrinic acid *a,c*-diamide by a cobalt chelatase ATPase (CobNST)^41^, putatively via CobW^22^. However, a better understanding of Co^II^-availability inside engineered *E. coli* strains (referred to hereafter as *E. coli\**) is required in order to optimise Co^II^ supply for the B_12_ pathway within the heterologous host.

A purpose of this work was to determine whether CobW can acquire Co^II^ and supply the metal to the aerobic B_12_ biosynthetic pathway. *E. coli\** strains have been used as the model because this has direct relevance to biomanufacturing, but also because high B_12_ production in these cells coupled with the close similarity between the DNA-binding metal sensors of *E. coli* and *Salmonella* both serve to make this system experimentally tractable: The metal sensors of *Salmonella* having been thermodynamically characterised^27^. Here we determine the metal affinities of CobW and discover that a high-affinity metal-binding site is assembled only upon association with Mg^II^ and GTP. We calculate the metal-occupancy of CobW *in vivo* using metal-availabilities in an idealised cell determined from the sensitivities of metal sensors. This establishes Co^II^ as the cognate metal, despite CobW also having a tight (sub-picomolar) Zn^II^-affinity. By calculating the Co^II^ availabilities in *E coli*\* from the response of the Co^II^-sensor RcnR, we show that Mg^II^GTP-CobW can be mis-metalated by Zn^II^ *in vivo*, but this is precluded when Co^II^ availability increases. These predictions are reflected in the CobW-dependent production of vitamin B_12_ in *E. coli*\*, establishing a role for CobW in Co^II^-supply for B_12_. Together, these data reveal a mechanism for Co^II^-acquisition and Co^II^-supply by CobW, with significance for understanding the actions of other COG0523 proteins. These data will also allow optimisation of B_12_ manufacture in *E. coli\** strains.

An easy-to-use metalation calculator has been developed which accounts for competition between metals at a protein metal-binding site, for competition from the intracellular milieu, and for variable metal availabilities in bacterial cells. The calculator can be readily applied by others to a diversity of metalloproteins across bioscience and biotechnology.

## Results

### Guanine nucleotides create two metal-sites in CobW

The first objective was to measure the Co^II^ affinities of the form of CobW that acquires metal inside a cell. A modelled structure of CobW (Fig. 1a) showed hypothetical nucleotide-binding sequences adjacent to a putative metal-binding motif, CxCC, and both of these features are conserved in the COG0523 subfamily^9,10^. To assess the effect of nucleotides on metal-binding, CobW was overexpressed and purified (Fig. 1b and Supplementary Fig. 1). The protein mass determined by ESI-MS (37,071 Da; Fig. 1c) is consistent with that expected for CobW after cleavage of the N-terminal methionine (37,072.6 Da).

**Fig 1.**
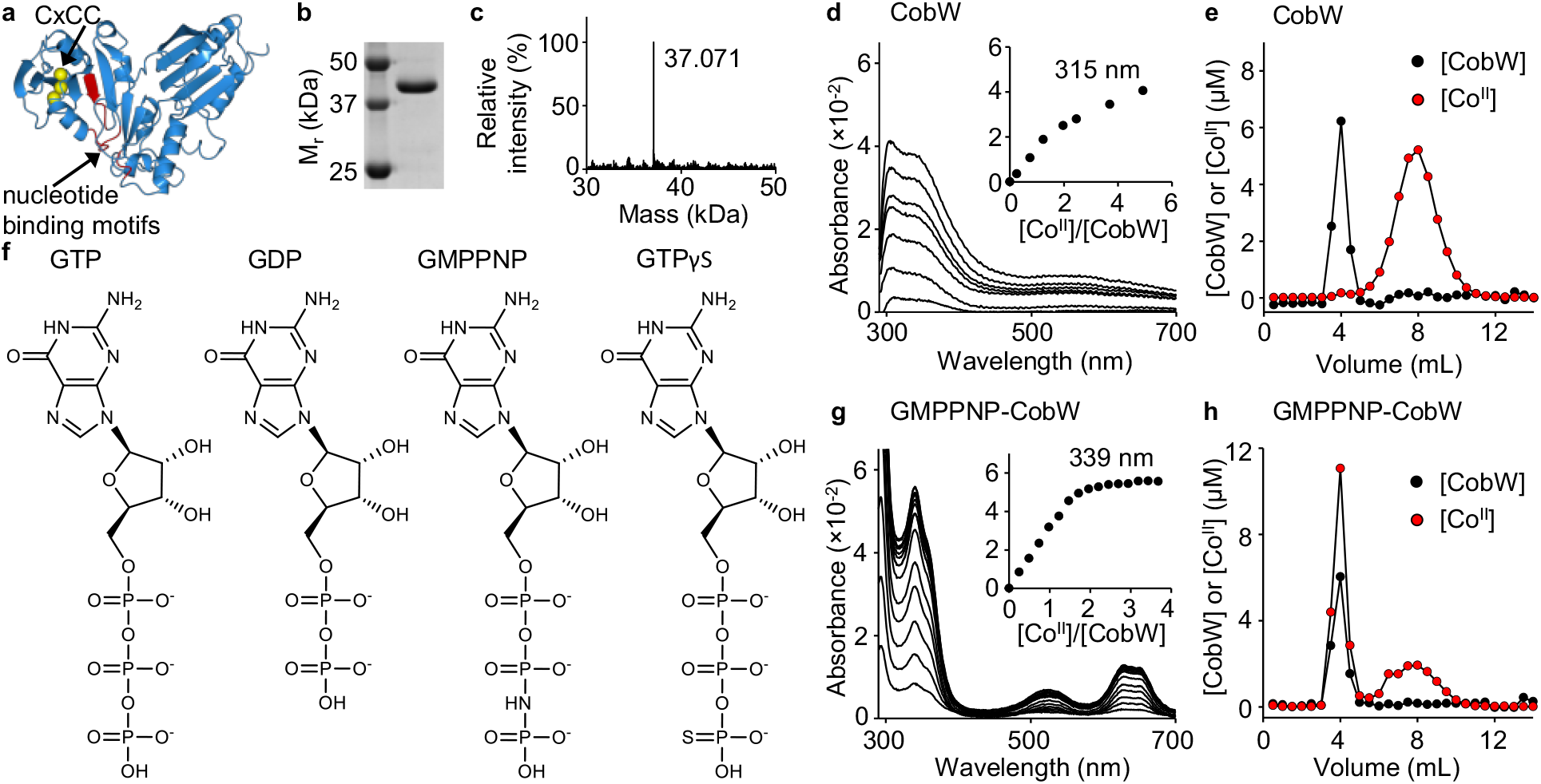
Co^II^ binding to CobW is enhanced by guanine nucleotides. **a** Homology model of CobW (generated with SWISS-MODEL^69^ using *E. coli* YjiA PDB entry 1NIJ^70^ as template) showing sulphur atoms from conserved CxCC motif (in yellow) and nucleotide-binding (GxxGxGK, hhhExxG, SKxD*) motifs^9,10^ (in red). *Ordinarily NKxD but [ST]KxD observed in some COG0523 proteins^9^. **b** Purified CobW analysed by SDS-PAGE (full image in Supplementary Fig. 1). **c** ESI-MS analysis (de-convoluted spectra) of purified CobW. **d** Apo-subtracted spectra of Co^II^-titrated CobW (26.1 μM); feature at 315 nm (inset) shows a non-linear increase. **e** Representative (n=2) elution profile following gel-filtration of a mixture of CobW (10 μM) and Co^II^ (30 μM) showing no co-migration of metal with protein. Fractions were analysed for protein by Bradford assay and for metal by ICP-MS. **f** Structures of nucleotides used in this work. **g** As in (**d**) for a mixture of CobW (24 μM) and GMPPNP (60 μM); feature at 339 nm (inset) showing a linear increase saturating at 2:1 ratio Co^II^:CobW. **h** As in (**e**) for a mixture of CobW (10 μM), Co^II^ (30 μM) and GMPPNP (30 μM) shows co-migration of 1.8 equivalents Co^II^ per CobW (mean value from peak integration, n=2 independent experiments).

Co^II^-titration of CobW alone (26.1 μM) produced a non-linear increase in absorbance at 315 nm (Fig. 1d) but gel-filtration of a mixture of CobW (10 μM) and Co^II^ (30 μM) resulted in their complete separation (Fig. 1e). Taken together, these results suggest only weak interactions between Co^II^ and CobW in the absence of cofactors. In the presence of excess GMPPNP (60 μM), a less readily hydrolysed analogue of GTP (Fig. 1f), Co^II^-titration of CobW (24 μM) produced an absorbance feature at 339 nm characteristic of ligand-to-metal charge transfer with an extinction coefficient (ε ~ 2,800 cm^−1^ M^−1^) indicative of coordination by three cysteine side-chains^42^ (Fig. 1g). Visible absorbance features (500 – 700 nm, ε ~ 300 – 700 cm^−1^ M^−1^) are characteristic of *d-d* transitions, diagnostic of tetrahedral Co^II^-coordination geometry (Fig. 1g and Supplementary Fig. 2). Equivalent experiments performed with GTP and an alternate stable analogue, GTPγS, generated indistinguishable spectra (Supplementary Fig. 3a,b). These absorbance features increased linearly saturating at 2:1 ratio Co^II^:CobW, and gel-filtration of a mixture of CobW (10 μM) and Co^II^ (30 μM) pre-incubated with GMPPNP (30 μM) resulted in co-migration of ~ 2 equivalents Co^II^ per protein monomer (Fig. 1h). These data show that binding of guanine nucleotides to CobW promotes tight coordination of two metals ions.

### Addition of cellular [Mg^II^] reveals one distinct Co^II^ site

The uniform absorbance increase observed across both metal-binding events in Fig. 1g, could be explained by either the presence of two sequentially filled sites with identical spectroscopic features, or two spectrally distinct sites being filled in a pairwise manner. Competition between GMPPNP-CobW and ethylene glycol tetraacetic acid (EGTA) for Co^II^ produced a sigmoidal binding isotherm indicating positive cooperativity (*K*_D2_ < *K*_D1_) between the two metal-sites (Fig. 2a). Such cooperativity will result in pairwise filling of the two metal-sites. Given that GTPases typically bind nucleotides in complex with Mg^II^, we hypothesised that the cognate metal for the first (weak-affinity) site is Mg^II^, and that Mg^II^ binding triggers assembly of the second (tight-affinity) metal-site in GMPPNP-CobW. Co^II^-titration of CobW (20 μM) with GMPPNP (60 μM) and Mg^II^ (2.7 mM, *ie* available idealised intracellular concentration, [Mg^II^]_cell_^27,30^) produced identical spectra to that observed without Mg^II^ but saturating at 1:1 ratio Co^II^:CobW (Fig. 2b). Equivalent experiments performed with GTP and GTPγS also revealed 1:1 Co^II^:CobW stoichiometry in the presence of [Mg^II^]_cell_ (Supplementary Fig. 3c,d). Thus, binding of Mg^II^ and guanine nucleotides preassembles one distinct Co^II^ site in CobW. Occupancy of the first site by Mg^II^ was spectroscopically silent in these experiments. The features at 339 nm and at 500 – 700 nm therefore correspond exclusively to a distinct tetrahedral Co^II^ site and the coordinating sulfhydryl side-chains likely derive (at least in part) from the CxCC motif adjacent to the nucleotide-binding site.

**Fig. 2.**
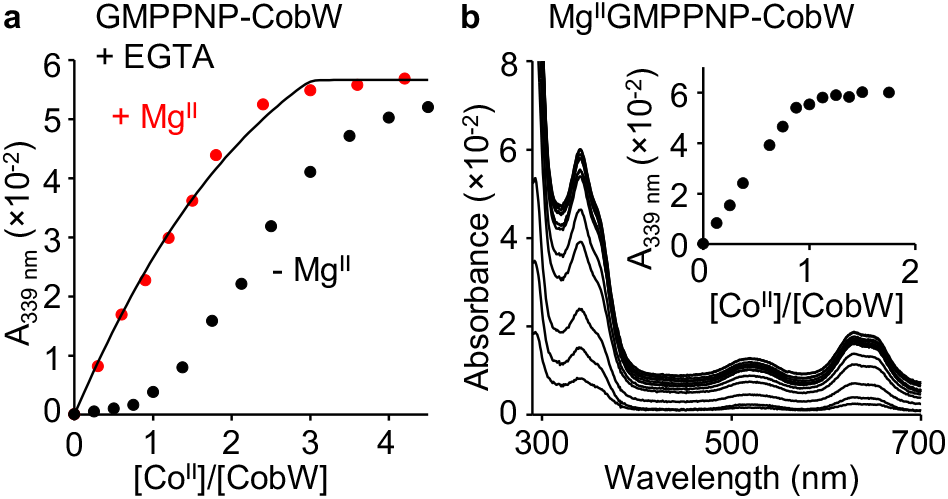
Mg^II^ is also required to assemble a high affinity Co^II^ site in CobW. **a** Absorbance (relative to Co^II^-free solution) of Co^II^-titrated CobW (20 μM) with GMPPNP (60 μM) in competition with EGTA (40 μM); titrations performed in the absence (black) or presence (red) of Mg^II^ (2.7 mM, *ie* intracellular concentration in a bacterium^27,30^). Solid trace shows curve-fit of the experimental data to a model in which CobW binds one molar equivalent of Co^II^ per protein monomer (in the presence of GMPPNP and Mg^II^). **b** Absorbance (relative to Co^II^-free solution) of Co^II^-titrated CobW (20 μM) with GMPPNP (60 μM) and Mg^II^ (2.7 mM) in the absence of competing ligand; feature at 339 nm (inset) showing linear increase saturating at 1:1 ratio Co^II^:CobW.

Mg^II^ had negligible impact on the conditional affinity of EGTA for Co^II^ at the concentrations used here (Supplementary Table 1): For this reason Mg^II^ was not incorporated into curve-fitting models. Competition between Mg^II^GMPPNP-CobW and EGTA for Co^II^ yielded a binding isotherm consistent with 1:1 stoichiometry for both Co^II^:protein and Co^II^:EGTA, and enabled *K*_Co(II)_ of 2.7 (±0.4) ×10^−9^ M for Mg^II^GMPPNP-CobW to be determined (Fig. 2a, Supplementary Fig. 4a,b and Supplementary Tables 2,3). Competition with EGTA revealed a Co^II^ affinity for Mg^II^GTPγS-CobW (*K*_Co(II)_ = 1.7 (±0.8) ×10^−10^ M; Supplementary Fig. 4c-e), that was more than 10-fold tighter than Mg^II^GMPPNP-CobW, establishing that the nature of the bound nucleotide exerts an effect on metal-binding to CobW.

### Co^II^ binds a thousand-fold tighter with GTP than GDP

Observed variation in Co^II^ affinities of CobW in association with Mg^II^GTPγS versus Mg^II^GMPPNP, prompted us to assess the Co^II^ affinities of all three anticipated biological species: nucleotide-free CobW, Mg^II^GTP-CobW and Mg^II^GDP-CobW. Co^II^ affinities of CobW and Mg^II^GDP-CobW were determined via competition with fura-2 (Fig. 3a,b and Supplementary Fig. 4f-i). Fura-2 is too weak to compete effectively with Mg^II^GTP-CobW (Supplementary Fig. 4j), but high concentrations of EGTA or nitrilotriacetic acid (NTA) imposed sufficient competition to enable *K*_Co(II)_ of 3.0 (±0.8) ×10^−11^ M to be determined (Fig. 3c and Supplementary Fig. 4k-m). GTP concentration was not a limiting factor in these affinity measurements (Supplementary Fig. 5). Under identical conditions used for affinity measurements, we confirmed that CobW-catalysed GTP hydrolysis is sufficiently slow such that nucleotides remain predominantly unhydrolysed over the duration of metal-binding experiments (Fig. 3d,e and Supplementary Fig. 6). Mg^II^GDP-CobW, despite displaying identical absorbance features indicating the persistence of the Cys-rich tetrahedral site (Supplementary Fig. 7), has a Co^II^ affinity more than one thousand-fold weaker than Mg^II^GTP-CobW and only marginally tighter than unbound CobW which lacks this site altogether (Supplementary Table 3). GTP also confers higher Co^II^ affinity than either of the tested non-hydrolysable analogues in which the γ-phosphates have been modified (Fig. 1f and Supplementary Table 3). Thus, the presence of an intact nucleotide γ-phosphate is a prerequisite for high-affinity Co^II^ binding.

**Fig. 3.**
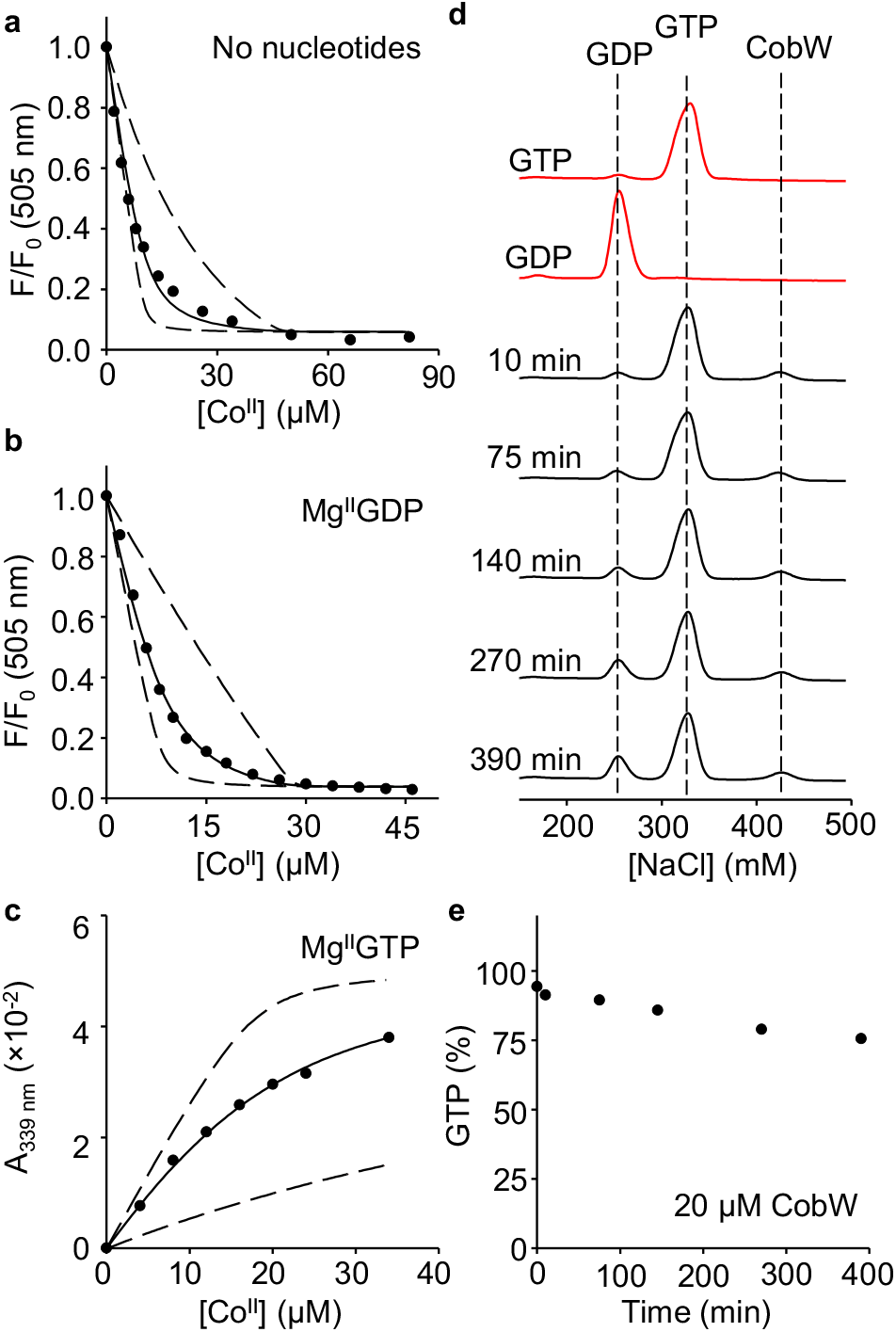
The γ-phosphate group of GTP is necessary for high affinity Co^II^ binding. **a-c** Representative *K*_Co(II)_ quantification for CobW in the absence or presence of nucleotides (n=3 independent experiments, full details in Supplementary Fig. 4 and Supplementary Table 2). Solid traces show curve fits of experimental data to a model where CobW binds one molar equivalent Co^II^ per protein monomer. Dashed lines show simulated responses for *K*_Co(II)_ tenfold tighter or weaker than the fitted value. **a** Fluorescence quenching of Co^II^-titrated Fura-2 (10 μM) in competition with CobW alone (37 μM). **b** Fluorescence quenching of Co^II^-titrated Fura-2 (8.1 μM) in competition with CobW (20 μM) in the presence of Mg^II^ (2.7 mM) and GDP (200 μM). **c** Absorbance (relative to Co^II^-free solution) of Co^II^-titrated CobW (18 μM) in competition with EGTA (2.0 mM) in the presence of Mg^II^ (2.7 mM) and GTP (200 μM). **d** Analysis of GTP hydrolysis by anion-exchange chromatography. Control samples of GTP and GDP elute as distinct peaks (red traces) measured by absorbance at 254 nm. Black traces show the extent of nucleotide hydrolysis when a solution of GTP (200 μM) was incubated with CobW (20 μM), Mg^II^ (2.7 mM) and Co^II^ (18 μM) and analysed at time intervals indicated. **e** Analysis of data from (**d**) showing % GTP remaining over time. After 390 mins incubation nucleotides remain primarily (>75 %) unhydrolysed.

### Cu^I^ and Zn^II^ bind Mg^II^GTP-CobW more tightly than Co^II^

In view of the challenges associated with correct metal-protein speciation, we sought to determine Mg^II^GTP-CobW affinities for other first-row transition metals (Fe^II^, Ni^II^, Cu^I^, Zn^II^). Fe^II^-titration into a mixture of Mg^II^GTP-CobW (50 μM) and probe ligand 4-(2-thiazolylazo)-resorcinol (Tar) (16 μM) showed Fe^II^ being withheld by Tar which revealed a limiting affinity (*K*_Fe(II)_ > 10^−6^ M) for Mg^II^GTP-CobW (Fig. 4a and Supplementary Fig. 8). Competition between Mg^II^GTP-CobW (10 μM) and mag-fura-2 (Mf2; 20 μM) for Ni^II^ showed that Mg^II^GTP-CobW has one Ni^II^-site which outcompetes Mf2 (*K*_Ni(II)_ < 10^−8^ M) in addition to two weaker sites which compete with Mf2 for Ni^II^ (*K*_Ni(II)_ ~ 10^−7^ M) and are also present in CobW alone (Supplementary Fig. 9a). Competition with Tar allowed the affinity of the tight Ni^II^-site in Mg^II^GTP-CobW to be determined (*K*_Ni(II)_ = 9.8 (±6.5) ×10^−10^ M; Fig. 4b and Supplementary Fig. 9b,c). The conditional β_2_ value (4.3 (±0.6) ×10^15^ M^−2^) for Ni(Tar)_2_ formation under experimental conditions (pH 7.0, 10 mM NaCl, 400 mM KCl) was independently established by competition with EGTA (Supplementary Fig. 10). Titration of Mg^II^GTP-CobW (15 μM) and bathocuproine disulfonate (Bcs; 30 μM) with Cu^I^ did not reach the expected intensity at saturating metal concentrations (Supplementary Fig. 11a) suggesting the presence of a stable ternary complex, which would preclude accurate affinity determinations^43^. An equivalent experiment with alternative Cu^I^-probe bicinchoninic acid (Bca) showed that Mg^II^GTP-CobW has two Cu^I^-sites which outcompete Bca and at least three additional weaker Cu^I^ sites which effectively compete with the probe (Supplementary Fig. 11b). Effective competition imposed by excess Bca enabled *K*_Cu(I)_ of 2.4 (±0.9) ×10^−16^ M to be determined (Fig. 4c, Supplementary Fig. 11c,d and Supplementary Fig. 12), assuming only the tightest Cu^I^-site can acquire metal at the limiting Cu^I^ availabilities employed (*eg* [Cu^I^_aq_] < 3×10^−16^ M in Fig. 4c). Zn^II^-titration into a mixture of quin-2 (20 μM) and Mg^II^GTP-CobW (10 μM) revealed one high-affinity Zn^II^-site in the protein which was too tight to be quantified by using quin-2 thus showing *K*_Zn(II)_ < 10^−12^ M (Fig. 4d).

**Fig. 4.**
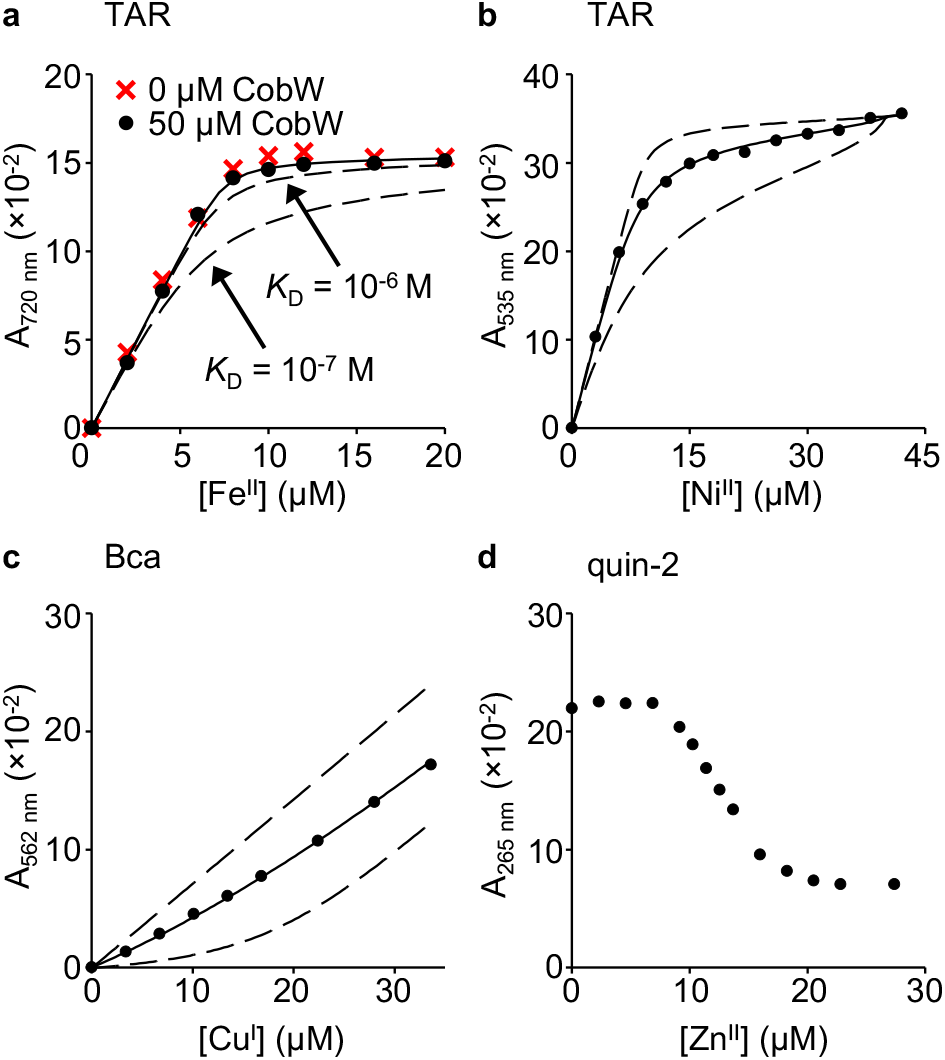
Binding of CobW-Mg^II^GTP to Fe^II^, Ni^II^, Cu^I^ and Zn^II^. **a** Absorbance upon Fe^II^-titration into a mixture of Tar (16 μM), Mg^II^ (2.7 mM) and GTP (500 μM) in the absence or presence of CobW (50 μM). Dashed lines show simulated responses for specified *K*_Fe(II)_ of CobW-Mg^II^GTP, providing limiting *K*_Fe(II)_ ≥ 10^−6^ M. Control Fe^II^-titration into a solution of Tar (16 μM) in buffer only (Supplementary Fig. 8a) confirmed that Mg^II^ and GTP did not inhibit stoichiometric Fe^II^Tar_2_ formation. **b** Absorbance change (relative to Ni^II^-free solution) of Ni^II^-titrated Tar (20 μM) in competition with CobW (30 μM) in the presence of Mg^II^ (2.7 mM) and GTP (300 μM). **c** Absorbance of Cu^I^-titrated Bca (1.0 mM) in competition with CobW (20 μM) in the presence of Mg^II^ (2.7 mM) and GTP (200 μM). In (**a-c**) solid traces show curve fits of experimental data to models where CobW binds one molar equivalent of metal per protein monomer. Supplementary Table 3 contains mean ± s.d. *K*_metal_ values from n=3 independent experiments (full details in Supplementary Figs. 8-12 and Supplementary Table 2). In (**b-c**) dashed lines show simulated responses for *K*_metal_ tenfold tighter or weaker than the fitted value. **d** Absorbance (relative to probe-free solution) upon titration of Zn^II^ into a mixture of quin-2 (10 μM), Mg^II^ (2.7 mM), GTP (100 μM) and CobW (10 μM).

Because of the limiting affinity of quin-2 we employed inter-metal competition, which presumably also occurs within the buffered intracellular milieu, to determine *K*_Zn(II)_ for Mg^II^GTP-CobW. *K*_Zn(II)_ was determined, relative to the known *K*_Co(II)_, via competition between the two metals. This approach required an excess of metal ions competing for a limited number of protein metal-sites (*ie* [Co^II^]_tot_ + [Zn^II^]_tot_ > [CobW]_tot_) thus it was essential to include a buffering ligand, in this case NTA, to control the speciation of all Co^II^ and Zn^II^ in solution (*ie* [NTA]_tot_ > [Co^II^]_tot_ + [Zn^II^]_tot_). The measured equilibrium (*K*_ex_ in Fig. 5a) was the exchange constant for Co^II^/Zn^II^ exchange between the protein (Mg^II^GTP-CobW) and buffering ligand (NTA). Equilibrium ratios of [Co^II^Mg^II^GTP-CobW]/[Zn^II^Mg^II^GTP-CobW] were determined (Fig. 5b-e and Supplementary Table 4): absorbance intensity at A_339 nm_ reported specifically on the Co^II^-protein complex and all remaining protein was Zn^II^-bound (since Mg^II^GTP-CobW was metal-saturated under experimental conditions; Supplementary Fig. 13). The concentrations of NTA-bound metals were determined from mass balance relationships (equations (6–8) in Methods). Experiments were conducted at multiple relative availabilities of Co^II^ and Zn^II^ and reciprocally (Fig. 5b-e), with consistent results (Supplementary Table 4), to confirm reliability of measured affinities. We thus determined a tight *K*_Zn(II)_ of 1.9 (±0.6) × 10^−13^ M for Mg^II^GTP-CobW (Supplementary Table 3).

**Fig. 5.**
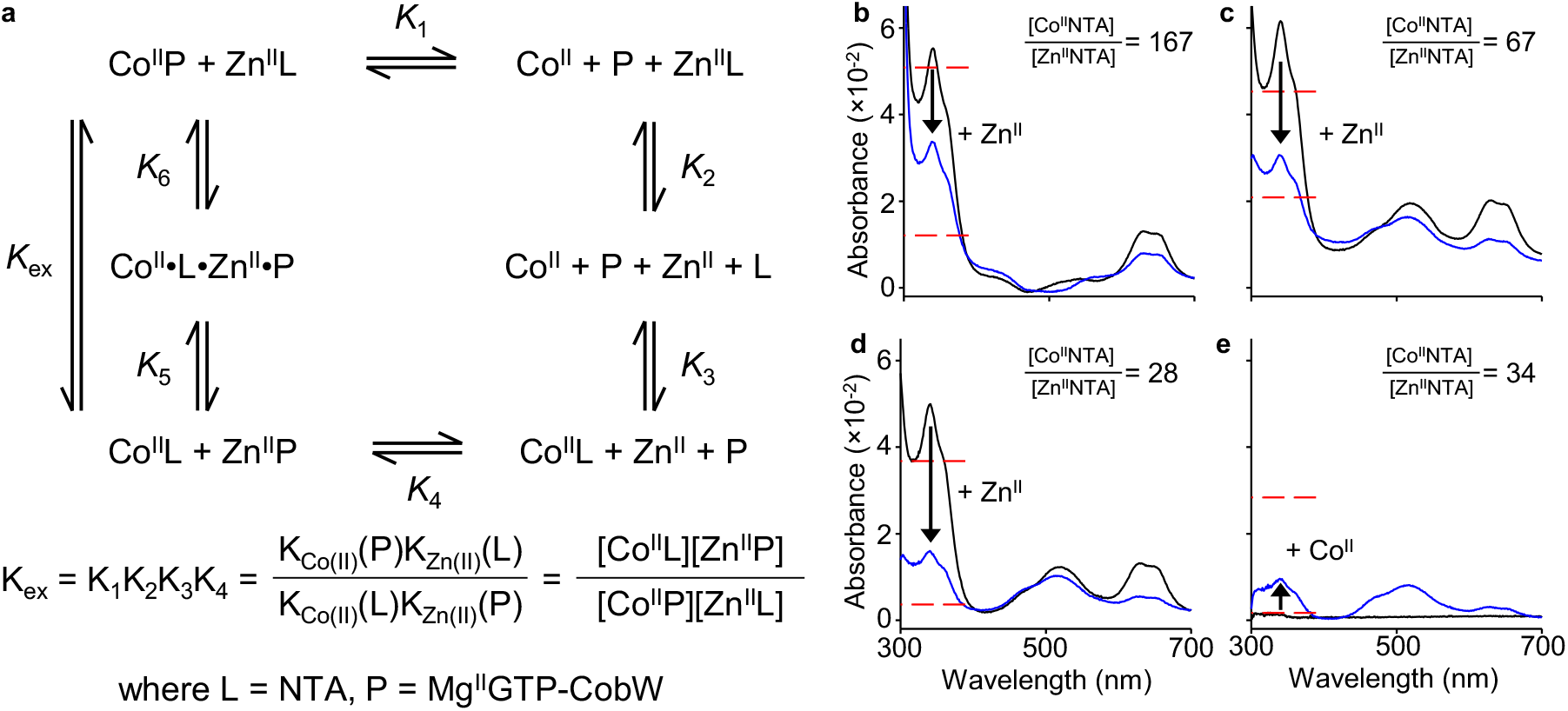
CobW-Mg^II^GTP binds Zn^II^ with sub-picomolar affinity. **a** Representation of the equilibrium for exchange of Co^II^ and Zn^II^ between ligand (L = NTA) and protein (P = Mg^II^GTP-CobW). **b-e** Absorbance (relative to metal-free solution) of solutions of CobW (17.9 – 20.4 μM), Mg^II^ (2.7 mM), GTP (200 μM) and NTA (0.4 – 4.0 mM) upon (**b – d**) first addition of Co^II^ (black trace) then Zn^II^ (blue trace) or (**e**) the reverse, at equilibrium. The absorbance peak at 339 nm corresponds to Co^II^-bound protein. An excess of ligand NTA was used to buffer both metals in each experiment: varying the ratios of ligand-bound metal ions ([Co^II^NTA]/[Zn^II^NTA] = 28 – 167) shifted the ratios of Co^II^- and Zn^II^-bound protein as predicted by the equilibrium exchange constant in (**a**). Consistent *K*_Zn(II)_ values for Mg^II^GTP-CobW were generated at all tested conditions (Supplementary Table 4). Dashed red lines show expected A_339 nm_ peak intensities for *K*_Zn(II)_ of Mg^II^GTP-CobW 10-fold tighter or weaker than calculated values.

### GTP not GDP will enable Co^II^ acquisition in cells

In the same manner that Fig. 4 considered competition between a ligand (Tar, Bca or quin-2) and a protein (Mg^II^GTP-CobW) for metal-binding *in vitro*, metal acquisition by proteins *in vivo* likewise involves competition with a surplus of cytosolic ligands that buffer metals to different availabilities^8,27,32,44,45^. Recent work has estimated the buffered availabilities of metals M (where M = Mg^II^, Mn^II^, Fe^II^, Co^II^, Ni^II^, Cu^I^, Zn^II^) in a reference bacterium (*Salmonella*^27^) expressed here as free energies (Δ*G*; Fig. 6). The intracellular available Δ*G* for each metal, Δ*G*_M_, is defined as the free energy required for a ligand to become 50% metalated from available and exchangeable intracellular metal (see Supplementary Note 1). Fig. 6 and Supplementary Fig. 14 show the intracellular available Δ*G*_M_ values in an ‘idealised cell’ (*ie* neither metal-deficiency nor –excess) defined as the metal availabilities at which each cognate sensor undergoes half of its transcriptional response. Bars show the changes in available intracellular Δ*G*_M_ as sensors shift from 10 – 90% (Fig. 6) or 1 – 99% (Supplementary Fig. 14) of their respective responses. The percentage occupancy of a protein, P, with metal, M, *in vivo* is governed by the difference between the free energy for protein metalation, Δ*G*_MP,_ and the intracellular available Δ*G*_M_ (equation (1)) and can be calculated via equation (2) (see Supplementary Note 1):

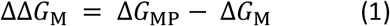

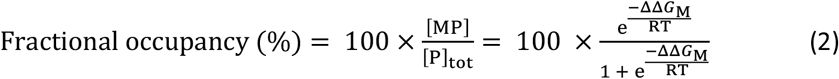

**Fig. 6.**
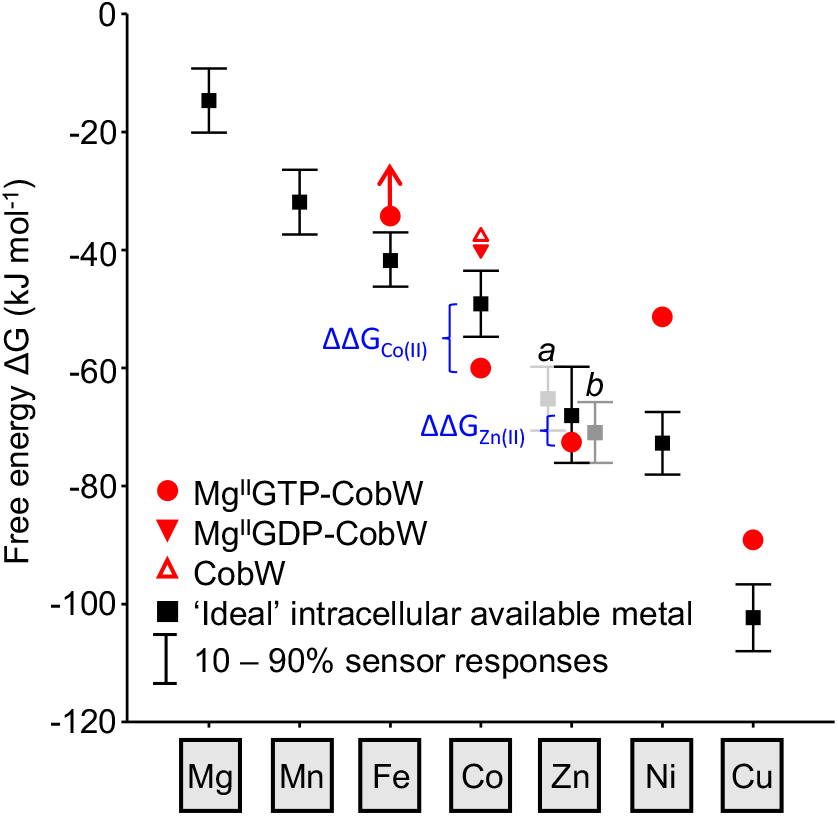
Mg^II^GTP-CobW is predicted to acquire Co^II^ or Zn^II^ in a bacterial cell. Free-energy change (Δ*G*) for metal-binding to Mg^II^GTP-CobW plotted against the intracellular available free energies for metal-binding in a reference bacterial cytosol (values correspond to *Salmonella*) under idealised conditions (*ie* where each metal sensor undergoes half of its transcriptional response) (ref.^27^). Intracellular available Δ*G*_Zn(II)_ is the mean of the values determined from the two Zn^II^-sensors ZntR (*a*) and Zur (*b*). Bars shows the change in intracellular available Δ*G* as cognate sensors shifts from 10-90% of their responses. Free energy differences (ΔΔ*G*) which favour acquisition of metals by Mg^II^GTP-CobW *in vivo* are indicated. Δ*G* values for Co^II^-complexes of CobW alone and Mg^II^GDP-CobW are also shown. For Fe^II^ binding to Mg^II^GTP-CobW, arrow indicates limiting Δ*G* > −34.2 kJ mol^−1^.

In an idealised cell, the Δ*G*_Co(II)_ for CobW and Mg^II^GDP-CobW were both significantly more positive than intracellular available Δ*G*_Co(II)_ (ΔΔ*G*_Co(II)_ ≫ 0; Fig. 6) resulting in negligible Co^II^-occupancies of 1.0% and 2.5% for these two protein forms, respectively. Conversely, Δ*G*_Co(II)_ for Mg^II^GTP-CobW was significantly more negative than intracellular available Δ*G*_Co(II)_ (ΔΔ*G*_Co(II)_ ≪ 0), resulting in almost complete protein metalation (99%). Thus, CobW needs Mg^II^GTP to acquire Co^II^ in a cell.

### Mg^II^GTP-CobW may also acquire Zn^II^

In addition to Co^II^ other metals also bound to Mg^II^GTP-CobW (Figs. 4 and 5). However, ΔΔ*G* for Fe^II^, Ni^II^ and Cu^I^, was significantly greater than zero (equation (1) and Fig. 6), thus preventing acquisition of these metals (equation (2) and Table 1). In contrast, ΔΔ*G*_Zn(II)_ was < 0 with *in vivo* Zn^II^ occupancy predicted to be 86% (Fig. 6 and Table 1). However, based on equation (2) the sum of metal occupancies of Mg^II^GTP-CobW gave an impossible total metalation > 100% (Table 1). Since ΔΔ*G* was < 0 for both Co^II^ and Zn^II^, a more sophisticated approach needs to account for competition between multiple buffered metals in order to predict how much Zn^II^ binds Mg^II^GTP-CobW *in vivo*.

**Table 1.** Calculated metal occupancies of CobW-Mg^II^GTP in an idealised cell ^*a*^

| Metal | Occupancy |  |
| --- | --- | --- |
|  | Equation 2<sup>b</sup> | Equation 4<sup>c</sup> |
| Fe<sup>II</sup> | < 4.6 % | < 0.1 % |
| Co<sup>II</sup> | 98.8 % | 91.9 % |
| Zn<sup>II</sup> | 86.2 % | 6.9 % |
| Ni<sup>II</sup> | 0.1 % | 0.0 % |
| Cu<sup>I</sup> | 0.5 % | 0.0 % |
| TOTAL | 190.3 % | 98.9 % |

<sup>a</sup>Based on metal availabilities in *Salmonella* under idealised conditions (ref.<sup>27</sup>)
<sup>b</sup>Does not account for inter-metal competition between different metals for the same high- affinity site in Mg<sup>II</sup>GTP-CobW.
<sup>c</sup>Takes into account competition between multiple intracellular metals for the same site in Mg<sup>II</sup>GTP-CobW.

### Calculating inter-metal competition in a cell

Figure 5 considered competition between Co^II^ and Zn^II^ for a single metal-binding site in a protein (Mg^II^GTP-CobW) when the metals were buffered to different availabilities *in vitro* by an excess of NTA. This can be represented as an available Δ*G*_M_ (Supplementary Table 4). The observed Co^II^ occupancy was a function of the protein’s affinities for both Co^II^ and Zn^II^ relative to their buffered availabilities in solution (*ie* ΔΔ*G* values), as described by equation (3) (see Supplementary Note 1).

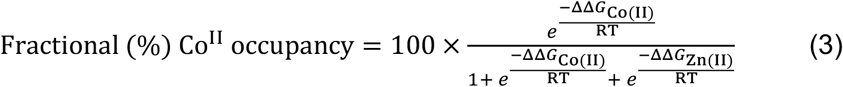

By analogy in a cytoplasm multiple metals, each buffered to different intracellular available Δ*G*_M_, compete for a single protein-binding site. We generalised equation (3) to account for n different metals (equation (4) and Supplementary Note 1).

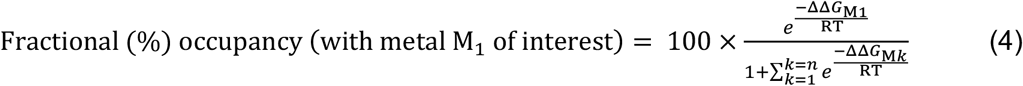

Thus, we developed a metalation calculator (Supplementary Data 1) for determining *in vivo* metal occupancies of proteins, accounting for multiple inter-metal competitions plus competition from components of the intracellular milieu.

### Co^II^ specificity under idealised conditions

Since ΔΔ*G* was < 0 for binding of both Co^II^ and Zn^II^ to Mg^II^GTP-CobW (Fig. 6), equation (4) was next used to predict *in vivo* metalation in an idealised cell. Between the five metals considered (Fe^II^, Co^II^, Ni^II^, Cu^I^ and Zn^II^), Mg^II^GTP-CobW will favour Co^II^-binding in a cell and calculations via equation (4) predicted occupancies of 92% and 7%, for Co^II^ and Zn^II^, respectively (Table 1). Thus, although Mg^II^GTP-CobW affinities for both Co^II^ and Zn^II^ are tight enough to extract either metal from the cytosolic buffer, Co^II^ will outcompete Zn^II^, rationalising specificity but only in an intracellular context where there is competition from other cellular components.

### Fine tuning ΔG for metalation in a cell

Calculated free energies for intracellular metalation (Δ*G*_M_) in Fig. 6 are based on an assumption that cellular metal availabilities are fixed at ‘ideal’ buffered concentrations where every metal sensor undergoes half of its transcriptional response (*ie* normalised fractional DNA occupancy ‘*θ*_D_’ = 0.5, see ref.^27^). In reality cellular metal availabilities, and consequently *θ*_D_ of sensors, fluctuate conditionally (*eg* in response to addition of metals or chelators to the growth media). For example, the dynamic response range (defined as *θ*_D_ = 0.99 – 0.01) of RcnR, the Co^II^ sensor from *Salmonella*, coincides with an increase in the intracellular available [Co^II^] from 2.4 × 10^−11^ to 2.7 × 10^−7^ M, corresponding to an increase in intracellular available Δ*G*_Co(II)_ from −60.6 to −37.5 kJ mol^−1^ (Fig. 7a and Supplementary Table 5).

**Fig. 7.**
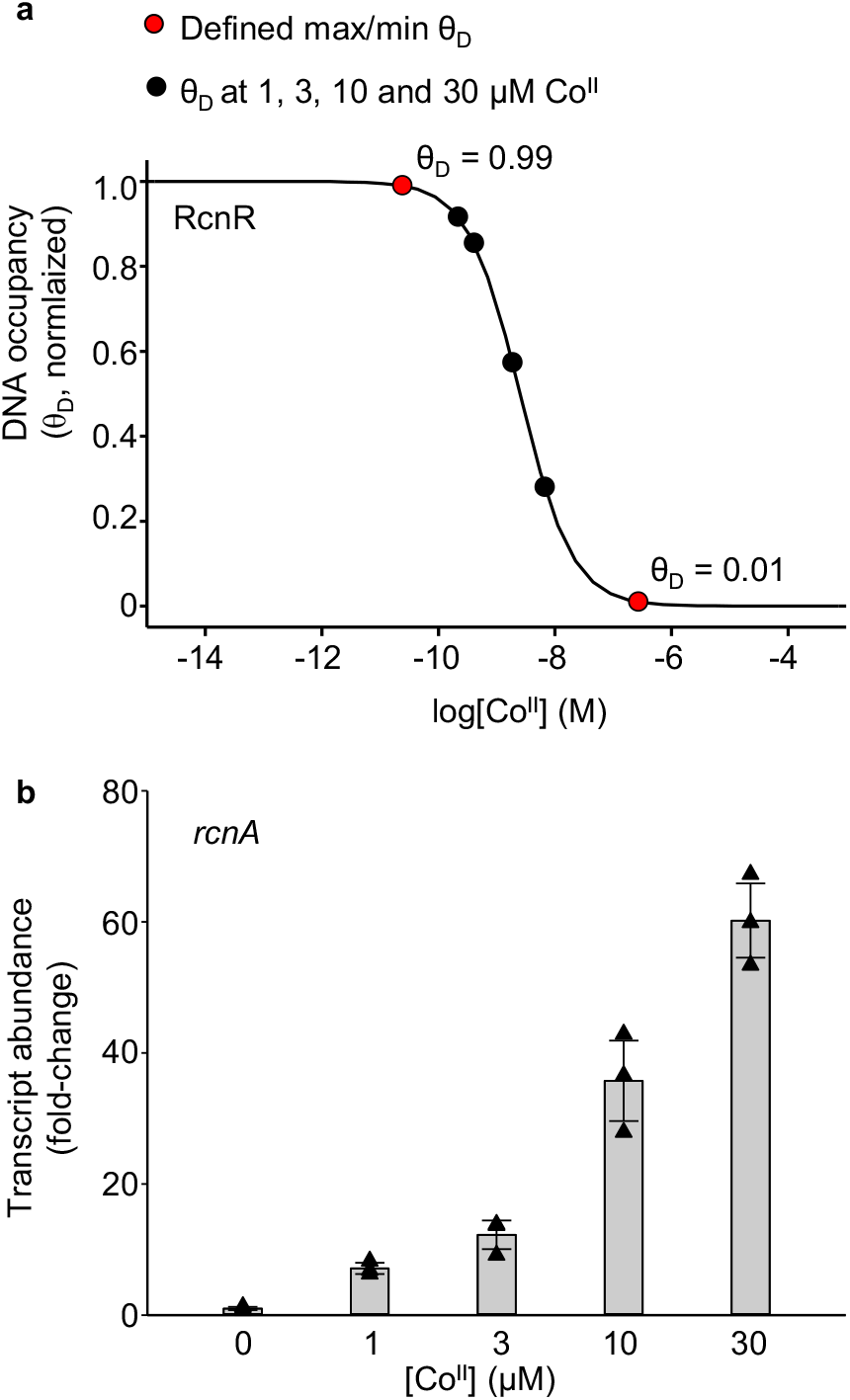
Calculations of conditional Co^II^ availabilities in B_12_-producing *E. coli\**. **a** Calculated relationship between intracellular Co^II^ availability and normalised DNA occupancy (*θ*_D_) by RcnR. *θ*_D_ of 0 and 1 are the maximum and minimum calculated DNA occupancies. The dynamic range (within which RcnR responds to changing intracellular Co^II^ availability) has been defined as *θ*_D_ of 0.01 to 0.99 (*ie* 1 – 99% of RcnR response). The calibrated maximum and minimum fold changes in *rcnA* transcript abundance (*ie* boundary conditions, see Supplementary Fig. 15) therefore correspond to *θ*_D_ of 0.01 and 0.99 in these calculations. *θ*_D_ for each growth condition (black circles) was calculated from the qPCR response in **b**, assuming a linear relationship between change in *θ*_D_ and change in transcript abundance (equation (9)). Corresponding Co^II^ availabilities were calculated as previously described^27^ and are listed in Supplementary Table 5. **b** Transcript abundance (relative to untreated control) of the RcnR-regulated gene *rcnA* following 1h exposure of *E. coli\** to increasing [Co^II^], measured by qPCR. Data are the mean ± s.d. of n=3 biologically independent replicates. Triangle shapes represent individual experiments.

In order to account for this variation, we developed a method to fine-tune the free energy calculations for Co^II^ under bespoke culture conditions using qPCR analysis of the RcnR-regulated gene *rcnA*. Fine-tuning was performed in *E. coli\** which has been engineered to synthesise vitamin B_12_ (*E. coli* and *Salmonella* RcnR share 93% sequence identity and equivalent responses to available Co^II^ were assumed). *E. coli\** cells were cultured in standard medium with increasing Co^II^ supplementation. The *rcnA* transcript abundance (Fig. 7b) was used to calculate *θ*_D_ of RcnR for each condition (via equation (9) in Methods) following calibration of the maximum and minimum responses (defined as *θ*_D_ = 0.99 and 0.01 at low and high [Co^II^] respectively; Supplementary Fig. 15). This enabled the intracellular Co^II^ availabilities, as conditional free energies, to be calculated from the RcnR *θ*_D_ for each condition (Fig. 7a, Supplementary Table 5).

### Co^II^-acquisition by Mg^II^GTP-CobW predicts B_12_ (corrinoid) synthesis

Does the amount of Co^II^ inserted into B_12_ follow the predicted metalation of Mg^II^GTP-CobW? Metal occupancies of Mg^II^GTP-CobW in *E. coli\** samples were recalculated (via equation (4)) using bespoke intracellular available free energies, Δ*G*_Co(II)_, for each growth condition (Fig. 7 and Supplementary Table 5). This predicted that in unsupplemented LB media the protein would be predominantly Zn^II^-bound (10% Co^II^ and 77% Zn^II^) and that Co^II^ occupancies would increase from 10% to 97% as added [Co^II^] increased from 0 – 30 μM (Fig 8a). Since intracellular Zn^II^ availability was also significant in our predictions, we confirmed that our previous estimation of Δ*G*_Zn(II)_ was valid in LB media (Supplementary Fig. 16). Corrin concentrations (presumed to be predominantly B_12_) were measured in *E. coli\** strains containing or missing *cobW* (Fig. 8b and Supplementary Fig. 17), under the growth conditions for which intracellular available Δ*G*_Co(II)_ was defined (Supplementary Table 5). As the added [Co^II^] increased so did B_12_ production in *cobW(+)*, consistent with the predicted loading of Mg^II^GTP-CobW with Co^II^ (Fig 8). At higher [Co^II^], CobW-independent B_12_ synthesis became evident. Notably, the synthesis of B_12_ which is dependent on CobW (Fig. 8b, compare *cobW*(+) with *cobW*(−)) closely matches the predicted metalation of Mg^II^GTP-CobW (Fig 8a).

**Fig. 8.**
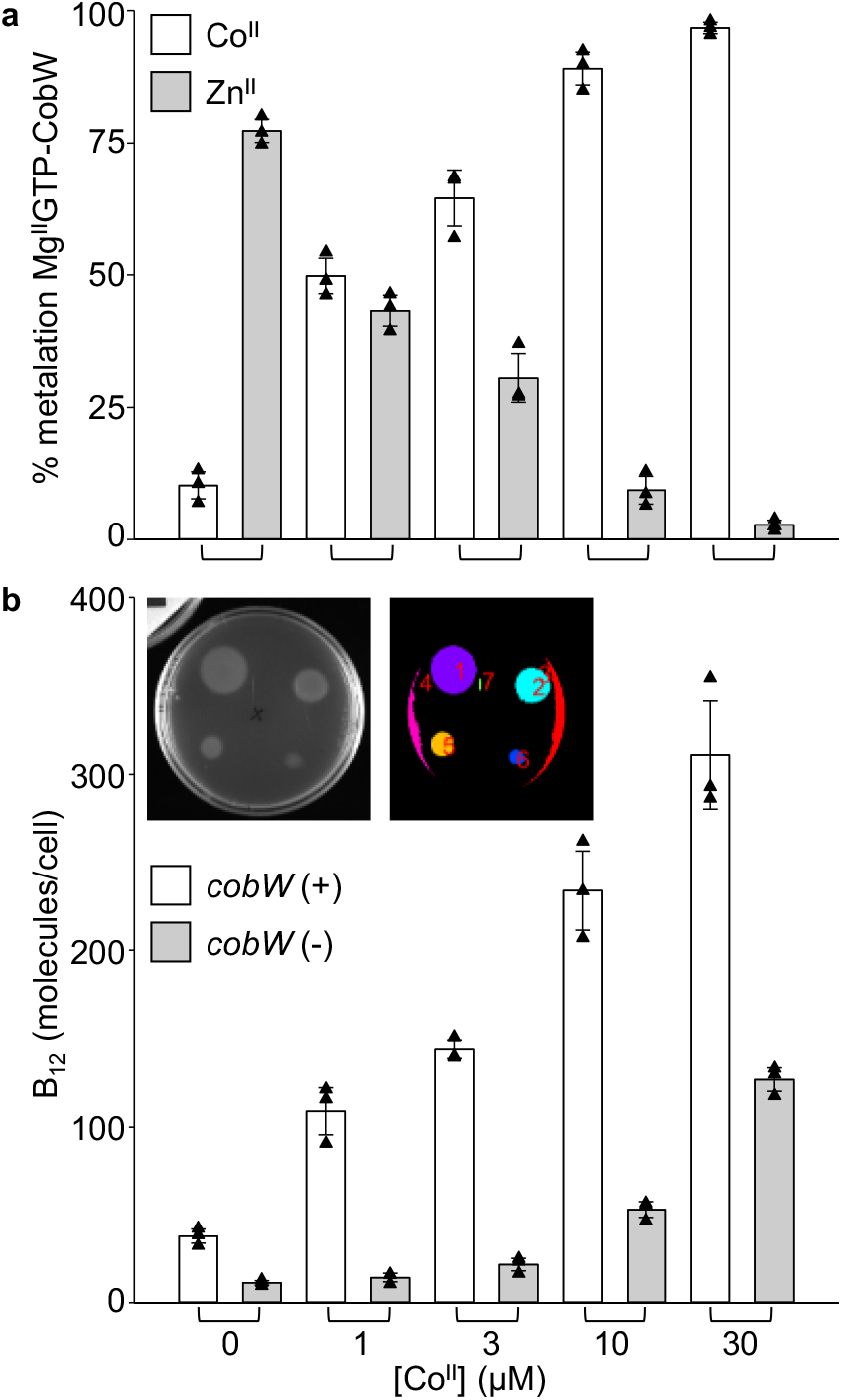
B_12_ production follows predicted metalation of Mg^II^GTP-CobW. **a** Predicted metalation of Mg^II^GTP-CobW with Co^II^ and Zn^II^ in samples treated with defined media [Co^II^]. Intracellular Δ*G*_Co(II)_ for each condition was calculated from *rcnA* expression (Fig. 7 and Supplementary Table 5). **b** B_12_ produced by *E. coli\** strains with and without *cobW* (open and grey bars, respectively) following 4h exposure to defined [Co^II^]. B_12_ was detected using a *Salmonella* AR2680 bioassay^37^ (detects corrins, expected to be predominantly B_12_; see Methods) and quantified by automated analysis of growth areas (Supplementary Fig. 17 and Supplementary Note 2). Inset shows original image and detected areas for representative (n=3) bioassay plate of B_12_ calibration standards. All data are the mean ± s.d. of n=3 biologically independent replicates (with errors in **a** propagated from qPCR data in Fig 7b). Triangles represent individual experiments.

## Discussion

CobW belongs to a ubiquitous family of putative metallochaperones (COG0523) but its cognate metal, target protein(s) and mechanism of action were undefined. Here we establish the connection between CobW and Co^II^ (Figs. 1-8). We show how CobW can acquire Co^II^ in a cell (Figs. 1-3, Fig. 6 and Table 1). Free-energy calculations reveal that in an idealised cell Co^II^ ions will not flow from the cellular milieu to nucleotide-free CobW (ΔΔ*G*_Co(II)_ > 0). Crucially, Co^II^ will flow from the cellular milieu to the Mg^II^GTP form of CobW (ΔΔ*G*_Co(II)_ < 0) (Fig. 6, Fig. 9a, Table 1 and Supplementary Table 3). Thus, CobW must first bind Mg^II^GTP in order to acquire Co^II^ inside a cell. In contrast, the product of GTP hydrolysis, Mg^II^GDP-CobW, will release Co^II^ to the cellular milieu (ΔΔ*G*_Co(II)_ > 0) (Fig. 6, Fig. 9b, Table 1 and Supplementary Table 3). Thus, the GTPase activity of CobW will facilitate Co^II^ release for example to CobNST for insertion into the corrin ring of B_12_ (Fig. 3d,e and Supplementary Fig. 6). We establish that CobW enhances B_12_ production when Co^II^ is limiting (Fig. 8b), and Fig. 9 illustrates the proposed mechanism.

**Fig. 9.**
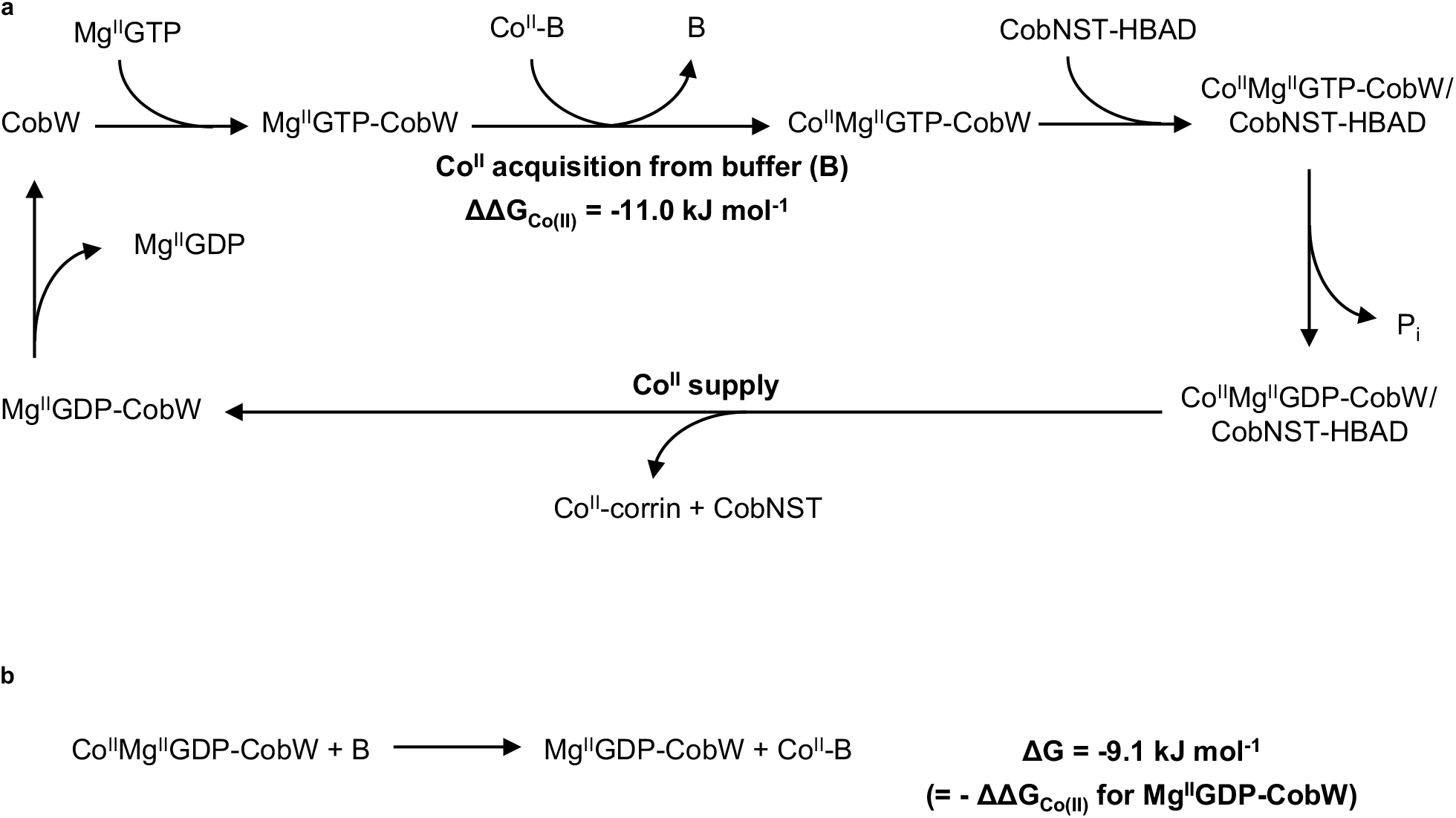
Proposed mechanism of CobW. **a** Binding of Mg^II^GTP enables CobW to acquire Co^II^ from intracellular buffer ligands (B) followed by interaction with the CobNST cobaltocheletase bound to its substrate hydrogenobyrinic acid *a,c*-diamide (HBAD). GTP hydrolysis will trigger Co^II^ release to CobNST-HBAD, since the reaction in **b** is thermodynamically favourable, for incorporation into the corrin ring of vitamin B_12_.

The intrinsic GTPase activity of CobW is slow (Fig. 3d,e and Supplementary Fig. 6), as observed for other COG0523 proteins^13–15,17^. Giedroc and co-workers hypothesised that interactions with partner proteins may stimulate GTP hydrolysis in similar proteins^17^.

Likewise, we speculate that CobNST could act as a guanine nucleotide activating protein (GAP) enabling Co^II^ release to be targeted to the cobaltocheletase. Release of Co^II^ mediated by CobNST acting as a guanine nucleotide exchange-factor (GEF) is also formally possible^46^. By analogy to ZTP-ZagA^18^, GTP-binding (and subsequent metal-acquisition) by CobW could promote interaction with CobNST and contribute to the reaction cycle (Fig. 9). Dissociation of Mg^II^GDP (or nucleotide exchange), resets the reaction cycle with GTPases thought to be saturated with nucleotide (either GTP or GDP) inside cells^47^.

Initial calculations here, and in previous work^27^, assume an idealised cell in which the metal sensors are at the mid-points of their dynamic ranges (*θ*_D_ = 0.5). Therefore, we have calculated the available Δ*G*_Co(II)_ in real (conditional) cells from the responses of RcnR (*θ*_D_) estimated experimentally by qPCR of *rcnA* (Fig. 7 and Supplementary Fig. 15). The observation that *R. capsulatus* CobW functions in *E. coli* cells suggests overlap in the dynamic ranges for Δ*G*_Co(II)_ in these two bacteria, although evidence here of limited metalation in LB without additional Co^II^ could be a function of the heterologous host (Fig. 8). Notably, a dedicated Co^II^ import system found in *R. capsulatus* (CbiMNQO) is not present in *E. coli* ^48^. As with other metallochaperones^28,49^, CobW is crucial when the cognate metal is limiting but at elevated Co^II^, CobW-independent synthesis of B_12_ occurs (Fig. 8b). CobNST must acquire Co^II^ directly from the cytosol at the higher available Δ*G*_Co(II)_. Importantly, CobW-dependent B_12_ synthesis tracked with the calculated Co^II^ occupancy of Mg^II^GTP-CobW in cells supplemented with different amounts of Co^II^ (Fig. 8). This is an encouraging first test of this approach, and of the easy-to-use spreadsheet (Supplementary Data 1), to calculate changes in the metalation state of a protein inside cells.

Mg^II^GTP-CobW binds Zn^II^ and Cu^I^ more tightly than Co^II^ (Fig. 3c, Fig. 4, Fig. 5, Supplementary Table 3). Notably, by taking into account intracellular metal availability, ΔΔ*G* for Cu^I^ was shown to be greater than zero in an idealised cell (Fig. 6), and also in conditional cells at either 90% or 99% of the dynamic range of the Cu^I^ sensor CueR (Fig. 6 and Supplementary Fig. 14). Thus Mg^II^GTP-CobW will not acquire Cu^I^. However, ΔΔ*G* for Zn^II^ was below zero in an idealised cell suggesting that Mg^II^GTP-CobW is at risk of mis-metalation with Zn^II^ (Fig. 6). Indeed, given that CobW binds Zn^II^ more tightly than many known Zn^II^-proteins^32,50^, it is remarkable that Zn^II^ is not the cognate metal. The data in Figure 5, plus Supplementary Table 4, illustrate how occupancies of Mg^II^GTP-CobW with Co^II^ versus Zn^II^ change as a function of change in relative buffered metal availabilities. By reference to intracellular available free energies, the metal with the most negative ΔΔ*G* will have the highest occupancy *in vivo* (equation (4)). In an idealised cell, ΔΔ*G* for Co^II^ is more negative than ΔΔ*G* for Zn^II^ and so the weaker binding metal dominates (Fig. 6, Supplementary Table 3). In conditional cells without added Co^II^, ΔΔ*G* for Zn^II^ becomes more negative than ΔΔ*G* for Co^II^ and the calculations show binding of Zn^II^ dominating (Fig. 8a). The previously intractable challenge to understand inter-metal competition in a cell now becomes tractable (Supplementary Data 1). Metallochaperones and chelatases may introduce kinetic contributions to the partitioning of metals and these can now become evident in departures from the thermodynamic predictions of the metalation calculator spreadsheet (Supplementary Data 1).

Future structural studies are necessary to understand how Mg^II^GTP-binding facilitates high affinity Co^II^ binding to CobW. Spectral features indicate that the Co^II^ site in Mg^II^GTP-CobW involves thiols, likely derived from the CxCC motif in the GTPase domain, and a tetrahedral geometry (Figs. 1, 2 and Supplementary Fig. 3). All COG0523 proteins contain the CxCC motif^10^, including those that putatively handle Fe^II^ (Nha3)^12,13,51^ and Zn^II^ (YeiR, YjiA, ZigA, ZagA)^14–16,18^. Differences in coordination spheres may alter the ΔΔ*G* values sufficiently to adjust the specificities of these proteins with respect to available intracellular Fe^II^, Co^II^ and Zn^II^. Intriguingly, Ni^II^-binding to Mg^II^GTP-CobW does not follow the order of stabilities of metal-binding predicted by the Irving-Williams series (Fig. 6). An appealing explanation is that the allosteric coupling of GTP- and metal-binding imposes a (tetrahedral) geometry on the metal site that would disfavour Ni^II^-coordination (the Irving-Williams series applies where there is no steric selection): Notably, related G3E GTPases involved in Ni^II^ homeostasis (HypB and UreG) display square planar Ni^II^-coordination geometry^52,53^.

In conclusion, CobW is calculated to be selective for acquiring Co^II^ in its Mg^II^GTP form under conditions of ideal metallostasis, but at risk of erroneously binding Zn^II^ when intracellular Co^II^ is low or Zn^II^ is high (Figs. 6, 7 and 8a). The lack of a dedicated Co^II^ import system could make under-metalation with Co^II^ (and resultant mis-metalation with Zn^II^) especially problematic in *E. coli ^48^*. This has tantalising implications for engineering bacterial strains suited to the manufacture of vitamin B_12_, either via enhanced Co^II^ uptake or impaired Zn^II^ accumulation. More generally, with so many enzymes requiring metals, an ability to calculate *in vivo* metalation should have widespread utility in industrial biotechnology (Supplementary Data 1).

## Methods

### CobW expression and purification

The DNA sequence coding CobW was amplified by PCR using primers 1 and 2 (Supplementary Table 6) with genomic DNA from *Rhodobacter capsulatus* SB1003 as template. The amplified fragment contained an NdeI restriction site at the 5’ end and a SpeI site at the 3’ end, allowing it to be cloned into a modified pET-3a vector as previously described^39^. *E. coli* pLysS, transformed with this pET-3a-CobW plasmid, were cultured in LB medium with antibiotics carbenicillin (100 mg L^−1^) and chloramphenicol (34 mg L^−1^). At mid-log phase, protein expression was induced with IPTG (0.4 mM) at 37°C (3-4h). Cells were resuspended in 20 mM sodium phosphate pH 7.4, 500 mM NaCl, 5 mM imidazole, 5 mM DTT and 1 mM PMSF for lysis (sonication). Lysate was loaded to a 5 mL HisTrap HP column (GE Heathcare) pre-equilibrated in suspension buffer. CobW binds to the HisTrap column courtesy of a natural His-rich region within the protein. The column was washed with suspension buffer (10 CVs), then eluted with suspension buffer containing 100 mM imidazole. Protein-containing fractions were incubated with excess (≥10-fold) EDTA for ≥ 1h before being loaded to a HiLoad 26/600 Superdex 75 size exclusion column equilibrated in 50 mM Tris pH 8.0, 150 mM NaCl, 5 mM DTT and eluted in the same buffer. Peak CobW-containing fractions (determined from SDS-PAGE) were pooled, concentrated to ~0.5 mL (using a Vivaspin® 15 Turbo centrifugal concentrator) then transferred to an anaerobic chamber. The sample was applied to a PD-10 Sephadex G-25 gel-filtration column (GE Healthcare) equilibrated in deoxygenated chelex-treated buffer (10 mM HEPES pH 7.0, 100 mM NaCl, 400 mM KCl) and eluted in the same buffer. Purified CobW samples were quantified by A_280 nm_ using extinction coefficient ε = 15, 300 cm^−1^ M^−1^ determined by quantitative amino acid analysis (performed by Alta Bioscience Ltd). Samples were confirmed to be of high purity (by SDS-PAGE) and ≥95% metal-free (by inductively coupled plasma-mass spectrometry; ICP-MS). ICP-MS was conducted using Durham University Bio-ICP-MS Facility. Protein cysteines were ≥ 90% reduced, determined by reaction with ~10-fold excess of Ellman’s reagent 5,5’-dithio-bis-[2-nitrobenzoic acid] (produces one equivalent of chromophore TNB^2−^ per protein thiol, A_412 nm_ = 14,150 cm^−1^ M^−1^)^54,55^.

Protein identity was confirmed using electrospray ionisation mass spectrometry (ESI-MS) by Durham University Department of Chemistry Mass Spectrometry Service. ESI-MS data were recorded on a QtoF Premier mass spectrometer coupled to an Acuity UPLC system (Waters). Protein samples were desalted prior to injection using a Waters MassPrep desalting cartridge (2.1 ×10 mm) and eluted with a linear acetonitrile gradient (20–80% v/v; 0.1% formic acid). Spectra were processed using Masslynx 4.1 and deconvoluted using MaxEnt 1.

### Preparation of metal stocks

All metal stocks were prepared in ultrapure water from appropriate salts (MgCl_2_, (NH_4_)_2_Fe(SO_4_)_2_, CoCl_2_, NiSO_4_, CuSO_4,_ ZnCl_2_) and quantified by ICP-MS analysis. Fe^II^ stocks were prepared by dissolving (NH_4_)_2_Fe(SO_4_)_2._6H_2_O in deoxygenated 0.1% (v/v) HCl in an anaerobic chamber. Reaction with excess ferrozine (~ 50-fold) confirmed that iron was ≥ 95% reduced (Fe^II^Fz_3_ ε_562 nm_ = 27,900 cm^−1^ M^−1^)^56^. Concentrated stocks were diluted daily in deoxygenated ultrapure water to prepare working solutions and confirmed to be ≥ 90% Fe^II^. Other metal stocks were prepared aerobically as concentrated stocks and diluted to working solutions with deoxygenated ultrapure water in an anaerobic chamber.

### Determination of Co^II^-binding stoichiometries

Metal-binding experiments were conducted in an anaerobic chamber in deoxygenated, chelex-treated 10 mM HEPES pH 7.0, 100 mM NaCl, 400 mM KCl. For stoichiometry determinations, Co^II^ was titrated into a solution of CobW (15 – 30 μM) together with relevant nucleotides (supplied in ~10-fold excess of protein concentration for GTP and GDP and ~3-fold excess for GMPPNP and GTPγS, as specified in figure legends) in the absence or presence of Mg^II^ (2.7 mM). Absorbance was recorded using a Lambda 35 UV-visible spectrophotometer (Perkin Elmer). The extinction coefficient of Co^II^Mg^II^GTP-CobW (ε_339 nm_ = 2,800 ± 100 cm^−1^ M^−1^, average ± s.d of n=3 independent titrations) was determined from absorbance at saturating metal concentrations (Supplementary Fig. 3d). Extinction coefficients of related complexes Co^II^Mg^II^GMPPNP-CobW, Co^II^Mg^II^GTPγS-CobW, Co^II^_2_GTP-CobW, Co^II^_2_GMPPNP-CobW and Co^II^_2_GTPγS-CobW were similarly determined (Figs. 1-2, Supplementary Fig. 3): within experimental error, all produced the same extinction coefficient as for Co^II^Mg^II^GTP-CobW thus ε_339 nm_ = 2,800 cm^−1^ M^−1^ was assigned to all species. Gel-filtration chromatography experiments were performed by incubating CobW (10 μM) and Co^II^ (30 μM) for 30 minutes with or without cofactor GMPPNP (30 μM) then applying 0.5 mL to a PD-10 Sephadex G-25 gel-filtration column (GE Healthcare). Eluted fractions (0.5 mL) were analysed for cobalt by ICP-MS and for protein by Bradford assay.

### Determination of metal affinities via ligand competition

Ligand competition experiments were conducted in an anaerobic chamber in deoxygenated, chelex-treated 10 mM HEPES pH 7.0, 100 mM NaCl, 400 mM KCl, except where high concentrations (≥ 1 mM) of competing ligand were employed, where 50 mM HEPES was used to maintain buffered pH 7.0. Absorbance was recorded using a Lambda 35 UV-visible spectrophotometer (Perkin Elmer). Fluorescence spectra were recorded using a Cary Eclipse fluorescence spectrophotometer (Agilent). Affinities were determined at a range of different competing conditions (by varying the competing ligand and/or the protein:ligand ratio) to ensure reliability: details are documented in Supplementary Table 2. Scripts used for data fitting (using Dynafit^57^) are provided in Supplementary Note 3. The effect of Mg^II^ (2.7 mM) on apparent dissociation constants of ligand standards (EGTA, NTA, Fura-2, Mf2 and quin-2) was calculated to be insignificant under the conditions of competition experiments (Supplementary Table 1). For probes with undefined Mg^II^ affinities (Tar, Bca) control experiments confirmed that addition of Mg^II^ (2.7 mM) had negligible effect on competition experiments (Supplementary Figs. 10d and 12). Thus, Mg^II^ was not incorporated into the curve-fitting models.

For determination of weaker (*K*_D_ >10 nM) Co^II^ binding affinities (CobW and CobW-Mg^II^GDP), Co^II^ was titrated into a solution of fura-2 (quantified by ε_363 nm_ = 28,000 cm^−1^ M^−1^)^58^ and CobW in the presence or absence of cofactors (Mg^II^ and GDP) and fluorescence emission (λ_ex_ = 360 nm; λ_max_ ~ 505 nm) was recorded at equilibrium. Co^II^-dependent fluorescence quenching of fura-2 was used to determine Co^II^ speciation. For determination of Co^II^ binding affinities tighter than 10 nM (CobW-Mg^II^GMPPNP, CobW-Mg^II^GTPγS and CobW-Mg^II^GTP), Co^II^ was titrated into a solution containing CobW, competing ligand (EGTA or NTA), Mg^II^ and nucleotide (GMPPNP, GTPγS or GTP). UV-visible absorbance (relative to metal-free solution) was recorded at equilibrium to determine Co^II^ speciation (ε_339 nm_ = 2,800 cm^−1^ M^−1^ for Co^II^-bound proteins). Data were fit using Dyanfit^57^ to models describing 1:1 binding stoichiometry for Co^II^:protein and 1:1 binding stoichiometry for Co^II^:ligand (ligand = Fura-2, EGTA or NTA). Ligand dissociation constants at pH 7.0: Fura-2 *K*_Co(II)_ = 8.6 × 10^−9^ M (ref.^59^); EGTA *K*_Co(II)_ = 7.9 ×10^−9^ M (ref.^60^); NTA *K*_Co(II)_ = 2.2 ×10^−8^ M (ref.^60^).

Fe^II^ was titrated into a solution of Tar (16 μM), Mg^II^ (2.7 mM) and GTP (500 μM) in the absence or presence of CobW (50 μM) and UV-visible absorbance recorded at equilibrium to define Fe^II^ speciation (Fe^II^Tar_2_ ε_720_ = 19,560 cm^−1^ M^−1^ under experimental conditions, Supplementary Fig 8a). Data were fit in Dynafit^57^ to a model describing 1:1 binding stoichiometry for Fe^II^:protein and 1:2 binding stoichiometry for Fe^II^:Tar using β_2,Fe(II)_ = 4.0 × 10^13^ M^−2^ for Tar at pH 7.0 (ref.^61^). Experimental data were compared to simulated fits with defined protein *K*_Fe(II)_ = 10^−6^ M, 10^−7^ M, allowing limiting *K*_D_ ≥ 10^−6^ M for CobW-Mg^II^GTP to be determined. Tar stock concentrations were quantified using ε_470 nm_ = 24,800 cm^−1^ M^−1^ (reported value at pH 7.0^61^) and verified by titration with metal stocks (Fe^II^ or Ni^II^, quantified by ICP-MS).

Ni^II^ was titrated into a solution of Tar (20 μM), CobW (10 – 30 μM), Mg^II^ (2.7 mM) and GTP (100 – 300 μM) and UV-visible absorbance recorded at equilibrium to determine Ni^II^ speciation (Ni^II^Tar_2_ Δε_535 nm_ = 3.8 (±0.1) × 10^4^ cm^−1^ M^−1^ relative to ligand only solution; Supplementary Fig. 10a). Tar stock concentrations were quantified as above. Data were fit using Dynafit^57^ to a model describing 1:1 stoichiometry Ni^II^:protein and 1:2 stoichiometry Ni^II^:Tar. β_2,Ni(II)_ = 4.3 (±0.6) × 10^15^ M^−2^ for Tar at pH 7.0 was independently determined by preparing a series of solutions of NiTar_2_ ([Ni^II^] = 15 μM, [Tar] = 36 μM) with varying EGTA concentrations (0 – 400 μM) and measuring UV-visible absorbance at equilibrium (following 1-2h incubation). EGTA *K*_Ni(II)_= 5.0 ×10^−10^ M at pH 7.0 (ref.^60^). Data were fit to equation (5)^62^ using Kaleidagraph (Synergy Software).

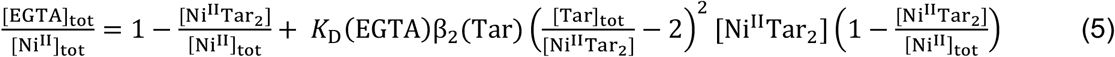

CuSO_4_ was titrated into a solution of Bca (1.0 mM), CobW (10 – 30 μM), Mg^II^ (2.7 mM), GTP (100 – 300 μM) and reductant NH_2_OH (1.0 mM) which quantitatively reduces Cu^II^ to Cu^I^ in the presence of a strong Cu^I^ ligand (*eg* Bca: β_2,Cu(I)_ = 1.6 × 10^17^ M^−2^ (ref.^60^)). UV-visible absorbance was recorded at equilibrium to define Cu^I^ speciation (Cu^I^Bca_2_ ε_562_ = 7,900 cm^−1^ M^−1^ (ref.^60^)) and data were fit using Dynafit^57^ to a model describing 1:1 stoichiometry Cu^I^:protein and 1:2 stoichiometry Cu^I^:Bca.

Zn^II^ was titrated into a solution containing quin-2 (10 μM), CobW (10 μM), Mg^II^ (2.7 mM) and GTP (50 μM) and UV-visible absorbance recorded at equilibrium. Quin-2 was quantified using ε_261 nm_= 37,000 cm^−1^ M^−1^ (ref.^63^). *K*_Zn(II)_ for CobW-Mg^II^GTP was beyond the range of this experiment (significantly tighter than quin-2) and only a limiting affinity was determined (*K*_Zn(II)_ < 10^−12^ M).

### Determination of Zn^II^ affinity of Mg^II^GTP-CobW via inter-metal competition

Solutions containing CobW (17.9 – 20.4 μM), Mg^II^ (2.7 mM), GTP (200 μM) and ligand NTA (0.4 – 4.0 mM) were titrated with Co^II^ (0.3 – 3.0 mM) and ZnSO_4_ (15.3 – 25.5 μM) and UV-visible absorbance was recorded at equilibrium to determine Co^II^ occupancy of CobW (ε_339 nm_ = 2,800 cm^−1^ M^−1^ for Co^II^Mg^II^GTP-CobW). Details of individual experiments are in Supplementary Table 4. The total concentration of Co^II^ and Zn^II^ in each solution was limiting, such that both metals were buffered by ligand NTA. Metal speciation was determined from the mass balance relationships given in equations (6–8) (cofactors Mg^II^GTP omitted for clarity). Thus, *K*_Zn(II)_ for CobW-Mg^II^GTP was calculated from the exchange equilibria (*K*_ex_) in Fig. 5a, relative to known *K*_Co(II)_ for the protein (Supplementary Table 3) and ligand dissociation constants (NTA *K*_Zn(II)_ = 1.18 × 10^−8^ M, *K*_Co(II)_ = 2.24 × 10^−8^ M (ref.^60^)). These calculations are valid given that [M]_free_ ≪ [M]_tot_ (M = Co^II^ or Zn^II^, buffered by excess NTA), the concentration of non-metalated protein is negligible (Supplementary Fig. 13) and potential ternary complexes involving metal, protein and NTA are transient species only with insignificant concentration at thermodynamic equilibrium (varying ratios of buffered metals, [Co^II^NTA]/[Zn^II^NTA], were used to confirm consistency of *K*_D_ values at multiple equilibria; see Fig. 5 and Supplementary Table 4).

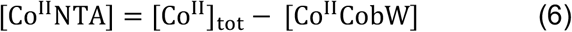

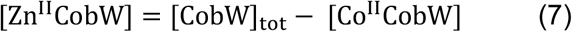

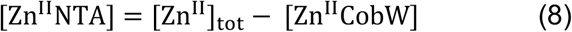

### GTPase activity assays

CobW (20 – 50 μM) was incubated with Co^II^ (0.9 equivalents Co^II^:protein) and GTP (200 μM) in an anaerobic chamber in N_2_-purged, chelex-treated 10 mM HEPES pH 7.0, 100 mM NaCl, 400 mM KCl. Aliquots of solution taken at various time intervals (0 – 390 mins) were loaded to a 5mL HiTrap Q HP column (GE Healthcare) equilibrated in buffer (20 mM HEPES pH 7.0, 100 mM NaCl) and eluted with a linear NaCl gradient (100 – 500 mM NaCl). Nucleotides were detected by UV absorbance (254 nm or 280 nm) and the ratio of GTP:GDP in each sample was calculated by integration of the respective peak areas.

### Growth of *E. coli\** strains

*E. coli\** strains used in this work are derived from *E. coli* MG1655 (DE3) engineered to contain the set of B_12_ biosynthesis genes from *R. capsulatus* (described in refs.^64,65^), except *cobG* and *cobE* are *Brucella melitensis* homologs (described in ref.^39^). Chromosomally-integrated B_12_ biosynthesis genes are IPTG-inducible under the control of the T7 promoter but in the current experiments IPTG was not added to cell cultures to avoid potential disruptions of cellular metal homeostasis caused by over-production of metalloproteins. All cultures and media were prepared in plasticware or acid-washed glassware to minimize trace metal contamination. LB medium was inoculated with overnight culture of *E. coli\** (OD_600 nm_ = 0.025) and incubated at 37°C with shaking until OD_600 nm_ reached ~ 0.2. Aliquots (5 mL or 50 mL) of this culture were treated with sterile Co^II^, H_2_O, EDTA or Zn^II^ (100 × concentrated stocks) to reach final concentrations as specified in figure legends (Figs. 7b, 8b and Supplementary Figs. 15 16a,b,d and 17c) and incubated under the same conditions for a further 1-4h. Samples used for RNA extraction were taken 1h after treatment. Samples for B_12_ quantification and OD_600 nm_ readings were taken 4h after treatment to ensure detectable corrinoid production.

### Determination of transcript abundance in *E. coli\**

Aliquots (1 mL) of *E. coli\** culture from each growth condition were stabilised in RNAProtect Bacteria Reagent (2 mL; Qiagen) and cells pellets were frozen at −80°C prior to processing. RNA was extracted using an RNeasy Mini Kit (Qiagen) as described by the manufacturer. RNA was quantified by absorbance at 260 nm and treated with DNAse I (2.5 U/μL; Fermentas). cDNA was generated using the ImProm-II Reverse Transcriptase System (Promega) with 300 ng RNA per reaction, and control reactions without reverse transcriptase were conducted in parallel. Transcript abundance was determined using primers 3 and 4 for *rcnA*, 5 and 6 for *zntA*, 7 and 8 for *znuA*, 9 and 10 for *rpoD*, each pair designed to amplify ~110 bp fragment (Supplementary Table 6). Quantitative PCR analysis was carried out in 20 μL reactions using 5 ng cDNA, 0.8 μM of each appropriate primer and PowerUp SYBR Green Master Mix (Thermo Fisher Scientific). Three technical replicates of each sample (*ie* biological replicate) were analysed using a Rotor-Gene Q 2plex (Qiagen), plus control reactions without cDNA template for each primer pair. The fold change, relative to the mean of the control condition for each sensor, was calculated using the 2^−ΔΔCT^ method^66^, with *rpoD* as the reference gene. *C*_q_ values were calculated with LinRegPCR after correcting for amplicon efficiency^67^.

### Determination of intracellular available ΔG_Co(II)_ under bespoke conditions

Fractional responses (*θ*_D_) of RcnR at bespoke growth conditions were calculated from transcript abundance of *rcnA* via equation (9):

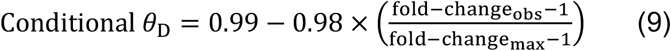

where fold-change_obs_ is the observed fold-change in *rcnA* transcript abundance at the bespoke condition and fold-change_max_ is the maximum fold-change in *rcnA* transcript abundance at the calibration limit (corresponding to maximum abundance); all fold-changes were determined relative to the defined control condition (untreated LB) corresponding to minimum *rcnA* transcript abundance (see Supplementary Fig. 15c). Equation (9) defines maximum and minimum transcript abundances as corresponding to *θ*_D_ of 0.01 and 0.99, respectively (see Fig. 7a), and assumes a linear relationship between change in *θ*_D_ and change in transcript abundance.

The intracellular available [Co^II^] concentration corresponding to each RcnR *θ*_D_ was calculated as described in ref.^27^ using properties determined for *Salmonella* RcnR to calculate the Co^II^-dependent response of *E. coli* RcnR (93% sequence identity). The intracellular available Δ*G*_Co(II)_ for each condition was calculated using equation (10), where [Co^II^] is the intracellular available Co^II^ concentration, R (gas constant) = 8.314 × 10^−3^ kJ K^−1^ mol^−1^ and T (temperature) = 298.15 K (see Supplementary Note 1).

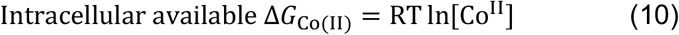

### Estimation of intracellular available ΔG_Zn(II)_ in LB media

Fractional responses (*θ*_D_) of Zur and ZntR in LB media were calculated from transcript abundance of *znuA* and *zntA*, via equations (9) and (11), respectively:

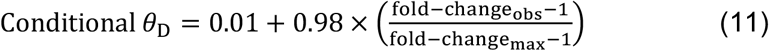

where fold-change_obs_ is the observed fold-change in transcript abundance in LB and fold-change_max_ is the maximum fold-change in transcript abundance at the calibration limit (corresponding to maximum abundance); all fold-changes were determined relative to defined control conditions corresponding to minimum transcript abundance (see Supplementary Fig 16a,b). Equation (11) defines maximum and minimum transcript abundances as corresponding to *θ*_D_ of 0.99 and 0.01, respectively, and assumes a linear relationship between change in *θ*_D_ and change in transcript abundance.

The intracellular available [Zn^II^] concentration corresponding to each *θ*_D_ was calculated as described in ref.^27^ using properties determined for *Salmonella* homologs to calculate the Zn^II^-dependent responses of *E. coli* ZntR and Zur (both > 92% sequence identity). The intracellular available Δ*G*_Zn(II)_ was calculated using equation (12), where [Zn^II^] is the intracellular available Zn^II^ concentration, R (gas constant) = 8.314 × 10^−3^ kJ K^−1^ mol^−1^ and T (temperature) = 298.15 K (see Supplementary Note 1).

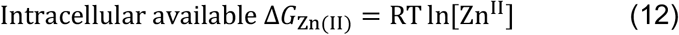

### Quantification of vitamin B_12_ in *E. coli\** cultures

Aliquots (20 mL) of *E. coli\** culture from each growth condition were taken, and cell pellets frozen at −20°C. To quantify corrin production (assumed to be predominantly B_12_, since *E. coli\** contains genes for the complete pathway), *E. coli\** pellets were thawed, resuspended in H_2_O (0.2 mL), boiled for 15 min (95°C) and centrifuged to remove cell debris. An aliquot (10 μL) of each supernatant was applied to bioassay plates containing *Salmonella typhimurium* AR2680 (*ΔmetE, ΔcbiB*) prepared as previously reported^37^ and incubated at 37°C overnight. Plates were imaged together with a 1 cm^2^ reference area on black background (see example in Supplementary Data 2) using a Gel-Doc XR + gel documentation system (BioRad). Images were analysed in MATLAB using the code in Supplementary Note 2 to determine the growth area (in cm^2^) of each sample. A calibration curve relating growth areas to B_12_ concentration was generated using B_12_ standards (cyanocobalamin; 1 – 100 nM; quantified by A_360 nm_ = 27,500 cm^−1^ M^−1^ at pH 10 (ref.^68^)) in parallel with *E. coli\** lysates, using the same batch of bioassay plates (Supplementary Fig. 17a-b). To determine the number of cells in each sample, solutions of *E. coli\** at varying cell densities (OD_600 nm_ = 0.2 – 0.9) were prepared, serially diluted (2000-fold), and the number of cells per mL quantified using a CASY® cell counter. The resulting correlation factor (4.4 ± 0.1 × 10^8^ cells mL^−1^ OD_600 nm-1_) was used to convert OD_600 nm_ to cell number (Supplementary Fig. 17c,d).

## Acknowledgements

This work was supported by a COFUND European Union/Durham University Junior Research Fellowship under EU grant agreement 609412 (T.R.Y.), a Royal Commission for the Exhibition of 1851 Research Fellowship (T.R.Y.), UKRI Future Leaders Fellowship MR/T019891/1 (R.J.M.), Biotechnology and Biological Sciences Research Council awards BB/S009787/1, BB/J017787/1, BB/R002118/1, BB/S002197/1, and Royal Society award INF\R2\180062. We thank Andrew Foster and Peter Chivers (Durham University, UK) and Arthur Glasfeld (Reed College, USA) for constructive scientific discussions.

## Author Contributions

T.R.Y. conducted the *in vitro* metal-binding experiments, GTP-hydrolysis assays, *in vivo* gene expression experiments and B_12_-production experiments. T.R.Y. and M.A.M. developed experimental protocols for determining metal sensor responses by qPCR. M.A.M. derived equations for the metalation calculator and produced the spreadsheet. R.J.M. and D.O. generated the MATLAB code for analysis of B_12_ bioassays. E.D. generated the CobW expression plasmid. E.D. and M.J.W. donated the B_12_-producing *E. Coli\** strains and advised on B_12_ biochemistry. E.D., M.J.W., and T.R.Y. co-designed the B_12_-production experiments. T.R.Y. and N.J.R. drafted the manuscript and, in conjunction with M.A.M and D.O., interpreted the significance of the data. T.R.Y. and N.J.R. had overall responsibility for the design and management of the project. All authors reviewed the results and edited and approved the final version of the manuscript.

## Competing Financial Interests

The authors declare no competing financial interests.

## Notes

### Competing Interest Statement

The authors have declared no competing interest.

