## Supplementary material for "Calculating metalation in cells reveals CobW acquires Co^II^ for vitamin B_12_ biosynthesis upon binding nucleotide": CobW_SI.pdf

### Table of contents

**Supplementary Table 1.** Calculated effect of 2.7 mM $Mg^{II}$ on apparent ligand dissociation constants at pH 7.0

**Supplementary Table 2.** Conditions of CobW ligand competition experiments

**Supplementary Table 3.** Metal affinities (and corresponding free energies) of CobW

**Supplementary Table 4.** Calculated stability constants ($K_D$ and $\Delta G$ values) for $Zn^{II}$-binding to $Mg^{II}$GTP-CobW under various conditions, from $Zn^{II}$-$Co^{II}$ inter-metal competition experiments (Fig. 5)

**Supplementary Table 5.** Calculations of conditional intracellular available [$Co^{II}$] (and corresponding $\Delta G_{Co(II)}$) for *E. coli** cultures in $Co^{II}$-supplemented LB media

**Supplementary Table 6.** Oligonucleotides used in this work

**Supplementary Figure 1.** Full gel image for Fig. 1b

**Supplementary Figure 2.** Visible absorbance features from Fig. 1g

**Supplementary Figure 3.** Different GTP analogues assemble similar metal-binding sites in CobW

**Supplementary Figure 4.** $K_{Co(II)}$ quantification for CobW, in the absence or presence of nucleotides, using a variety of competing conditions

**Supplementary Figure 5**. $K_{Co(II)}$ quantification for $Mg^{II}$GTP-CobW at varying nucleotide concentrations

**Supplementary Figure 6.** CobW-catalysed GTP hydrolysis is also slow at 4:1 ratio GTP:CobW

**Supplementary Figure 7.** $Mg^{II}$GDP-CobW has a similar $Co^{II}$ site to $Mg^{II}$GTP-CobW but with weaker $K_{Co(II)}$

**Supplementary Figure 8.** Further details of $K_{Fe(II)}$ quantification for $Mg^{II}$GTP-CobW using Tar

**Supplementary Figure 9.** Further details of $K_{Ni(II)}$ quantification for $Mg^{II}$GTP-CobW

**Supplementary Figure 10.** Determination of $\beta_2$ for $Ni^{II}Tar_2$ at pH 7.0

**Supplementary Figure 11.** Further details of $K_{Cu(I)}$ quantification for $Mg^{II}$GTP-CobW

**Supplementary Figure 12.** Effect of $Mg^{II}$ on $Cu^{I}$-competition experiments using Bca

**Supplementary Figure 13.** $Mg^{II}$GTP-CobW is metal-saturated in Fig. 5 inter-metal competition experiments

**Supplementary Figure 14.** The accessible range of intracellular available free energies for metal-binding as sensors shift from 1 – 99% of their responses

**Supplementary Figure 15.** Calibration of maximum and minimum *rcnA* responses

**Supplementary Figure 16.** Estimation of $Zn^{II}$ availability within *E. coli** cells in defined growth conditions (conditional cells)

**Supplementary Figure 17.** Quantification of vitamin $B_{12}$ in *E. coli**

**Supplementary Table 1.** Calculated effect of 2.7 mM $Mg^{II}$ on apparent ligand dissociation constants at pH 7.0

| **Ligand (L)** | **$K_{Mg(II)}$ (M)** [a] | **$K_{Mg(II)}$ (M) (pH 7.0)** [b] | **α coefficient (2.7 mM $Mg^{II}$, pH 7.0)** [c] |
|---|---|---|---|
| EGTA | $6.2 \times 10^{-6}$ [d] | $9.7 \times 10^{-2}$ [e] | 0.97 |
| NTA | $3.9 \times 10^{-6}$ [d] | $2.1 \times 10^{-3}$ [e] | 0.44 |
| Fura-2 | | $9.8 \times 10^{-3}$ [f] | 0.78 |
| Mag-fura-2 | | $2.7 \times 10^{-3}$ [g] | 0.50 |
| quin-2 | | $2.0 \times 10^{-3}$ [h] | 0.43 |

[a] Absolute dissociation constants ($K_D$) for $Mg^{II}L$ complexes (0.1 M ionic strength)

[b] Conditional dissociation constants ($K_D$) for $Mg^{II}L$ complexes at pH 7.0 (0.1 M ionic strength)

[c] $\alpha = 1/(1+[Mg]/K_{Mg(II)})$ describes the effect of 2.7 mM $Mg^{II}$ on the apparent ligand affinity for other metals at pH 7.0: $K_D' = K_D/\alpha$, where $K_D$ is the ligand dissociation constant at pH 7.0 in the absence of $Mg^{II}$ and $K_D'$ is the apparent ligand dissociation constant at pH 7.0 in the presence of 2.7 mM $Mg^{II}$.

[d] From ref [1]

[e] Calculated from absolute dissociation constant as described in ref [2]

[f] At pH 7.2, from ref [3]

[g] At pH 7.4, from ref [4]

[h] From refs[3,5]

**Supplementary Table 2.** Conditions of CobW ligand competition experiments [a]

| **Metal (*M*)** | **Nucleotide** [b] | **Competing ligand (*L*)** [c] | **[CobW]$_{tot}$ (μM)** | **[*L*]$_{tot}$ (μM)** | **[*M*]$_{max}$ (μM)** | **Species detected (signal)** [d] | ***K*$_D$ (M)** | **Figure ref.** |
|---|---|---|---|---|---|---|---|---|
| $Co^{II}$ | None | fura-2 | 34 | 10 | 80 | $Co^{II}$fura-2 | $1.1 \times 10^{-7}$ | S4f |
| | | | 37 | 10 | 82 | ($F_{550\ nm}$) | $2.7 \times 10^{-7}$ | 3a |
| | | | 50 | 10 | 70 | | $3.9 \times 10^{-7}$ | S4g |
| | GDP | | 10 | 10 | 22 | | $8.8 \times 10^{-8}$ | S4i |
| | | | 20 | 8.1 | 46 | | $1.0 \times 10^{-7}$ | 3b |
| | | | 30 | 8.1 | 50 | | $1.1 \times 10^{-7}$ | S4h |
| | GMPPNP | EGTA | 20 | 20 | 40 | $Co^{II}$CobW | $2.3 \times 10^{-9}$ | S4a |
| | | | 20 | 40 | 84 | ($A_{339\ nm}$) | $3.2 \times 10^{-9}$ | 2a |
| | | | 20 | 40 | 90 | | $2.6 \times 10^{-9}$ | S4b |
| | GTPγS | | 20 | 200 | 222 | | $1.1 \times 10^{-10}$ | S4c |
| | | | 20 | 1000 | 40 | | $2.8 \times 10^{-10}$ | S4d |
| | | | 20 | 2000 | 40 | | $1.2 \times 10^{-10}$ | S4e |
| | GTP | | 20 | 100 | 170 | | nd [e] | S4k |
| | | | 20 | 500 | 122 | | $4.0 \times 10^{-11}$ | S4l |
| | | | 18 | 2000 | 34 | | $2.7 \times 10^{-11}$ | 3c |
| | | NTA | 18 | 4000 | 34 | | $2.2 \times 10^{-11}$ | S4m |
| $Fe^{II}$ | | Tar | 50 | 16 | 80 | $Fe^{II}(Tar)_2$ | $< 10^{-6}$ [f] | 4a/S8b |
| | | | 50 | 16 | 60 | ($A_{720\ nm}$) | $< 10^{-6}$ [f] | S8b |
| | | | 50 | 16 | 60 | | $< 10^{-6}$ [f] | S8b |
| $Ni^{II}$ | | | 10 | 20 | 40 | $Ni^{II}(Tar)_2$ | $3.8 \times 10^{-10}$ | S9b |
| | | | 20 | 20 | 38 | ($A_{535\ nm}$) | $1.9 \times 10^{-9}$ | S9c |
| | | | 30 | 20 | 42 | | $6.9 \times 10^{-10}$ | 4b |
| $Cu^{I}$ | | Bca | 10 | 1000 | 34 | $Cu^{I}(Bca)_2$ | $2.6 \times 10^{-16}$ | S11c |
| | | | 20 | 1000 | 34 | ($A_{562\ nm}$) | $1.2 \times 10^{-16}$ | 4c |
| | | | 30 | 1000 | 32 | | $3.4 \times 10^{-16}$ | S11d |

[a] All experiments conducted anaerobically in $N_2$-purged, chelex-treated HEPES (10 – 50 mM) pH 7.0, 100 mM NaCl, 400 mM KCl

[b] GTP and GDP were supplied in ~10-fold excess of protein concentration and non-hydrolysable analogues (GMPPNP and GTPγS) were supplied in ~3-fold excess of protein concentration (as specified in figure legends). All affinity determinations in presence of nucleotides included 2.7 mM $Mg^{II}$.

[c] fura-2, EGTA and NTA form 1:1 $Co^{II}$:ligand complexes: at pH 7.0 fura-2 $K_{Co(II)} = 8.6 \times 10^{-9}$ M (ref.[6]); EGTA $K_{Co(II)} = 7.9 \times 10^{-9}$ M (ref.[2]); and NTA $K_{Co(II)} = 2.2 \times 10^{-8}$ M (ref.[2]). Tar forms 1:2 metal:ligand complexes at pH 7.0: $\beta_{2,Fe(II)} = 4.0 \times 10^{13}$ $M^{-2}$ (ref.[7]); $\beta_{2,Ni(II)} = 4.3 \times 10^{15}$ $M^{-2}$ (Supplementary Fig. 10). Bca forms a 1:2 $Cu^{I}$:ligand complex with $\beta_2 = 1.6 \times 10^{17}$ $M^{-2}$ (ref.[8])

[d] For cases where titration end-point was reached, probe responses were defined as variable parameters in Dynafit model (final fitted responses were all consistent with known extinction coefficients ± experimental error). For cases where titration end-point was not reached, probe responses were fixed to known extinction coefficients of metal-bound species in Dynafit models: $Co^{II}Mg^{II}$GTP*-CobW (where GTP* = GTP, GTPγS, GMPPNP) $\varepsilon_{339\ nm} = 2{,}800$ $cm^{-1}$ $M^{-1}$ (Fig. 2 and Supplementary Fig. 3), $Fe^{II}(Tar)_2$ $\varepsilon_{720\ nm} = 19{,}560$ $cm^{-1}$ $M^{-1}$ (Supplementary Fig. 8a), $Ni^{II}(Tar)_2$ $\Delta\varepsilon_{535\ nm} = 38{,}000$ $cm^{-1}$ $M^{-1}$ wrt ligand only (Supplementary Fig. 10a), $Cu^{I}(Bca)_2$ $\varepsilon_{562\ nm} = 7{,}900$ $cm^{-1}$ $M^{-1}$ (ref.[2]).

[e] Insufficient [EGTA] for effective competition with $Mg^{II}$GTP-CobW, data not used for $K_D$ determination.

[f] Negligible competition between probe Tar and protein for $Fe^{II}$ binding, only a limiting affinity was determined (n=3 independent experiments, same conditions).

**Supplementary Table 3.** Metal affinities (and corresponding free energies) of CobW

| **Species** | **Metal** | **$K_D$ (M)** [a] | **$\Delta G_{MP}$ (kJ mol$^{-1}$)** |
|---|---|---|---|
| CobW | $Co^{II}$ | 2.5 (±1.1) ×$10^{-7}$ | -37.7 ± 1.2 |
| CobW-$Mg^{II}$GDP | | 1.0 (±0.1) ×$10^{-7}$ | -39.9 ± 0.3 |
| CobW-$Mg^{II}$GMPPNP | | 2.7 (±0.4) ×$10^{-9}$ | -48.9 ± 0.4 |
| CobW-$Mg^{II}$GTPγS | | 1.7 (±0.8) ×$10^{-10}$ | -55.7 ± 1.2 |
| CobW-$Mg^{II}$GTP | | 3.0 (±0.8) ×$10^{-11}$ | -60.1 ± 0.7 |
| | $Fe^{II}$ | > $10^{-6}$ [b] | > -34.2 [b] |
| | $Ni^{II}$ | 9.8 (±6.5) ×$10^{-10}$ | -51.4 ± 1.7 |
| | $Cu^{I}$ | 2.4 (±0.9) ×$10^{-16}$ [c] | -89.2 ± 1.0 [c] |
| | $Zn^{II}$ | 1.9 (±0.6) ×$10^{-13}$ | -72.6 ± 0.8 |

[a]Data are mean ±s.d. of n=3 independent experiments conducted at a range of competing conditions (see Supplementary Table 2), except $K_{Zn(II)}$ of CobW-$Mg^{II}$GTP is mean ±s.d. of n=4 independent experiments (see Supplementary Table 4).
[b]Probe Tar outcompetes CobW-$Mg^{II}$GTP in $Fe^{II}$-binding experiments (see Fig. 4a), limiting $K_D$ value only.
[c]Assuming only tightest affinity site binds $Cu^{I}$ under experimental conditions (one-site model). A model with two sites of equivalent affinity fits to marginally-weaker $K_D$ values (6.1 (±2.0) ×$10^{-16}$ M).

**Supplementary Table 4.** Calculated stability constants ($K_D$ and Δ*G* values) for $Zn^{II}$-binding to $Mg^{II}$GTP-CobW under various conditions, from $Zn^{II}$-$Co^{II}$ inter-metal competition experiments (Fig. 5)

| **Total concentrations added** | | | | **Calculated ratios at equilibrium** | | ***In vitro* available $\Delta G_M$ (kJ mol$^{-1}$)** $^a$ | | **CobW $K_{Zn(II)}$ (M)** $^b$ | **$\Delta G_{Zn(II)\text{-}CobW}$ (kJ mol$^{-1}$)** $^b$ |
|---|---|---|---|---|---|---|---|---|---|
| **$[CobW]_{tot}$ (μM)** | **$[NTA]_{tot}$ (μM)** | **$[Co^{II}]_{tot}$ (μM)** | **$[Zn^{II}]_{tot}$ (μM)** | **$[Co^{II}NTA]$/ $[Zn^{II}NTA]$** | **$[Co^{II}CobW]$/ $[Zn^{II}CobW]^b$** | **$\Delta G_{Co(II)}$** | **$\Delta G_{Zn(II)}$** | | |
| 19.7 | 4000 | 3000 | 25.5 | 167 | 1.58 | -40.9 | -55.2 | $1.5 \times 10^{-13}$ | -73.2 |
| 20.4 | 400 | 300 | 15.3 | 67 | 0.99 | -41.2 | -53.2 | $2.4 \times 10^{-13}$ | -72.0 |
| 18.2 | 3710 | 302 | 23.2 | 28 | 0.45 | -49.7 | -59.5 | $2.6 \times 10^{-13}$ | -71.8 |
| 17.9 | 3710 | 302 | 23.2 | 34 | 0.24 | -49.7 | -60.0 | $1.1 \times 10^{-13}$ | -73.9 |

$^a$*in vitro* available $\Delta G_{Co(II)}$ and $\Delta G_{Zn(II)}$ determined at equilibrium; *ie* $\Delta G_{Co(II)}$ = RT ln$[Co^{II}]$ = RT $K_{Co(II)}$(NTA)×($[Co^{II}$-NTA]/[NTA])

$^b$ In the presence of excess $Mg^{II}$ (2.7 mM) and GTP (200 μM), omitted for clarity.

**Supplementary Table 5.** Calculations of conditional intracellular available [$Co^{II}$] (and corresponding $\Delta G_{Co(II)}$) for *E. coli** cultures in $Co^{II}$-supplemented LB media

| **[$Co^{II}$] (μM)** | **$\theta_D$ (RcnR)** | **[$Co^{II}$]$_{buffered}$ (M)** | **$\Delta G_{Co(II)}$ (kJ $mol^{-1}$)** |
|---|---|---|---|
| 0 [a] | 0.99 | $2.4 \times 10^{-11}$ | -60.6 |
| 1 | 0.92 | $2.2 \times 10^{-10}$ | -55.1 |
| 3 | 0.85 | $4.1 \times 10^{-10}$ | -53.6 |
| 10 | 0.57 | $1.9 \times 10^{-9}$ | -49.8 |
| 30 | 0.28 | $6.7 \times 10^{-9}$ | -46.6 |
| 300 [a] | 0.01 | $2.7 \times 10^{-7}$ | -37.5 |

[a] Defined as limits of RcnR dynamic response range (see Supplementary Fig. 15).

**Supplementary Table 6** Oligonucleotides used in this work

| No. | Primer Name | Sequence | Reference |
|---|---|---|---|
| 1 | Rc_cobW_F | 5'-CATCATATGTCCGATCTGACCAAAATCC-3' | This work |
| 2 | Rc_cobW_R | 5'-CATACTAGTCGACATGCATCAGGCGGC-3' | This work |
| 3 | Ec_rcnA_F | 5'-GAACCAGGGCACTCAAAAAC-3' | ref.[9] |
| 4 | Ec_rcnA_R | 5'-TGCGGTATGCGAAATAGTTG-3' | ref.[9] |
| 5 | Ec_zntA_F | 5'-TCCGGCAACGGGTATTAGTG-3' | This work |
| 6 | Ec_zntA_R | 5'-GTTCAGCAACCTGTGCTTCG-3' | This work |
| 7 | Ec_znuA_F | 5'-GTTTGGACTGACACCGCTTG-3' | This work |
| 8 | Ec_znuA_R | 5'-ACGCAGGTTGCTTTTTGCTC-3' | This work |
| 9 | Ec_rpoD_F | 3'-GTGGCTTGCAGTTCCTTGAC-5' | This work |
| 10 | Ec_rpoD_R | 3'-AGGTTGCGTAGGTGGAGAAC-5' | This work |

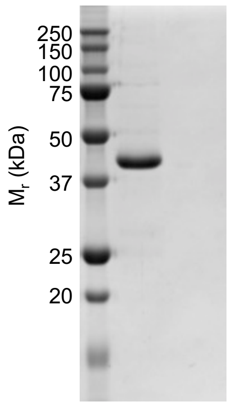


**Supplementary Figure 1. Full gel image for Fig. 1b.**

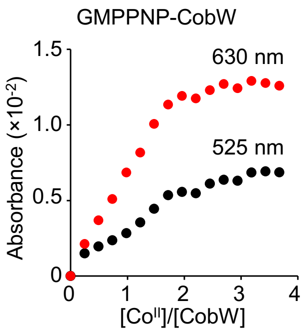


**Supplementary Figure 2. Visible absorbance features from Fig. 1g.** Absorbance of features at 525 nm and 630 nm from Fig. 1g show a linear increase saturating at 2:1 ratio $Co^{II}$:CobW.

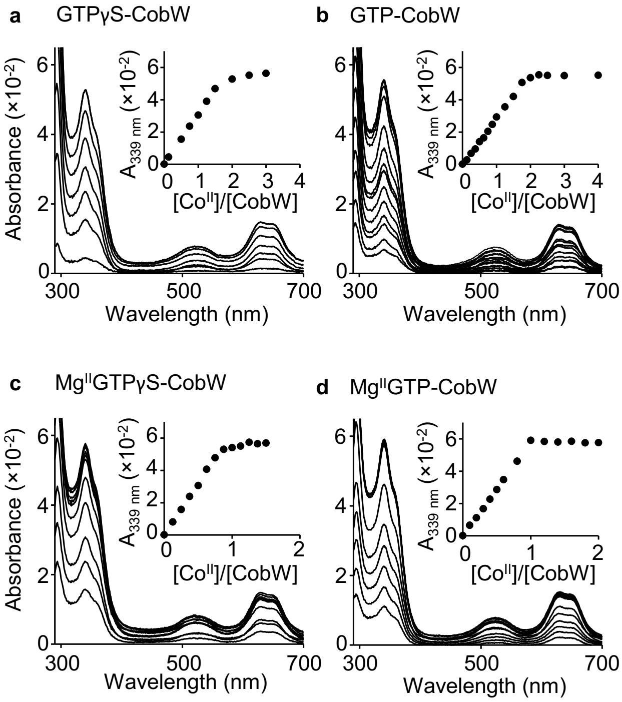


**Supplementary Figure 3. Different GTP analogues assemble similar metal-binding sites in CobW. a-b** Representative apo-subtracted spectra of $Co^{II}$-titrated CobW (20 μM) in the presence of **a** 60 μM GTPγS or **b** 200 μM GTP; features at 339 nm (insets) show linear increase saturating at 2:1 ratio $Co^{II}$:CobW. **c** As in **a** with added $Mg^{II}$ (2.7 mM); feature at 339 nm (inset) shows a linear increase saturating at 1:1 ratio $Co^{II}$:CobW. **d** As in **b** with added $Mg^{II}$ (2.7 mM); feature at 339 nm (inset) shows a linear increase saturating at 1:1 ratio $Co^{II}$:CobW.

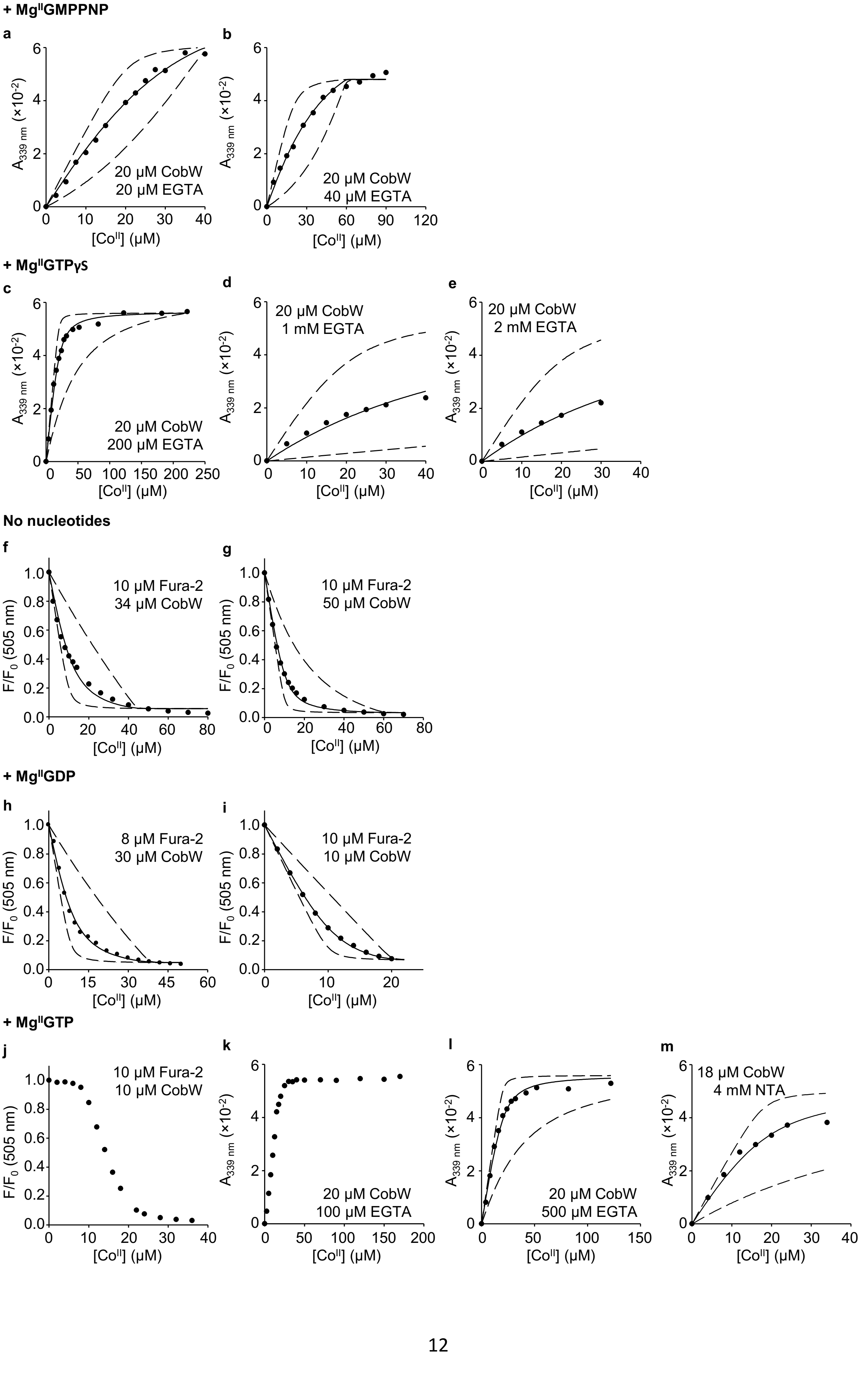
+ Mg^II GMPPNP
a
20 μM CobW
20 μM EGTA
b
20 μM CobW
40 μM EGTA
+ Mg^II GTPγS
c
20 μM CobW
200 μM EGTA
d
20 μM CobW
1 mM EGTA
e
20 μM CobW
2 mM EGTA
No nucleotides
f
10 μM Fura-2
34 μM CobW
g
10 μM Fura-2
50 μM CobW
+ Mg^II GDP
h
8 μM Fura-2
30 μM CobW
i
10 μM Fura-2
10 μM CobW
+ Mg^II GTP
j
10 μM Fura-2
10 μM CobW
k
20 μM CobW
100 μM EGTA
l
20 μM CobW
500 μM EGTA
m
18 μM CobW
4 mM NTA
A339 nm (×10^-2)
F/F0 (505 nm)
[Co^II] (μM)

**Supplementary Figure 4. $K_{Co(II)}$ quantification for CobW, in the absence or presence of nucleotides, using a variety of competing conditions. a-e, k-m** Change in absorbance at 339 nm (wrt $Co^{II}$-free solution) when $Co^{II}$ was titrated into CobW in competition with EGTA or NTA (as specified in each panel) in the presence of $Mg^{II}$ (2.7 mM) and nucleotide (**a-b**) GMPPNP (60 μM), (**c-e**) GTPγS (60 μM), or (**k-m**) GTP (10-fold excess of protein concentration). **f-j** Fluorescence quenching of $Co^{II}$-titrated fura-2 in competition with (**f-g**) CobW alone, (**h-i**) CobW in the presence of $Mg^{II}$ (2.7 mM) and GDP (10-fold excess of protein concentration), (**j**) CobW in the presence of $Mg^{II}$ (2.7 mM) and GTP (100 μM). In **a-i, l-m** solid traces show curve fits of experimental data to models where CobW binds one molar equivalent $Co^{II}$ per protein monomer. Dashed lines show simulated responses for $K_{Co(II)}$ tenfold tighter or weaker than the fitted value. In **j, k** $Co^{II}$ was almost entirely withheld by $Mg^{II}$GTP-CobW, preventing meaningful determination of $K_{Co(II)}$ under these conditions. Further details and fitted $K_{Co(II)}$ values are listed in Supplementary Table 2.

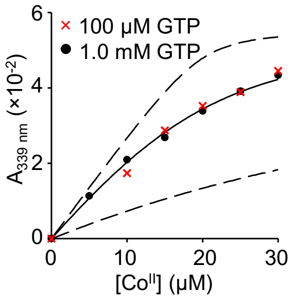


**Supplementary Figure 5. $K_{Co(II)}$ quantification for Mg$^{II}$GTP-CobW at varying nucleotide concentrations.** Change in absorbance at 339 nm (relative to Co$^{II}$-free solution) when Co$^{II}$ was titrated into CobW (20 μM) in competition with EGTA (2.0 mM) in the presence of Mg$^{II}$ (2.7 mM) and GTP (100 μM or 1.0 mM). Experiments performed in 50 mM HEPES pH 7.0, 100 mM NaCl, 400 mM KCl. Solid trace shows representative curve fit of the experimental data (for 1.0 mM GTP dataset) to a model where CobW binds one molar equivalent Co$^{II}$ per protein monomer. Dashed lines show simulated responses for $K_{Co(II)}$ tenfold tighter or weaker than the fitted value. At both nucleotide concentrations the measured $K_{Co(II)}$ ($1.8 \times 10^{-11}$ M and $1.9 \times 10^{-11}$ M) were within experimental error.

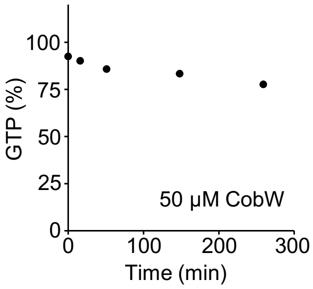


**Supplementary Figure 6. CobW-catalysed GTP hydrolysis is also slow at 4:1 ratio GTP:CobW.** Analysis of GTP hydrolysis when a solution of GTP (200 μM) was incubated with CobW (50 μM), $Mg^{II}$ (2.7 mM) and $Co^{II}$ (45 μM). Nucleotides were separated by anion-exchange and detected by UV absorbance (280 nm).

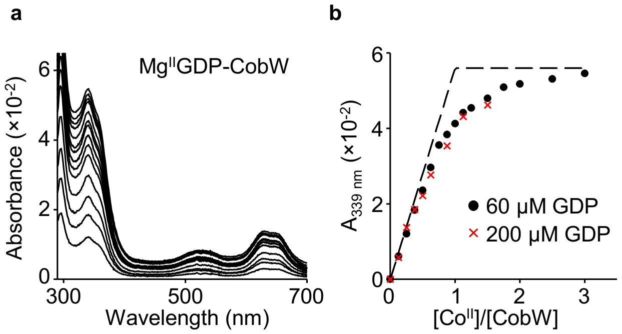


**Supplementary Figure 7. Mg$^{II}$GDP-CobW has a similar Co$^{II}$ site to Mg$^{II}$GTP-CobW but with weaker $K_{Co(II)}$.** **a** Apo-subtracted spectra of Co$^{II}$-titrated CobW (20 µM) in the presence of Mg$^{II}$ (2.7 mM) and GDP (60 µM) showed that Mg$^{II}$GDP-CobW possesses a similar Cys-rich, tetrahedral Co$^{II}$ site to that of Mg$^{II}$GTP-CobW (*cf* Supplementary Fig. 3d). **b** Absorbance feature from (**a**) at 339 nm (black circles) lacks a sharp turning point at 1 equivalent Co$^{II}$:CobW suggesting competition between the tetrahedral site in Mg$^{II}$GDP-CobW and weak-binding solution components (*eg* buffer, salts or alternative Co$^{II}$ sites within CobW itself). An equivalent experiment with 200 µM GDP (red crosses) gave indistinguishable results, showing that nucleotide concentration was not a limiting factor for metal-binding. Dotted trace shows simulated response assuming the same extinction coefficient for Co$^{II}$MgGDP-CobW as for other nucleotide-bound forms (GMPPNP, GTPγS and GTP).

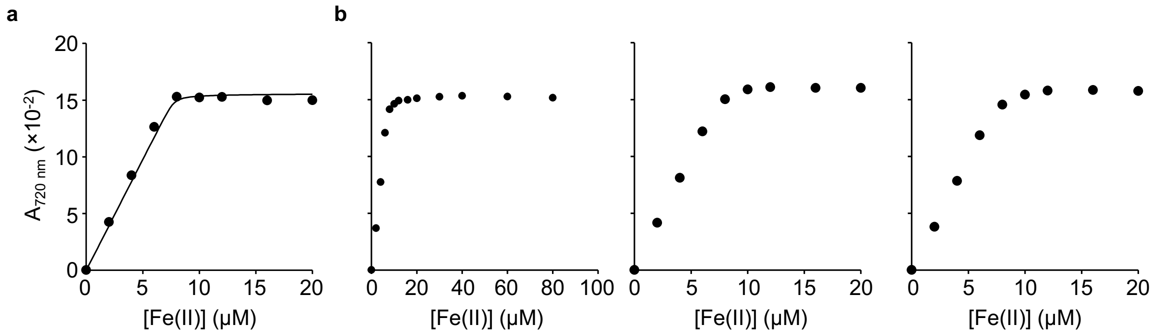


**Supplementary Figure 8. Further details of $K_{Fe(II)}$ quantification for Mg$^{II}$GTP-CobW using Tar.** **a** Titration of Tar (16 μM) with Fe$^{II}$ in buffer only. Solid trace shows fitted extinction coefficient of $\varepsilon_{720\ nm}$ = 1.956 × $10^4$ $cm^{-1}$ $M^{-1}$ assuming stoichiometric Fe$^{II}$Tar$_2$ formation (in agreement with reported value at pH 7.0 (ref.[7])). **b** Source data for Fig. 4a showing entire collected dataset (up to 80 μM added Fe$^{II}$), and two replicated experiments (with up to 20 μM added Fe$^{II}$) confirming that Mg$^{II}$GTP-CobW (50 μM) cannot compete with Tar (16 μM) for binding Fe$^{II}$.

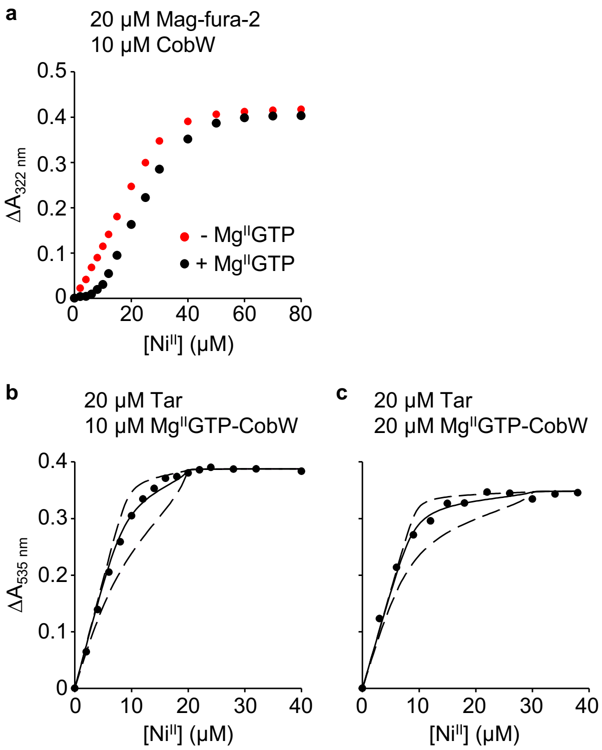


**Supplementary Figure 9. Further details of $K_{Ni(II)}$ quantification for $Mg^{II}$GTP-CobW. a** Absorbance change at 322 nm upon $Ni^{II}$-titration of CobW (10 μM) in competition with mag-fura-2 (Mf2; 20 μM) with or without $Mg^{II}$ (2.7 mM) and GTP (100 μM). In the absence of $Mg^{II}$GTP, CobW binds two $Ni^{II}$ ions with a similar affinity to Mf2 ($K_{Ni(II)} = 5 \times 10^{-8}$ M (ref.[10])); but in the presence of $Mg^{II}$GTP, CobW shows an additional $Ni^{II}$ site which outcompetes Mf2. **b-c** Absorbance change (relative to $Ni^{II}$-free solution) upon $Ni^{II}$-titration of Tar in competition with CobW in the presence of $Mg^{II}$ (2.7 mM) and GTP (100 μM in **b** and 200 μM in **c**). In **b-c** solid traces show curve fits of experimental data to models where CobW binds one molar equivalent $Ni^{II}$ per protein monomer. Dashed lines show simulated responses for $K_{Ni(II)}$ tenfold tighter or weaker than the fitted value. Fitted $K_{Ni(II)}$ values are listed in Supplementary Table 2.

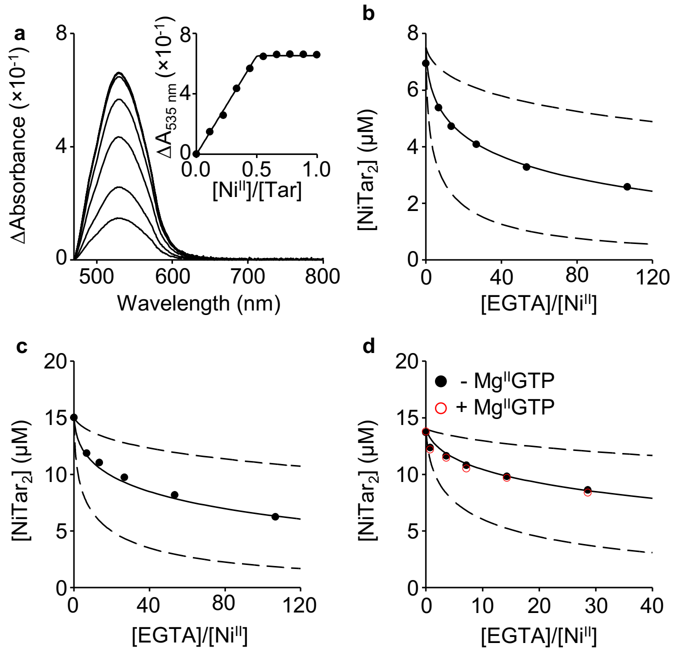


**Supplementary Figure 10. Determination of $\beta_2$ for $Ni^{II}Tar_2$ at pH 7.0. a** Representative (n=3) change in absorbance (relative to metal-free probe) upon $Ni^{II}$-titration of Tar (34 μM). Inset shows linear increase of feature at 535 nm saturating at 1:2 ratio $Ni^{II}$:Tar; and solid trace shows fitted extinction coefficient of ε = 3.8 (±0.1) × $10^4$ $cm^{-1}$ $M^{-1}$ (mean ± s.d. of n=3 experiments) assuming stoichiometric $Ni^{II}Tar_2$ formation. **b-d** Addition of increasing [EGTA] into probe solutions containing (**b**) 7.5 μM $Ni^{II}$, 18 μM Tar; (**c**) 15 μM $Ni^{II}$, 36 μM Tar; and (**d**) 14 μM $Ni^{II}$, 36 μM Tar; shows competition for $Ni^{II}$-binding at equilibrium. Solid traces are curve fitting of experimental data to equation (5) (Methods) resulting in $\beta_2$ = 4.3 (±0.6) × $10^{15}$ $M^{-2}$ for $Ni^{II}Tar_2$ formation at pH 7.0 (mean ± s.d. from n=3 experiments in **b-d**). Dashed lines are models of binding 10-fold tighter or weaker than measured values. Addition of $Mg^{II}$ (2.7 mM) and GTP (500 μM) to solutions in **d** had negligible effect on $Ni^{II}$-binding equilibria.

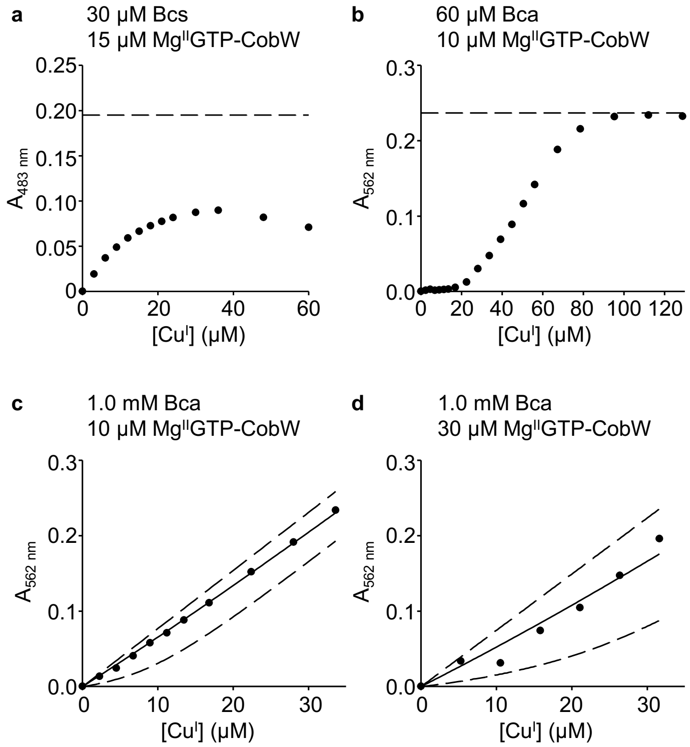


**Supplementary Figure 11. Further details of $K_{Cu(I)}$ quantification for $Mg^{II}$GTP-CobW. a** $Cu^I$-titration into a mixture of Bcs (30 µM), CobW (15 µM), $Mg^{II}$ (2.7 mM) and GTP (200 µM) does not reach theoretical absorbance maximum for complete formation of $Cu^IBcs_2$ (dashed line, $\varepsilon_{483\,nm}$ = 13,000 $cm^{-1}$ $M^{-1}$ (ref.[2])) at saturating [$Cu^I$] which is suggestive of ternary complex formation[11]. **b** Absorbance (562 nm) upon $Cu^I$-titration into a mixture of Bca (60 µM), CobW (10 µM), $Mg^{II}$ (2.7 mM) and GTP (200 µM) shows that the protein binds 2 equivalents $Cu^I$ with tighter affinity than Bca and a further 3 equivalents $Cu^I$ with sufficient affinity to compete with the probe; theoretical absorbance maximum for complete formation of $Cu^IBca_2$ (dashed line, $\varepsilon_{562\,nm}$ = 7,900 $cm^{-1}$ $M^{-1}$ (ref.[2])) is reached upon addition of ~ 80 µM $Cu^I$ (*ie* sufficient to saturate both probe Bca and ~ 5 protein sites). **c-d** Absorbance of $Cu^I$-titrated Bca (1.0 mM) in competition with (**c**) 10 µM CobW or (**d**) 30µM CobW in the presence of $Mg^{II}$ (2.7 mM) and GTP (200 µM). In **c-d** solid traces are curve fits of experimental data to model assuming only the tightest site in CobW binds $Cu^I$ at the limiting availabilities employed (fitted $K_{Cu(I)}$ in Supplementary Table 2); dashed lines show simulated responses for $K_{Cu(I)}$ tenfold tighter or weaker than the fitted value.

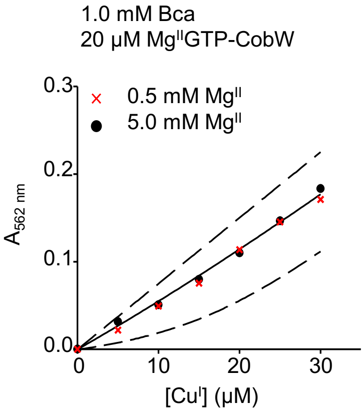


**Supplementary Figure 12. Effect of $Mg^{II}$ on $Cu^{I}$-competition experiments using Bca.** $Cu^{I}$-competition experiment as in Fig. 4c but with varying [$Mg^{II}$] (0.5 and 5.0 mM). Solid trace shows representative curve fit of the experimental data (for 5.0 mM $Mg^{II}$ dataset) to a model where CobW binds one molar equivalent $Cu^{I}$ per protein monomer. Dashed lines show simulated responses for $K_{Cu(I)}$ tenfold tighter or weaker than the fitted value. At both $Mg^{II}$ concentrations the fitted $K_{Cu(I)}$ (2.5 × $10^{-16}$ M and 2.2 × $10^{-16}$ M) were the same (within experimental error) as the value determined at cellular [$Mg^{II}$] (Supplementary Table 3). Thus, the presence of cellular [$Mg^{II}$] had negligible effect on the $Cu^{I}$-binding equilibria in these experiments.

**a**

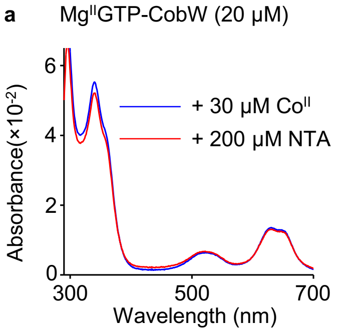


**b**

$$Co^{II}L + P \underset{}{\overset{K_{ex}}{\rightleftharpoons}} Co^{II}P + L$$

$$\frac{[Co^{II}P]}{[P]} = \frac{K_{CoII}(L)}{K_{CoII}(P)} \times \frac{[Co^{II}L]}{[L]}$$

$$= 733 \times \frac{[Co^{II}L]}{[L]}$$

where L = NTA, P = Mg$^{II}$GTP-CobW

**Supplementary Figure 13. Mg$^{II}$GTP-CobW is metal-saturated in Fig. 5 inter-metal competition experiments. a** Change in absorbance of Mg$^{II}$GTP-CobW (20 µM) upon addition of 30 µM Co$^{II}$ (blue trace), followed by 200 µM NTA (red trace) at equilibrium. **b** Reaction describing Co$^{II}$ exchange between NTA and Mg$^{II}$GTP-CobW, and corresponding relationships valid at equilibrium. In **a**, addition of Co$^{II}$ leads to stoichiometric formation of Co$^{II}$Mg$^{II}$GTP-CobW (20 µM); addition of NTA reduces $A_{339\ nm}$ to 95% of original intensity at equilibrium ratio of [Co$^{II}$NTA]/[NTA] = 0.06 consistent with the predicted 2% dissociation of Co$^{II}$ from the protein complex (calculated using the relationships in **b**) plus a dilution factor (2%) from NTA addition. The measured equilibria in Fig. 5 were all conducted at ratios [Co$^{II}$NTA]/[NTA] > 0.06 thus the high affinity site of Mg$^{II}$GTP-CobW is > 95% metalated (with either Co$^{II}$ or Zn$^{II}$) at all tested conditions.

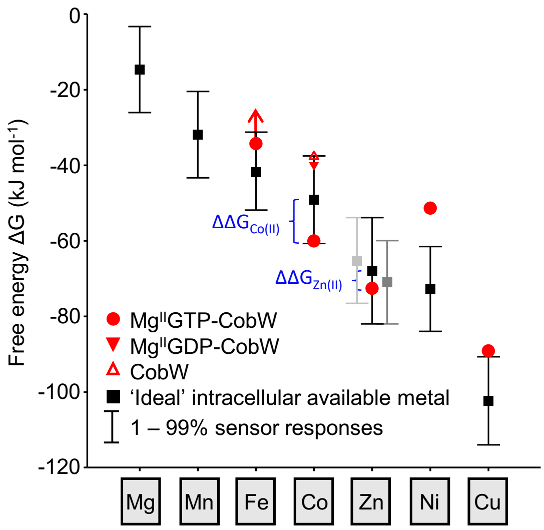


**Supplementary Figure 14. The accessible range of intracellular available free energies for metal-binding as sensors shift from 1 – 99% of their responses.** As in Fig. 6 except bars show the change in intracellular available $\Delta G$ as cognate sensors shifts from 1-99% of their responses.

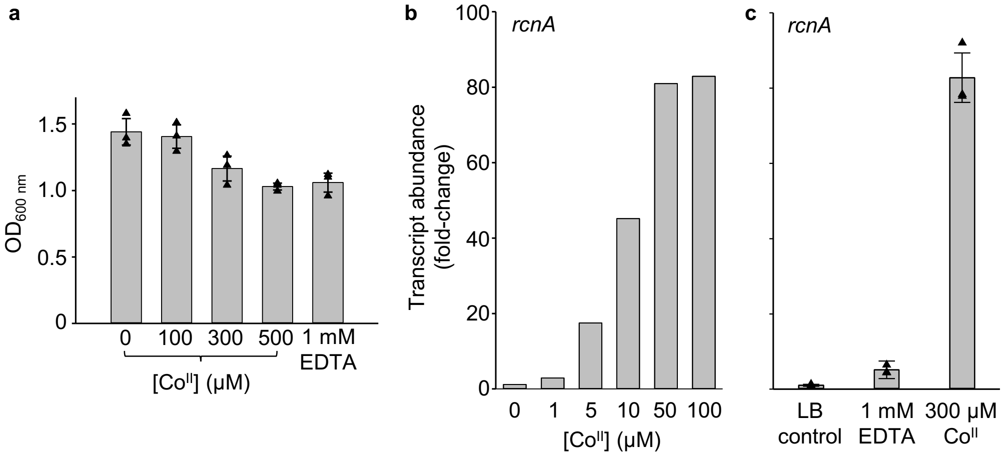


**Supplementary Figure 15. Calibration of maximum and minimum *rcnA* responses. a** $OD_{600\ nm}$ of *E. coli** cultures following 4h exposure to $Co^{II}$ or EDTA (added when $OD_{600\ nm}$ ~ 0.2). 300 µM $Co^{II}$ and 1 mM EDTA moderately inhibited growth (by 19% and 26%, respectively) relative to untreated control, and were selected as boundary conditions for *rcnA* calibration. **b-c** Transcript abundance (relative to untreated control condition, see Methods) of *rcnA* following 1h exposure of $B_{12}$-producing *E. coli** to **b** non-inhibitory $Co^{II}$ concentations (0 – 100 µM), and **c** boundary conditions determined from **a** (1 mM EDTA or 300 µM $Co^{II}$). Addition of 1 mM EDTA did not lead to further repression of *rcnA* expression but instead produced a slight increase in *rcnA* transcript abundance relative to untreated LB; thus the calibrated minimum and maximum were determined from cells grown in untreated LB and 300 µM $Co^{II}$, respectively. Data in **a** and **c** are the mean ± s.d. of n=3 biologically independent replicates; triangles represent individual experiments. Note that untreated control sample shown in **c** is replicated data from Fig 7b (these samples were cultured, and RNA collected, simultaneously, as part of a single experiment).

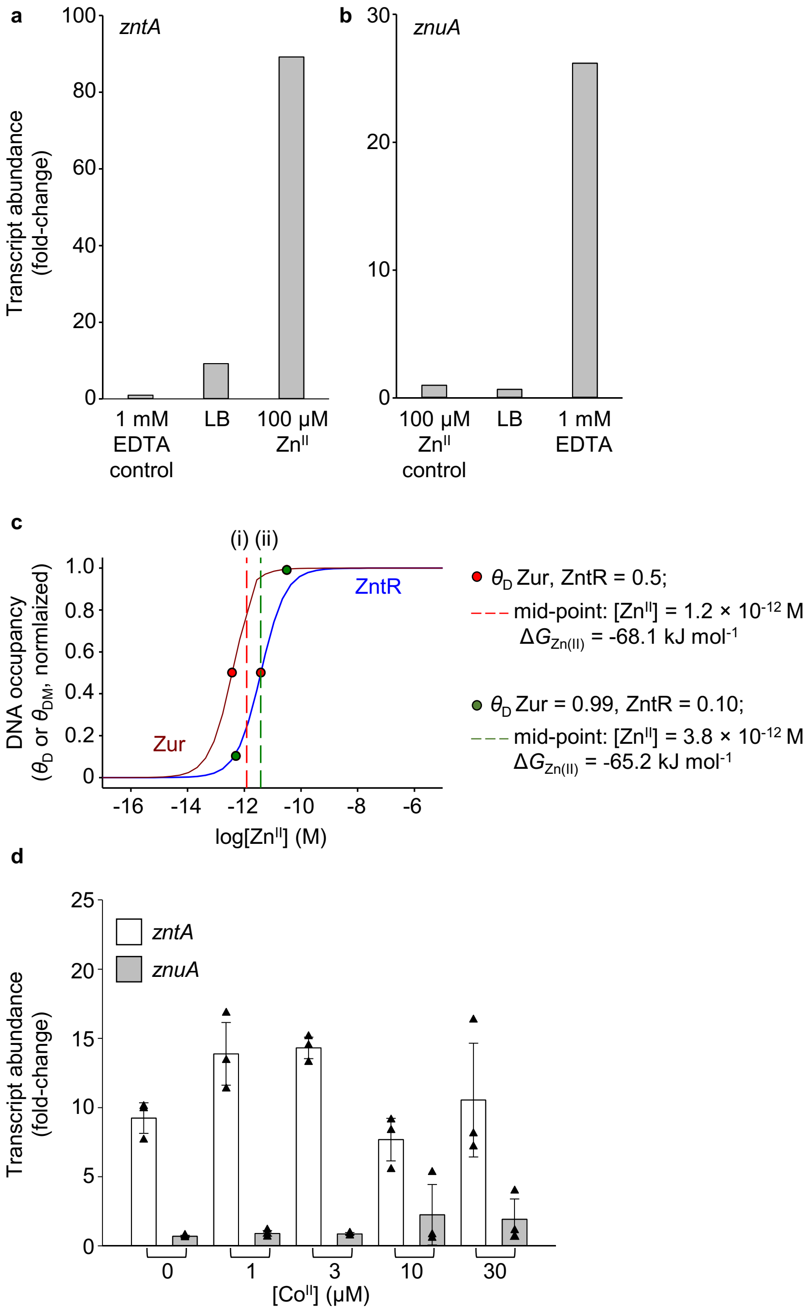


**Supplementary Figure 16. Estimation of Zn$^{II}$ availability within *E. coli** cells in defined growth conditions (conditional cells). a** Transcript abundance of *zntA* (regulated by activator ZntR) following 1h exposure of *E. coli** to EDTA (1 mM), Zn$^{II}$ (100 μM) or LB only; fold-changes were calculated relative to the EDTA-treated sample (control condition), in which minimum *zntA* expression was observed. The calculated ZntR response in LB media (equation (11)) was $\theta_D$ = 0.10. **b** Transcript abundance of *znuA* (regulated by co-repressor Zur) following 1h exposure of *E. coli** to EDTA (1 mM), Zn$^{II}$ (100 μM) or LB only; fold-changes were calculated relative to the Zn$^{II}$-treated sample (control condition), in which minimum *znuA* expression is expected. *znuA* abundance was comparable in LB and Zn$^{II}$-treated samples, thus we inferred $\theta_D \geq 0.99$ in LB media. Zur may not be fully de-repressed by addition of 1 mM EDTA, as achieving a maximal Zur-response is known to be difficult in cultured cells[12]. **c** The intracellular available [Zn$^{II}$] corresponding to half of each sensors response (red circles) differ and we previously reasoned that the intracellular available [Zn$^{II}$] in an idealised cell (*ie* neither Zn$^{II}$-deficiency nor -excess) will be mid-way between these values ('i', dotted red trace)[13]. The intracellular [$Zn^{II}$] corresponding to estimated sensor responses in LB media (from **a** and **b**; green circles) and the [$Zn^{II}$] mid-way between these values ('ii', dotted green trace) are also shown. Since (i) and (ii) differed only marginally, for the calculations in Fig. 8a we assumed the intracellular available [$Zn^{II}$] to be that of an idealised cell ($\Delta G_{Zn(II)}$ = -68.1 kJ $mol^{-1}$). **d** Transcript abundance of *zntA* and *znuA* within *E. coli** following 1h treatment with $Co^{II}$, measured by qPCR. The data show that intracellular $Zn^{II}$ availability remains constant (within experimental error) in all samples, and is consistent with that estimated in unsupplemented LB (from **a** and **b**). Data are the mean ± s.d. of n=3 biologically independent replicates (fold-changes are calculated relative to control conditions specified in **a** and **b**, where n=1 for control conditions)**.** Triangles represent individual experiments.

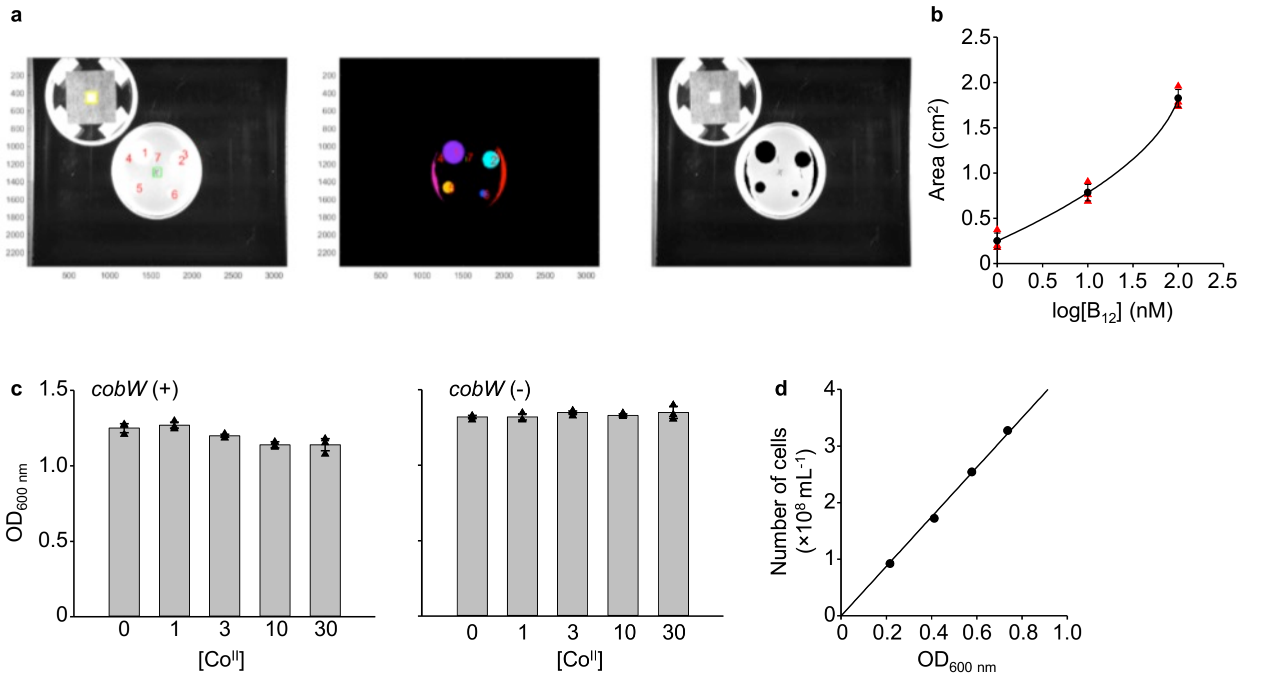


**Supplementary Figure 17. Quantification of vitamin $B_{12}$ in *E. coli*.* a** Source figure for Fig. 8b (inset): output of automated analysis of AR2680 growth areas from representative (n=3) bioassay plate of $B_{12}$ calibration standards (1-1000 nM) imaged together with a 1.0 $cm^2$ reference area. **b** Calibration curve correlating $B_{12}$ concentration and growth area from data in **a** (1 – 100 nM only, since experimental samples contained ≤ 100 nM $B_{12}$). Data (black) show average ± s.d. from n=3 experiments, red triangles are individual experiments. **c** Final $OD_{600\ nm}$ of *E. coli** cultures when harvested for $B_{12}$ analysis. **d** Representative calibration curve relating number of cells with $OD_{600\ nm}$ for *E. coli**. A correlation factor of $4.4 \pm 0.1 \times 10^8$ cells $mL^{-1}$ $OD_{600\ nm}{}^{-1}$ (mean ± s.d. of n=3 biological replicates) was determined.
