## Supplementary figures and images for "Calculating metalation in cells reveals CobW acquires Co^II^ for vitamin B_12_ biosynthesis upon binding nucleotide"

### Supplementary Data 2.tiff

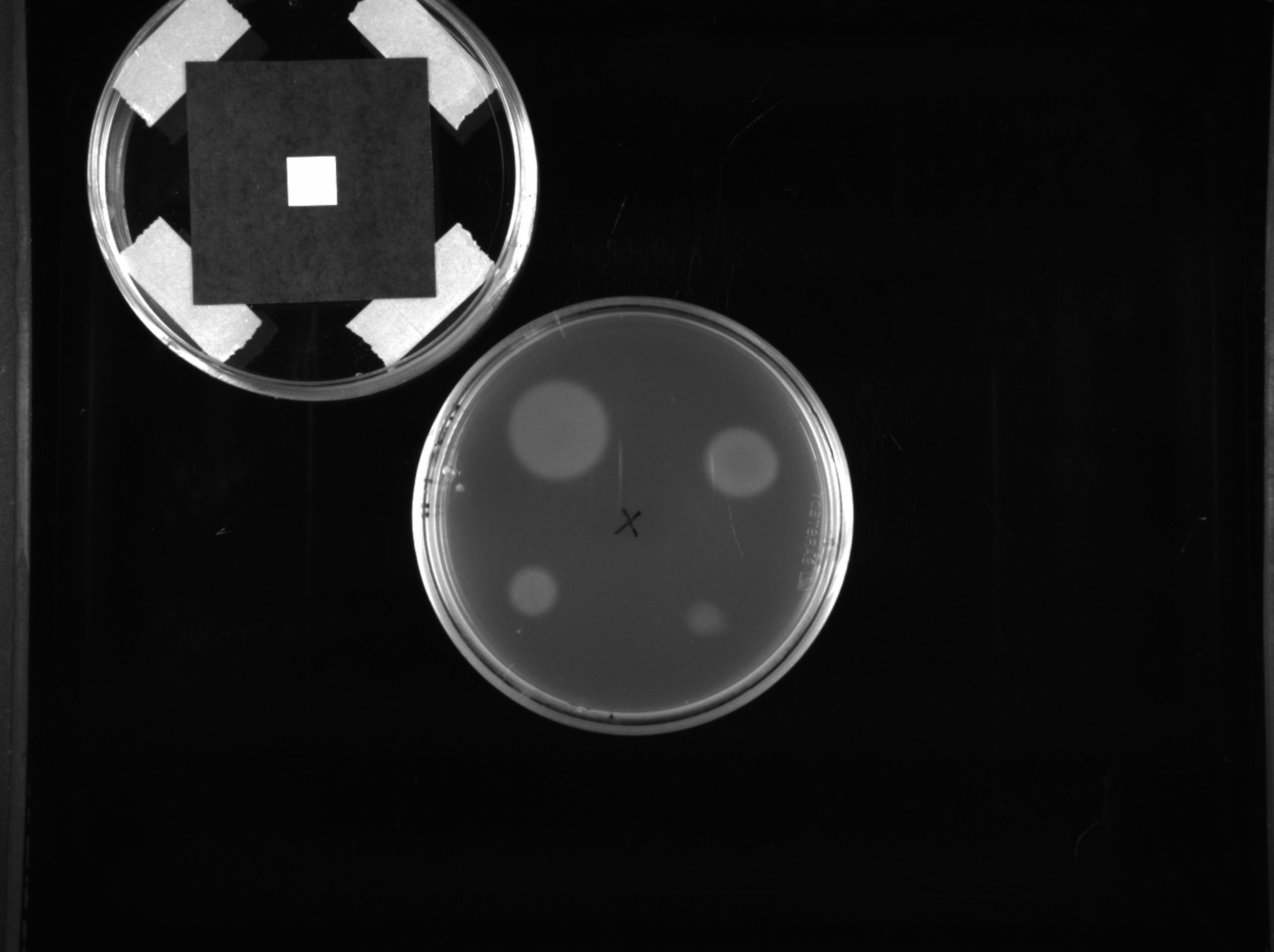
