## Supplementary Note 1 for "Calculating metalation in cells reveals CobW acquires Co^II^ for vitamin B_12_ biosynthesis upon binding nucleotide"

**Derivation of equations for calculating the *in vivo* metal occupancy of a protein (equations 1-4 in main text).**

**Derivation of equations 1 and 2**

The following equilibrium describes the dissociation of a complex of metal (M) and protein (P)

$$MP\rightleftharpoons M+P$$

The equilibrium constant for the dissociation reaction is

$$K_{\mathrm{MP}}=\frac{\left[ M \right][P]}{[MP]}$$

(1)

The standard free energy corresponding to the formation of the metal-protein complex is given by

$${\Delta G}_{\mathrm{MP}}=-\mathrm{RT}\ln\left( \frac{1}{K_{\mathrm{MP}}} \right)=RT\ln\left( K_{\mathrm{MP}} \right)$$

(2)

where R (gas constant) = 8.314 × 10^−3^ kJ K^-1^ mol^-1^ and T (temperature) = 298.15 K.

Equation (2) can be rearranged to

$$K_{\mathrm{MP}}=e^{\frac{{\Delta G}_{\mathrm{MP}}}{\mathrm{RT}}}$$

(3)

Similar equations can be written for the dissociation of a complex between a metal and any hypothetical ligand. Intracellular available ${\Delta G}_{M}$, defined as the free energy required for a hypothetical ligand to become 50% metalated from available intracellular metal (that is, [MP] = [P]), can be derived from (1) and (2), considering that [M] is the concentration of intracellular available metal

$${\Delta G}_{M}=\mathrm{RT}\ln\left( \left[ M \right] \right)$$

(4)

Equation (4) can thus be rearranged to express [M]

$$[M]=e^{\frac{{\Delta G}_{M}}{\mathrm{RT}}}$$

(5)

The difference between the free energy for protein metalation, ${\Delta G}_{\mathrm{MP}}$, and intracellular available ${\Delta G}_{M}$ is defined as

$$\Delta{\Delta G}_{M}= {\Delta G}_{\mathrm{MP}}-{\Delta G}_{M}$$

(6)

The fractional occupancy of a protein with metal is defined as

$$Fractional occupancy \left( \% \right)=100\times\frac{[MP]}{{[P]}_{\mathrm{tot}}}$$

(7)

where

$$\left[ P \right]_{\mathrm{tot}}=\left[ P \right]+[MP]$$

(8)

is the mass balance equation for the protein.

After combining (7) and (8), equation (1) can be used to give

$$Fractional occupancy \left( \% \right)=100\times\frac{\frac{\left[ M \right][P]}{K_{\mathrm{MP}}}}{\left( P+\frac{\left[ M \right]\left[ P \right]}{K_{\mathrm{MP}}} \right)}$$

$$\Rightarrow Fractional occupancy \left( \% \right)=100\times\frac{\frac{\left[ M \right]}{K_{\mathrm{MP}}}}{\left( 1+\frac{\left[ M \right]}{K_{\mathrm{MP}}} \right)}$$

Substituting (3) and (5), the equation can be written as a function of $\Delta G_{M}$ and $\Delta G_{\mathrm{MP}}$

$$Fractional occupancy (\%) =100 \times\frac{e^{\frac{{\Delta G}_{M}{-\Delta G}_{\mathrm{MP}}}{\mathrm{RT}}}}{\left( 1+e^{\frac{{\Delta G}_{M}{-\Delta G}_{\mathrm{MP}}}{\mathrm{RT}}} \right)}$$

Finally, using equation (6) gives

$$\Rightarrow Fractional occupancy (\%) =100 \times\frac{e^{\frac{{-\Delta\Delta G}_{M}}{\mathrm{RT}}}}{\left( 1+e^{\frac{{-\Delta\Delta G}_{M}}{\mathrm{RT}}} \right)}$$

**Derivation of equation 3**

Based on exchange constant in Fig. 5a

$$\frac{\left[ \mathrm{Co}^{\mathrm{II}}L \right]}{{[Zn}^{\mathrm{II}}L]}\times\frac{\left[ \mathrm{Zn}^{\mathrm{II}}P \right]}{{[Co}^{\mathrm{II}}P]}=\frac{{K_{\mathrm{Zn}^{\mathrm{II}}L}K}_{\mathrm{Co}^{\mathrm{II}}P}}{K_{\mathrm{Co}^{\mathrm{II}}L}K_{\mathrm{Zn}^{\mathrm{II}}P}}$$

Equation (1) can be used to substitute $K_{\mathrm{Zn}^{\mathrm{II}}L}$ and $K_{\mathrm{Co}^{\mathrm{II}}L}$, giving

$$\frac{\left[ \mathrm{Zn}^{\mathrm{II}}P \right]}{{[Co}^{\mathrm{II}}P]}=\frac{\left[ \mathrm{Zn}^{\mathrm{II}}L \right]}{{[Co}^{\mathrm{II}}L]}\times\frac{\left[ L \right]\left[ \mathrm{Zn}^{\mathrm{II}} \right]}{\left[ \mathrm{Zn}^{\mathrm{II}}L \right]}\times\frac{\left[ \mathrm{Co}^{\mathrm{II}}L \right]}{\left[ \mathrm{Co}^{\mathrm{II}} \right]\left[ L \right]}\times\frac{K_{\mathrm{Co}^{\mathrm{II}}P}}{K_{\mathrm{Zn}^{\mathrm{II}}P}}$$

$$\Rightarrow\frac{\left[ \mathrm{Zn}^{\mathrm{II}}P \right]}{{[Co}^{\mathrm{II}}P]}=\frac{\left[ \mathrm{Zn}^{\mathrm{II}} \right]}{\left[ \mathrm{Co}^{\mathrm{II}} \right]}\times\frac{K_{\mathrm{Co}^{\mathrm{II}}P}}{K_{\mathrm{Zn}^{\mathrm{II}}P}}$$

The equation can thus be expressed as a function of $\Delta G_{\mathrm{Zn}^{\mathrm{II}}P}$ and $\Delta G_{\mathrm{Co}^{\mathrm{II}}P}$by using (3)

$$\frac{\left[ \mathrm{Zn}^{\mathrm{II}}P \right]}{{[Co}^{\mathrm{II}}P]}=\frac{\left[ \mathrm{Zn}^{\mathrm{II}} \right]}{\left[ \mathrm{Co}^{\mathrm{II}} \right]}\times\frac{e^{\frac{{\Delta G}_{\mathrm{Co}^{\mathrm{II}}P}}{\mathrm{RT}}}}{e^{\frac{{\Delta G}_{\mathrm{Zn}^{\mathrm{II}}P}}{\mathrm{RT}}}}$$

Substituting (5) gives

$$\frac{\left[ \mathrm{Zn}^{\mathrm{II}}P \right]}{{[Co}^{\mathrm{II}}P]}=\frac{e^{\frac{{\Delta G}_{\mathrm{Co}^{\mathrm{II}}P}-{\Delta G}_{\mathrm{Co}^{\mathrm{II}}}}{\mathrm{RT}}}}{e^{\frac{{\Delta G}_{\mathrm{Zn}^{\mathrm{II}}P}-{\Delta G}_{\mathrm{Zn}^{\mathrm{II}}}}{\mathrm{RT}}}}$$

which can be rearranged using equation (6)

$$\frac{\left[ \mathrm{Zn}^{\mathrm{II}}P \right]}{{[Co}^{\mathrm{II}}P]}=\frac{e^{\frac{{\Delta\Delta G}_{\mathrm{Co}^{\mathrm{II}}P}}{\mathrm{RT}}}}{e^{\frac{{\Delta\Delta G}_{\mathrm{Zn}^{\mathrm{II}}P}}{\mathrm{RT}}}}$$

(9)

The mass balance equation for the protein species present in the system in Fig. 5a can be written as

$$\left[ P \right]_{\mathrm{tot}}=\left[ \mathrm{Co}^{\mathrm{II}}P \right]+\left[ \mathrm{Zn}^{\mathrm{II}}P \right]+[P]$$

(10)

This can be rearranged by dividing both terms by $[\mathrm{Co}^{\mathrm{II}}P]$

$$\Rightarrow\frac{\left[ P \right]_{\mathrm{tot}}}{[\mathrm{Co}^{\mathrm{II}}P]}=\frac{\left[ \mathrm{Co}^{\mathrm{II}}P \right]+\left[ \mathrm{Zn}^{\mathrm{II}}P \right]+[P]}{[\mathrm{Co}^{\mathrm{II}}P]}$$

$$\Rightarrow\frac{\left[ P \right]_{\mathrm{tot}}}{[\mathrm{Co}^{\mathrm{II}}P]}=1+ \frac{[\mathrm{Zn}^{\mathrm{II}}P]}{[\mathrm{Co}^{\mathrm{II}}P]}+\frac{\left[ P \right]}{[\mathrm{Co}^{\mathrm{II}}P]}$$

$$\Rightarrow\frac{[\mathrm{Co}^{\mathrm{II}}P]}{\left[ P \right]_{\mathrm{tot}}}=\frac{1}{1+ \frac{[\mathrm{Zn}^{\mathrm{II}}P]}{[\mathrm{Co}^{\mathrm{II}}P]}+\frac{\left[ P \right]}{[\mathrm{Co}^{\mathrm{II}}P]}}$$

Substituting (1) gives

$$\frac{[\mathrm{Co}^{\mathrm{II}}P]}{\left[ P \right]_{\mathrm{tot}}}=\frac{1}{1+ \frac{[\mathrm{Zn}^{\mathrm{II}}P]}{[\mathrm{Co}^{\mathrm{II}}P]}+\frac{K_{\mathrm{Co}^{\mathrm{II}}P}}{[\mathrm{Co}^{\mathrm{II}}]}}$$

Substituting (3) and (5) gives

$$\frac{[\mathrm{Co}^{\mathrm{II}}P]}{\left[ P \right]_{\mathrm{tot}}}=\frac{1}{1+ \frac{[\mathrm{Zn}^{\mathrm{II}}P]}{[\mathrm{Co}^{\mathrm{II}}P]} + \frac{e^{\frac{{\Delta G}_{\mathrm{Co}^{\mathrm{II}}P}}{\mathrm{RT}}}}{e^{\frac{{\Delta G}_{\mathrm{Co}^{\mathrm{II}}}}{\mathrm{RT}}}}}$$

which can then be rearranged using (6)

$$\frac{[\mathrm{Co}^{\mathrm{II}}P]}{\left[ P \right]_{\mathrm{tot}}}=\frac{1}{1+ \frac{[\mathrm{Zn}^{\mathrm{II}}P]}{[\mathrm{Co}^{\mathrm{II}}P]} +e^{\frac{{\Delta\Delta G}_{\mathrm{Co}^{\mathrm{II}}}}{\mathrm{RT}}}}$$

Finally, substituting (9) and rearranging the equation gives

$$\frac{[\mathrm{Co}^{\mathrm{II}}P]}{\left[ P \right]_{\mathrm{tot}}}=\frac{1}{1+e^{\frac{{\Delta\Delta G}_{\mathrm{Co}^{\mathrm{II}}}-{\Delta\Delta G}_{\mathrm{Zn}^{\mathrm{II}}}}{\mathrm{RT}}} +e^{\frac{{\Delta\Delta G}_{\mathrm{Co}^{\mathrm{II}}}}{\mathrm{RT}}}}$$

$$\Rightarrow\frac{[\mathrm{Co}^{\mathrm{II}}P]}{\left[ P \right]_{\mathrm{tot}}}=\frac{1}{e^{\frac{{\Delta\Delta G}_{\mathrm{Co}^{\mathrm{II}}}}{\mathrm{RT}}} \left( e^{\frac{{-\Delta\Delta G}_{\mathrm{Co}^{\mathrm{II}}}}{\mathrm{RT}}}+e^{\frac{-{\Delta\Delta G}_{\mathrm{Zn}^{\mathrm{II}}}}{\mathrm{RT}}} +1 \right)}$$

$$\Rightarrow\frac{[\mathrm{Co}^{\mathrm{II}}P]}{\left[ P \right]_{\mathrm{tot}}}=\frac{e^{\frac{{-\Delta\Delta G}_{\mathrm{Co}^{\mathrm{II}}}}{\mathrm{RT}}}}{\left( e^{\frac{{-\Delta\Delta G}_{\mathrm{Co}^{\mathrm{II}}}}{\mathrm{RT}}}+e^{\frac{-{\Delta\Delta G}_{\mathrm{Zn}^{\mathrm{II}}}}{\mathrm{RT}}} +1 \right)}$$

The fractional occupancy of the protein with Co^II^ can thus be calculated using (7)

$$\Rightarrow Fractional occupancy \left( \% \right)=100\times\frac{[\mathrm{Co}^{\mathrm{II}}P]}{\left[ P \right]_{\mathrm{tot}}}=100\times\frac{e^{\frac{{-\Delta\Delta G}_{\mathrm{Co}^{\mathrm{II}}}}{\mathrm{RT}}}}{\left( e^{\frac{{-\Delta\Delta G}_{\mathrm{Co}^{\mathrm{II}}}}{\mathrm{RT}}}+e^{\frac{-{\Delta\Delta G}_{\mathrm{Zn}^{\mathrm{II}}}}{\mathrm{RT}}} +1 \right)}$$

**Derivation of equation 4**

Let us consider a protein which can bind n different metals (namely M_1_ to M_n_). The mass balance equation for the protein can be written as

$${[P]}_{\mathrm{tot}}=\left[ P \right]+\left[ M_{1}P \right]+\left[ M_{2}P \right]+\ldots+\left[ M_{n}P \right]$$

(11)

The fractional occupancy of protein with any of the metals (e.g. with metal M_i_) can be calculated using the expression

$$Fractional occupancy \left( \% \right)=100\times\frac{[M_{i}P]}{{[P]}_{\mathrm{tot}}}$$

Using (1), this can be rewritten as

$$Fractional occupancy \left( \% \right)=100\times\frac{\left[ M_{i} \right]\left[ P \right]}{K_{M_{i}}{[P]}_{\mathrm{tot}}}$$

Substituting (11) gives

$$Fractional occupancy (\%) =100 \times\frac{\left[ M_{i} \right]\left[ P \right]}{K_{M_{i}}\left( \left[ P \right]+\left[ M_{1}P \right]+\left[ M_{2}P \right]+\ldots+[M_{n}P] \right)}$$

which can be then rearranged to

$$Fractional occupancy (\%) =100 \times\frac{\left[ M_{i} \right]}{K_{M_{i}}\left( 1+\frac{\left[ M_{1}P \right]}{[P]}+\frac{\left[ M_{2}P \right]}{[P]}+\ldots+\frac{\left[ M_{n}P \right]}{[P]} \right)}$$

Equation (1) can be used once again at the denominator, giving

$$Fractional occupancy (\%) =100 \times\frac{\left[ M_{i} \right]}{K_{M_{i}}\left( 1+\frac{[M_{1}]}{K_{M_{1}}}+\frac{[M_{2}]}{K_{M_{2}}}+\ldots+\frac{[M_{n}]}{K_{M_{n}}} \right)}$$

Using equations (3) and (5), the fractional occupancy can be written as a function of intracellular available ${\Delta G}_{M_{i}}s$ and of the free energies of protein-metal complex formation ${\Delta G}_{M_{i}P}s$

$$Fractional occupancy (\%) =100 \times\frac{e^{\frac{{\Delta G}_{M_{i}}}{\mathrm{RT}}}}{e^{\frac{{\Delta G}_{M_{i}P}}{\mathrm{RT}}}\left( 1+\frac{e^{\frac{{\Delta G}_{M_{1}}}{\mathrm{RT}}}}{e^{\frac{{\Delta G}_{M_{1}P}}{\mathrm{RT}}}}+\frac{e^{\frac{{\Delta G}_{M_{2}}}{\mathrm{RT}}}}{e^{\frac{{\Delta G}_{M_{2}P}}{\mathrm{RT}}}}+\ldots+\frac{e^{\frac{{\Delta G}_{M_{n}}}{\mathrm{RT}}}}{e^{\frac{{\Delta G}_{M_{n}P}}{\mathrm{RT}}}} \right)}$$

This equation can be rearranged, and substituting (6) gives

$$Fractional occupancy (\%) =100 \times\frac{e^{\frac{{\Delta G}_{M_{i}}{-\Delta G}_{M_{i}P}}{\mathrm{RT}}}}{\left( 1+e^{\frac{{\Delta G}_{M_{1}}{-\Delta G}_{M_{1}P}}{\mathrm{RT}}}+e^{\frac{{\Delta G}_{M_{2}}{-\Delta G}_{M_{2}P}}{\mathrm{RT}}}+\ldots+e^{\frac{{\Delta G}_{M_{n}}{-\Delta G}_{M_{n}P}}{\mathrm{RT}}} \right)}$$

$$\Rightarrow Fractional occupancy (\%) =100 \times\frac{e^{\frac{{-\Delta\Delta G}_{M_{i}}}{\mathrm{RT}}}}{\left( 1+e^{\frac{{-\Delta\Delta G}_{M_{1}}}{\mathrm{RT}}}+e^{\frac{{-\Delta\Delta G}_{M_{2}}}{\mathrm{RT}}}+\ldots+e^{\frac{{-\Delta\Delta G}_{M_{n}}}{\mathrm{RT}}} \right)}$$

Finally, the equation for the fractional occupancy can be written the more compact form

$$\Rightarrow Fractional occupancy (\%) =100 \times\frac{e^{\frac{{-\Delta\Delta G}_{M_{i}}}{\mathrm{RT}}}}{\left( 1+\sum_{k=1}^{k=n} e^{\frac{{-\Delta\Delta G}_{M_{k}}}{\mathrm{RT}}} \right)}$$
