## Supplementary Note 2 for "Calculating metalation in cells reveals CobW acquires Co^II^ for vitamin B_12_ biosynthesis upon binding nucleotide"

**MATLAB code for automated area analysis of bacterial growth spots**

This file contains:

1. **Description of the MATLAB code**
2. **The MATLAB code and instructions for use**

**1. Description of the MATLAB code**

The objective of the code is to provide a form of image segmentation; identifying spots of bacterial growth on an agar plate and measuring their area.

Digital photographic images form a set of values that correspond to the photons registered at a certain pixel on the camera sensor (alternatively this can be described as the intensity or brightness). Areas of bacterial growth are brighter than the background media, enabling detection and separation of the two, following identification of a suitable threshold value of intensity: The image is split into pixels we identify as either bacteria or media. The act of choosing a threshold value is central to many image segmentation methods.

In addition to areas of bacterial growth and media, there are artefacts present in the image. An artefact is any item that is not of interest e.g. the Petri dish, dirt or scratches on the dish lid/base. The first step in image processing uses a mask to remove any artefacts outside of the Petri dish and the Petri dish edge, leaving only the plate area. Remaining issues include (i) scratches/dirt on Petri dishes, (ii) uneven illumination of the media due to reflected light from the Petri dish edge and (iii) uneven thickness of the agar across the plate. These are difficult to correct in an automated way and should be minimised as much as possible during plate preparation and image capture.

To choose the threshold value, we first need to identify the values of intensity that describe the media. The media is relatively homogenous but will likely contain slight variations in brightness across the plate. Moreover, there will be noise in the image, which arises from photon counting (Poisson noise) and sensor performance. To characterise media intensity, the MATLAB routine calculates the mean ($\mu$) and standard deviation ($\sigma$) of 1000 pixels near the centre of the Petri dish that contain only media (i.e. no culture pixels). The centre of the plate can be marked manually with a black cross before image capture without any effect on this calculation.

The threshold is then set to be a value, $N$, that is a number of standard deviations above the mean value of the media, i.e., $N=\mu+m\sigma$. The best value of $m$ is somewhat subjective, and largely depends on what works in each individual case (there will be variability e.g. culture, media, instrument). There is no objective way to set the threshold value, as there is no discrete edge to the bacterial growth spot. The edge of the growth spot and media are blurred, meaning the intensity curve from spot centre to media decreases smoothly: the $m$ value should be selected following optimisation with different $m$ values. We have found that varying $m$ over a range of 3, e.g. $6\leq m\leq8$ , has negligible influence on the returned value of bacterial growth area. The same $m$ value should be used across a batch of plates which includes at least one calibration plate.

After setting the threshold value, the image is thresholded into bacterial and media pixels. The routine will then find and label each set of contiguous image pixels as defined regions on the plate. Each will be a different bacterial growth spot, but may also include other areas of brightness e.g. from the plate edge; the latter can be ignored. Once labelled, the number of pixels in each region is counted and provides the area of each bacterial growth spot. The area in cm^2^ is determined with reference to a square centimetre included in the image.

**2. The MATLAB code and instructions for use**

**System requirements**

This code has been tested using MATLAB R2019a Version 9.6.0.1072779 running on Windows 10.

**Installation guide**

MATLAB is available to download here: [https://uk.mathworks.com/products/matlab.html](about:blank)

**Input**

This code calculates areas (in cm^2^) of bacterial growth spots on a Petri dish, prepared as described in Methods. Required input is one or more .tif image files taken on black background: Each .tif file must include both the plate with growth spots to be quantified and a 1cm^2^ reference area in the same image (for scaling), as described above and in Methods. An example image is included in Supplementary Data 2.

**Instructions for use**

To run the MATLAB routine:

1. In MATLAB open a new script.
2. Copy the following MATLAB routine (courier new font, below) to the MATLAB editor window and save to the MATLAB folder as “find_blobs_new.m”.
3. Type “find_blobs_new” in the command window and press enter.
4. When prompted, select picture (.tif) file(s) for analysis. Single or multiples .tif files can be selected.
5. When prompted, select points at the bottom left and top right of the 1cm^2^ reference area, then press enter. This will be used to locate the 1cm^2^ reference area.
6. When prompted, select a position in the centre of the Petri dish (where no bacterial growth is observed), then press enter. This will be used to determine the background signal.
7. An annotated image (.png) file will be generated in the current folder window, showing all detected spots and assigning an integer number to each. Outputs in the command window display the area and radius corresponding to each of the detected spots.
8. Repeat steps 5-7 for each image.

**Variable settings (e.g. defining** $\boldsymbol{m}$**)**

Variable settings are editable at the start of the routine and include (i) the diameter of the Petri dish, (ii) the minimum size of bacterial growth spot (blob) and (iii) the detection threshold. For the current analysis, default values for all variable settings were used. To change the detection threshold, use the command “find_blobs_new(‘StandDev’,m)”, where *m* is a number > 0. The default setting (in find_blobs_new.m) is 8. A smaller value of *m* will lower the detection threshold: This allows fainter spots to be detected but may increase false-positive signals, and vice versa (see above description of the MATLAB code).

**MATLAB Routine**

START COPYING:

```
function find_blobs_new(varargin)

p=inputParser;

%default values
expand_box=20 ; % for finding the cm box - incase user clicks to close to edge
addOptional(p,'expand_box',expand_box,@isnumeric);

PetriDishDiam=1200;  % diameter of petri dish
addOptional(p,'PetriDishDiam',PetriDishDiam,@isnumeric);

rad_low=150; % annulus radii for masks
addOptional(p,'rad_low',rad_low,@isnumeric);

mediaBoxLen=50; % half length of media box
addOptional(p,'mediaBoxLen',mediaBoxLen,@isnumeric);

min_blob=500; % minimum size of blobs
addOptional(p,'min_blob',min_blob,@isnumeric);

StandDev=8; % standard deviation for threshold
addOptional(p,'StandDev',StandDev,@isnumeric);

manualSelect='no'; % turn on/off background selection
validSelect = {'no','yes'};
checkSelect = @(x) any(validatestring(x,validSelect));

addOptional(p,'manualSelect',manualSelect,checkSelect);

debugPlot='no'; % turn on/off debugging plots
addOptional(p,'debugPlot',debugPlot,checkSelect);

parse(p,varargin{:});


%Import image file
[FileName,PathName,FilterIndex] = uigetfile('.tif','Select location of tif image file.','multiselect','on');
DateString=strrep(datestr(datetime('now')),':','_');


 %------------------------
 %Open file to write to
 OutputFile=strcat('blob_details_',DateString,'.csv');
 fileID = fopen(OutputFile,'w');
 %------------------------

if iscellstr(FileName)==1
    numberFiles=length(FileName);
else
   numberFiles=1;
end

for jj=1:numberFiles
    if iscellstr(FileName)==0 ThisFile=FileName; else ThisFile=FileName{jj}; end

    Image = imread(strcat(PathName,ThisFile));
    ImageSize=size(Image);

    % Get rid of transparency channel
    if ImageSize(3)== 4 Image(:,:,4) = []; end
    GrayImage=rgb2gray(Image);
    im_sz=size(GrayImage);

    %find 1cm mark
    [BoxPos,scale]=find_square(GrayImage,p);

    [masks,xPetrieCenter,yPetrieCenter]=create_mask(GrayImage,im_sz,p);

    GrayImageMasked=imgaussfilt(double(GrayImage),2);
    GrayImageMasked(find(masks == 0))=0;

    thresholdLevel=find_threshold(GrayImageMasked,masks,p,xPetrieCenter,yPetrieCenter);

    %Threshold image
    maxGray=max(GrayImageMasked(:));
    BinaryImage = imbinarize(GrayImageMasked/double(maxGray),thresholdLevel);

    [labeledImage, numBlobs] = bwlabel(BinaryImage);

    [blobSizes,labeledImage]=choose_blobs(numBlobs,labeledImage,p);
    [numBlobsLarge,na]=size(blobSizes);

    plot_blob_images(GrayImage,labeledImage,BinaryImage,BoxPos,DateString,blobSizes,ThisFile,xPetrieCenter,yPetrieCenter,p);

    %------------------------
    fprintf(fileID,'\n');
    fprintf(fileID,strrep(ThisFile,'.tif',''));
    fprintf(fileID,'\n');
    fprintf(fileID,'%s,%s,%s,%s\n','Blob Number','Area (Pixels)','Area (cm2)','Radius (cm)');

    for i=1:numBlobsLarge
        area=blobSizes(i,1)/scale/scale;
        radius=(area/pi)^0.5;
        fprintf('%d has area %4.3f cm2 and radius cm %4.3f.\n',i,area,radius);
        fprintf(fileID,'%d, %d, %4.3f, %4.3f\n',i,blobSizes(i,1),area,radius);
    end
    fprintf(fileID,'\n');
     %------------------------
end

% close text file
fclose(fileID);

end

function [BoxPos,scale]=find_square(GrayImage,p)
    fig = figure('position', [0, 0, 1000, 1000]);
    imshow(GrayImage);
    text(500,100,['Select points near bottom left and top right of'...
                  ' square. Press Enter once finished.'],'Color','red','FontSize',18);
    axis on
    fprintf(['Select points near bottom left and top right of'...
                ' square. Press Enter once finished.\n']);
    [xbox,ybox]=getpts;

    close(fig)

    if xbox(1) > xbox(2)
        temp=xbox(1);
        xbox(1)=xbox(2); xbox(2)=temp;
    end
    if ybox(1) > ybox(2)
        temp=ybox(1);
        ybox(1)=ybox(2); ybox(2)=temp;
    end
    xbox(1)=round(xbox(1))-p.Results.expand_box; xbox(2)=round(xbox(2))+p.Results.expand_box;
    ybox(1)=round(ybox(1))-p.Results.expand_box; ybox(2)=round(ybox(2))+p.Results.expand_box;

    BinaryImage = imbinarize(GrayImage(ybox(1):ybox(2),xbox(1):xbox(2)));
    [labeledImage, numBlobs] = bwlabel(BinaryImage);
    [YPixbox,XPixbox]=find(labeledImage==1);
    LowerLeftCornerBox=[min(XPixbox)+xbox(1),min(YPixbox)+ybox(1)]; % bottom left hand corner box (as shown in imshow)


    cm=max(XPixbox)-min(XPixbox); rm=max(YPixbox)-min(YPixbox);
    scale= sqrt(numel(XPixbox)) ; % scaling, pixels per cm

    BoxPos=[LowerLeftCornerBox(1),LowerLeftCornerBox(2),cm,rm];
end

function [masks,xPetrieCenter,yPetrieCenter]=create_mask(GrayImage,im_sz,p)

    %Select center of Petri dish
    fig = figure('position', [0, 0, 1000, 1000]);
    imshow(GrayImage);
    text(500,100,['Select center of Petri dish'...
                  '. Press Enter once finished.'],'Color','red','FontSize',18);
    axis on
    fprintf(['Select center of Petri dish'...
                '. Press Enter once finished.\n']);
    [xPetrieCenter,yPetrieCenter]=getpts;

    close(fig)

    % create polar coords for mask
    y=imresize(1:im_sz(1),[im_sz(2),im_sz(1)]);
    y=transpose(y);
    x=imresize(1:im_sz(2),[im_sz(1),im_sz(2)]);

    masks=ones(im_sz(1),im_sz(2),'uint8');
    MaskBoundary=p.Results.PetriDishDiam/2-p.Results.rad_low;

    temp=ones(im_sz(1),im_sz(2),'uint8');
    r=((y-yPetrieCenter).^2+(x-xPetrieCenter).^2).^0.5;
    temp(find(r> MaskBoundary ))=0;
    masks(:,:)=temp;

end

function thresholdLevel=find_threshold(GrayImageMasked,masks,p,xPetrieCenter,yPetrieCenter)
%Select media to calculate background level for threshold
    if strcmp(p.Results.manualSelect,'yes')
        [row,col]=find(masks ~= 0);
        low_bound=[min(row),min(col)];
        up_bound=[max(row),max(col)];

        fig=figure('position', [0, 0, 1000, 1000]);
        imshow(GrayImageMasked(low_bound(1):up_bound(1),low_bound(2):up_bound(2)),[0 80])
        text(50,50,['Draw box over media to set background'...
                  '. Double click in the box once finished.'],'Color','red','FontSize',16);
        h = imrect;
        position = wait(h);
        boxVals=h.getPosition;
        mediaBox=GrayImageMasked(low_bound(1)+boxVals(2):low_bound(1)+...
                                  boxVals(2)+boxVals(4),low_bound(2)+boxVals(1):low_bound(2)+boxVals(1)+boxVals(3));

        close(fig)
        boxMean=median(mediaBox(:));
        boxSTD=std(double(mediaBox(:)));
    else
        mediaBox=GrayImageMasked(fix(yPetrieCenter)-p.Results.mediaBoxLen:fix(yPetrieCenter)+p.Results.mediaBoxLen,...
                                fix(xPetrieCenter)-p.Results.mediaBoxLen:fix(xPetrieCenter)+p.Results.mediaBoxLen);

        if strcmp(p.Results.debugPlot,'yes')
            fig_debug=figure('position', [0, 0, 500, 500],'visible','off');
            image(mediaBox);
            text(1,1,'Region selected for background calcution','Color','red','FontSize',16);
            colormap gray
            saveas(fig_debug,'media_region_debug.png')
            close(fig_debug);
        end

        Gaussian=@(x,p) p(1)*exp(-(x-p(2)).^2/2./p(3))+p(4);
        %doubleGauss=@(x,p) p(1)*exp(-(x-p(2)).^2/2./p(3))+p(4)*exp(-(x-p(5)).^2/2./p(6))+p(7);
        objFn=@(q,x,data) sum( (data-Gaussian(x,q)).^2);

        histfig=figure('visible','off');
        h=histogram(mediaBox(:));
        xvals=h.BinEdges(1:h.NumBins)+h.BinWidth/2;
        yvals=h.Values;
        close(histfig);
        [peakHeight,loc]=findpeaks(yvals);
        [peakHeightd,locdesc]=sort(peakHeight,'descend');

        peaklocs=loc( locdesc(1) );
        xvalred=xvals(max([peaklocs-10,1]):min([peaklocs+10,length(xvals)]));
        yvalred=yvals(max([peaklocs-10,1]):min([peaklocs+10,length(xvals)]));
        guessParam=[peakHeightd(1),xvalred(11),2,min(yvalred)];
        options = optimoptions('fsolve','Display','none','Algorithm','levenberg-marquardt');
        f= @(p) objFn(p,xvalred,yvalred);
        [param,fval,exitflag,output]=fsolve(f, guessParam ,options);

        if strcmp(p.Results.debugPlot,'yes')
            fig_debug=figure('visible','off');
            plot(xvalred,yvalred,'r');
            hold on
            plot(xvalred,Gaussian(xvalred,param),'b' )
            saveas(fig_debug,'media_region_fit_debug.png')
            hold off
            close(fig_debug);

        end

        boxMean=param(2);
        boxSTD=sqrt(param(3));

    end
  maxGray=max(GrayImageMasked(:));
  thresholdLevel=(double(boxMean)+p.Results.StandDev*boxSTD)/double(maxGray);
end

function [blob_sizes,labeledImage]=choose_blobs(numBlobs,labeledImage,p)
    %find large blobs and order
    blob_sizes=[];
    for i=1:numBlobs
        [r,c]=find(labeledImage==i);
        if numel(r) > p.Results.min_blob
           blob_sizes=[blob_sizes; [numel(r),min(r),min(c),i]];
        else
            labeledImage(r,c)=0;
        end
    end
    [vals,index]=sort(blob_sizes(:,1),'descend');
    blob_sizes=blob_sizes(index,:);
end

function plot_blob_images(GrayImage,labeledImage,BinaryImage,BoxPos,DateString,blob_sizes,ThisFile,xPetrieCenter,yPetrieCenter,p)
    [numBlobsLarge,na]=size(blob_sizes);

    sav_fig=figure('visible','off');
    fig_pos = get(sav_fig,'Position');
    fig_pos(3)=fig_pos(3).*3.;
    set(sav_fig,'Position',fig_pos);

    sav_fig_axa = axes('Position',[0.075+0*(0.05+0.25) 0.05 0.25 0.8]);
    sav_fig_axb = axes('Position',[0.075+1*(0.05+0.25) 0.05 0.25 0.8]);
    sav_fig_axc = axes('Position',[0.075+2*(0.05+0.25) 0.05 0.25 0.8]);

    %original image
    image(sav_fig_axa,GrayImage);
    title(sav_fig_axa,ThisFile,'FontSize',18,'FontWeight','Bold');
    rectangle(sav_fig_axa,'Position',BoxPos,'EdgeColor','y','LineWidth',2);
    mediaBoxPos=[xPetrieCenter-p.Results.mediaBoxLen,yPetrieCenter-p.Results.mediaBoxLen,...
                2.*p.Results.mediaBoxLen,2.*p.Results.mediaBoxLen];
    rectangle(sav_fig_axa,'Position',mediaBoxPos,'EdgeColor','g','LineWidth',1);
    axis off

    coloredLabels = label2rgb (labeledImage, 'hsv', 'k', 'shuffle');
    image(sav_fig_axb,coloredLabels);
    for i=1:numBlobsLarge
        radius=(blob_sizes(i,1)/pi)^0.5;
        XCenter=blob_sizes(i,3)+radius;
        YCenter=blob_sizes(i,2)+radius;
        text(sav_fig_axa,XCenter,YCenter,num2str(i),'Color','red','FontSize',14);
        text(sav_fig_axb,XCenter,YCenter,num2str(i),'Color','red','FontSize',14) ;
    end
    axis off

    image(sav_fig_axc,double(GrayImage).*imcomplement(BinaryImage))
    axis off
    colormap gray

    saveas(sav_fig,strcat('blob_pictures_',strrep(ThisFile,'.tif','_'),DateString,'.png'))

end
```

STOP COPYING.
