## Supplementary Note 3 for "Calculating metalation in cells reveals CobW acquires Co^II^ for vitamin B_12_ biosynthesis upon binding nucleotide"

**Dynafit Scripts**

**Dynafit script to describe competition between EGTA and Mg^II^GMPPNP-CobW for Co^II^**

```
[model]
EGTA binds 1 Co(II) per monomer. Mg(II)GMPPNP-CobW binds 1 Co(II) per monomer.

[components]
; C = EGTA (competitor)
; P = CobW (protein)
; M = Co(II) (metal)

[task]
task = fit
data = equilibria

[mechanism]
C + M <==> MC : Keq1 dissociation
P + M <==> MP : Keq2 dissociation

[concentrations]; micromolar
C = 40
P = 20

[constants]; micromolar
Keq1 = 0.00789
Keq2 = 0.001 ?

[responses]
MP = 0.0028?

[data]
variable M
offset = auto
set MgGMPPNPCobW

[set:MgGMPPNPCobW]
0	0
6	0.008139
12	0.016892
18	0.022708
24	0.029845
30	0.036144
36	0.043848
48	0.052472
60	0.054842
72	0.055745
84	0.056828
96	0.05663
[end]
```

**Dynafit script to describe competition between EGTA and Mg^II^GTPγS-CobW for Co^II^**

```
[model]
EGTA binds 1 Co(II) per monomer. Mg(II)GTPyS-CobW binds 1 Co(II) per monomer.

[components]
; C = EGTA (competitor)
; P = CobW (protein)
; M = Co(II) (metal)

[task]
task = fit
data = equilibria

[mechanism]
C + M <==> MC : Keq1 dissociation
P + M <==> MP : Keq2 dissociation


[concentrations]; micromolar

C = 1000
P = 20

[constants]; micromolar

Keq1 = 0.00789
Keq2 = 0.000001 ?


[responses]

MP = 0.0028

[data]

variable M
offset = auto
set MgGTPySCobW

[set:MgGTPySCobW]

0	0
5	0.006432
10	0.010445
15	0.014342
20	0.017468
25	0.019309
30	0.021115
40	0.023736

[end]
```

**Dynafit script to describe competition between Fura-2 and CobW for Co^II^**

```
[model]
Fura2 binds 1 Co(II) per monomer. CobW binds 1 Co(II) per monomer.

[components]
; C = Fura2 (competitor)
; P = CobW (protein)
; M = Co(II) (metal)

[task]
task = fit
data = equilibria

[mechanism]
C + M <==> MC : Keq1 dissociation
P + M <==> MP : Keq2 dissociation


[concentrations]; micromolar
P = 37
C = 10

[constants]; micromolar

Keq1 = 0.00864
Keq2 = 0.5 ?


[responses]
MC = -1 ?

[data]
variable M
offset = auto
set CobW

[set:CobW]

0.000		1
2.000		0.786898009
4.000		0.616778285
6.000		0.496297227
8.000		0.399378345
10.000	0.338180665
14.000	0.242332963
18.000	0.192537668
26.000	0.125832535
34.000	0.093327689
50.000	0.048680581
66.000	0.032373569
82.000	0.041403781

[end]
```

**Dynafit script to describe competition between Fura-2 and Mg^II^GDP-CobW for Co^II^**

```
[model]
Fura-2 binds 1 Co(II) per monomer. Mg(II)GDP-CobW binds 1 Co(II) per monomer.

[components]
; C = Fura-2 (competitor)
; P = CobW (protein)
; M = Co(II) (metal)

[task]
task = fit
data = equilibria

[mechanism]
C + M <==> MC : Keq1 dissociation
P + M <==> MP : Keq2 dissociation


[concentrations]; micromolar
C = 8.06
P = 20

[constants]; micromolar
Keq1 = 0.00864
Keq2 = 0.01 ?

[responses]
MC = -1 ?

[data]
variable M
offset = auto
set MgGDPCobW

[set:MgGDPCobW]
0	1
2	0.870760656
4	0.670936169
6	0.495969137
8	0.359041645
10	0.26705313
12	0.197223607
15	0.154501562
18	0.116076522
22	0.07842911
26	0.060364067
30	0.047021687
34	0.040090394
38	0.035768074
42	0.031218785
46	0.028668037

[end]
```

**Dynafit script to describe competition between EGTA and Mg^II^GTP-CobW for Co^II^**

```
[model]

EGTA binds 1 Co per monomer. Mg(II)GTP-CobW binds 1 Co(II) per monomer.

[components]

; C = EGTA (competitor)
; P = CobW (protein)
; M = Co(II) (metal)

[task]

task = fit
data = equilibria

[mechanism]

C + M <==> MC : Keq1 dissociation
P + M <==> MP : Keq2 dissociation


[concentrations]; micromolar

C = 2000
P = 18

[constants]; micromolar

Keq1 = 0.00789
Keq2 = 0.0001 ?

[responses]

MP = 0.0028

[data]

variable M
offset = auto
set MgGTPCobW

[set:MgGTPCobW]

0	0
4	0.007629
8	0.015819
12	0.02097
16	0.025836
20	0.029532
24	0.031528
34	0.03799

[end]
```

**Dynafit script to estimate extinction coefficient for Fe^II^Tar_2_ complex at 720 nm**

```
[model]

2 Tar monomers bind 1 Fe(II) ion.
Estimate extinction coefficient at 720 nm.

[components]

; C = Tar (competitor)
; M = Fe(II) (metal)

[task]

task = fit
data = equilibria

[mechanism]

C + C + M <==> MC2 : Keq1 dissociation

[concentrations]; molar

C = 16e-6

[constants]; molar

Keq1 = 2.51e-14

[responses]

MC2 = 19000 ?

[data]
variable M
offset = auto
set 720

[set:720]

0		0
0.000002	0.042471
0.000004	0.083526
0.000006	0.126292
0.000008	0.152762
0.00001	0.152132
0.000012	0.152632
0.000016	0.149657
0.00002	0.149805
```

[end]

**Dynafit script to describe competition between Tar and Mg^II^GTP-CobW for Fe^II^**

```
[model]
2 Tar monomers bind 1 Fe(II) ion.
Mg(II)GTP-CobW binds 1 Fe(II) per monomer.

[components]
; C = TAR (competitor)
; P = CobW (protein)
; M = Co(II) (metal)

[task]
task = fit
data = equilibria

[mechanism]
C + C + M <==> MC2 : Keq1 dissociation
P + M <==> MP : Keq2 dissociation

[concentrations]; molar
C = 16e-6
P = 50e-6

[constants]; molar
Keq1 = 2.512e-14
Keq2 = 0.000001 ?

[responses]
MC2 = 19560

[data]
variable M
offset = auto
set MgGTPCobW

[set:MgGTPCobW]
0		0
0.000002	0.036798
0.000004	0.077242
0.000006	0.120745
0.000008	0.141377
0.00001	0.146101
0.000012	0.148976
0.000016	0.149556
0.00002	0.150987
0.00003	0.152325
0.00004	0.153229
0.00006	0.152552
0.00008	0.151422

[end]
```

**Dynafit script to determine extinction coefficient for Ni^II^Tar_2_ complex at 535 nm**

```
[model]

2 Tar monomers bind 1 Ni(II) ion.
Fit extinction coefficient at 535 nm.

[components]
; C = Tar (competitor)
; M = Ni(II) (metal)

[task]
task = fit
data = equilibria

[mechanism]
C + C + M <==> MC2 : Keq1 dissociation


[concentrations]; molar
C = 34e-6

[constants]; molar
Keq1 = 2.3e-16

[responses]
MC2 = 35000 ?


[data]
variable M
offset = auto
set 535

[set:535]

0		0
0.0000033	0.1257
0.0000066	0.248
0.0000099	0.3727
0.0000132	0.4883
0.0000165	0.6058
0.0000198	0.6472
0.0000231	0.6644
0.0000264	0.6636
0.0000297	0.6622
0.000033	0.6607
```

[end]

**Dynafit script to describe competition between Tar and Mg^II^GTP-CobW for Ni^II^**

```
[model]
2 Tar monomers bind 1 Ni(II) ion. Mg(II)GTP-CobW binds 1 Ni(II) per monomer.

[components]
; C = TAR (competitor)
; P = CobW (protein)
; M = Co(II) (metal)

[task]
task = fit
data = equilibria

[mechanism]
C + C + M <==> MC2 : Keq1 dissociation
P + M <==> MP : Keq2 dissociation

[concentrations]; molar
C = 20e-6
P = 30e-6

[constants]; molar
Keq1 = 2.3e-16
Keq2 = 0.000001 ?

[responses]
MC2 = 38000 ?

[data]
variable M
offset = auto
set MgGTPCobW

[set:MgGTPCobW]

0		0
0.000003	0.103169
0.000006	0.198488
0.000009	0.253271
0.000012	0.278219
0.000015	0.299028
0.000018	0.308315
0.000022	0.311758
0.000026	0.325144
0.00003	0.332409
0.000034	0.336594
0.000038	0.350309
0.000042	0.355477
```

[end]

**Dynafit script to describe competition between Bca and Mg^II^GTP-CobW for Cu^I^**

```
[model]

2 Bca monomers bind 1 Cu(I) ion. Mg(II)GTP-CobW binds 1 Cu(I) per monomer.

[components]

; C = Bca (competitor)
; P = CobW (protein)
; M = Cu(I) (metal)

[task]
task = fit
data = equilibria

[mechanism]
C + C + M <==> MC2 : Keq1 dissociation
P + M <==> MP : Keq2 dissociation


[concentrations]; molar
C = 1000e-6
P = 20e-6

[constants]; molar
Keq1= 6.3e-18
Keq2 = 0.0000000000000001 ?

[responses]

MC2 = 7900

[data]
variable M
offset = auto
set MgGTPCobW

[set:MgGTPCobW]

0		0
0.00000336	0.013419
0.00000672	0.028628
0.00001008	0.045342
0.00001344	0.060609
0.0000168	0.077346
0.0000224	0.10741
0.000028	0.140089
0.0000336	0.171832

[end]
```
